# RNA-ligand complexes and the attenuation of neutral confinement in the evolution of RNA secondary structures

**DOI:** 10.64898/2025.12.19.695547

**Authors:** Antonio Loreto, Edgardo Ugalde, Carlos Espinosa-Soto

## Abstract

RNA molecules with identical nucleotide sequence can adopt different structures. Mutations can alter their properties; for example, some mutations increase the stability of a functionally relevant structure at the expense of other structures’ stability. Interestingly, the structural diversity that a sequence produces is correlated to the number of structures that it can access upon mutation. Thus, enhancing a structure’s stability can lead to neutral confinement, an evolutionary dead-end in which mutational access to novel structures is increasingly difficult. If structure is critical to biological function, how do RNA molecules escape neutral confinement? We developed a model in which an RNA molecule’s function depends on binding to a ligand and we applied it to study sequences that fold according to RNA biophysics, also simulating their evolution. Our analyses and simulations identified effects that decrease the selective advantage of augmenting a structure’s stability. By disfavouring evolution of highly stable structures and favouring the accumulation of genetic variation, these effects hinder neutral confinement. The most important effect stems from the sequestration of high affinity structures in RNA-ligand complexes and their replenishment through thermal fluctuations. In this perspective, a common scenario may help to explain how RNA evolution avoids coming to a halt.

## Introduction

After presumably playing a leading role in the early stages of evolution of life on Earth [1], RNA molecules have diversified and acquired a wide range of cellular functions, some of them as fundamental as gene activity regulation, protein synthesis, gene expression, cell differentiation, response to infectious agents and many others [2]. Since structure is critical to define a molecule’s binding partners, many of RNA’s functions depend substantially on the molecule’s structure [3–9].

Recent research supports that a cell’s multiple copies of the same RNA sequence do not fold into the same molecular structure [3; 9–13]. The explanation is simple: one region of an RNA molecule may be able to bind, through Watson-Crick base-pairing, two or more distinct regions of the same molecule. Therefore, RNA molecules with exactly the same nucleotide sequence may adopt different structures. In thermodynamic equilibrium, the probability of finding a specific structure follows the Boltzmann distribution, increasing with the structure’s stability [12–15]. Some such structural variants may not be able to perform the same functions as, for example, the sequence’s minimum free-energy structure (MFES), which is the most stable and abundant structural configuration. Some may even be able to perform other functions that the molecules that adopt the MFES cannot perform [3; 9; 16].

One may be tempted to consider the ability of an RNA sequence to produce structural variation as an evolutionary asset; after all, secondary functions performed by low frequency structures could be the seed for the evolution of new biochemical abilities, as some have suggested for proteins [17]. However, an important and influential computational study on the evolution of RNA structures not only opposes this idea but also supports that such structural variation is indeed an evolutionary handicap [18]. In that study, Ancel and Fontana simulated the evolution of RNA considering a sequence’s *plastic repertoire*, the set of structures that a sequence can fold into. They found that populations often failed to evolve the optimum structure. The reason stems from one of the authors’ important observations concerning the plastic repertoire of a sequence *s*_*i*_ and its *mutational repertoire*, which comprises all the MFES in the set of sequences that differ from *s*_*i*_ by a single nucleotide substitution. They found that structures in the plastic repertoire of a sequence *s*_*i*_ often also appear in *s*_*i*_’s mutational repertoire and that the number of distinct structures in the two repertoires are positively associated. Ancel and Fontana refer to this association as *plastogenetic congruence*, as it shows that plastic effects, resulting from non-genetic perturbations like thermal fluctuations, are aligned with genetic effects. Since then, multiple studies have supplemented statistical evidence for such plastogenetic congruence in RNA molecules [14; 19–23].

Ancel and Fontana assumed, correctly in our opinion, that fitness depends on each structure’s ability to perform a function and on the amount of molecules that adopt such structure. Thus, in their model, a structure’s contribution to fitness reflects how similar is that structure to a pre-defined optimal structure and its expected abundance, according to the Boltzmann distribution [18]. In these circumstances, mutation can increase fitness either by finding new structures more similar to the optimum or by increasing the thermodynamic stability of the better structures in the plastic repertoire. If, during their evolution, sequences encounter more often the latter kind of mutations, fitness would increase but it would do so at the expense of other structures in the plastic repertoire. After all, increasing the relative abundance of one structure implies diminishing that of others, eroding the plastic repertoire’s structural diversity. Because of plastogenetic congruence, this comes at the cost of lower chances to access new better structures through mutation. Thus, selection leads to an evolutionary cul-de-sac in which a population is confined to sequences that yield the same MFES with a high thermodynamic stability and that have little mutational access to further improvement. Ancel and Fontana called this situation *neutral confinement* [15; 18; 24].

Given that RNA sequences yield large, diverse ensembles of structures and that RNA structure is typically crucial to a molecule’s function, we wonder how RNA molecules have avoided the evolutionary dead-end alley that neutral confinement implies. Ancel and Fontana suggested that recombination of distinct RNA molecules could be a way out of neutral confinement [15; 18]. Indeed, recombination seems to be important in the evolution of genomes of RNA viruses [25; 26] and recombination of different RNA molecules could have been specially important in early evolution of life [27]. However, we consider unlikely that recombination suffices to solve the apparent paradox. Here we put forward an additional answer.

Neutral confinement occurs in the Ancel-Fontana model because there is a strong selective incentive to increase the stability of the RNA secondary structures associated to enhanced functional efficiency [18]. Conditions that abate such an incentive may make neutral confinement wane. To search for those conditions we take into account that macromolecules do not exert their function in isolation; rather, they usually need to bind other molecules to play a role useful to the organism.We assumed that the function of an RNA molecule depends on binding to a ligand that, for simplicity, we consider that maintains a constant concentration. A cell’s free RNA molecules with identical sequence may fold into any of several possible structures that differ in their ability to bind the ligand. We argue that structures that better fit the ligand would form RNA-ligand complexes more easily. Remaining free RNA molecules, subject to thermal fluctuations, would convert randomly from one of the sequence’s compatible structures to another, replenishing the pool of RNA molecules that fold into structures best suitable for ligand binding. We hypothesized that, in this scenario, increasing a high affinity structure’s stability would imply only a marginal evolutionary advantage: Consider the case in which structures with a high affinity for the ligand are not very stable. Even then, the sequestration of such structures in RNA-ligand complexes would impel other molecules to build up the stock of high affinity configurations. If this were the case, we may observe a significant attenuation of neutral confinement.

To evaluate the validity of our hypothesis, we developed a differential equations model that takes into account the structural diversity that an RNA sequence can produce, the thermal fluctuation-driven transitions from one structure in the plastic repertoire to another, and the dynamics of binding and unbinding of RNA and ligand molecules. We focused on secondary structures because this level of RNA structure description determines most of the molecule’s stability [15; 28], it provides the basic elements upon which tertiary structure assembles [29; 30] and because we have access to computationally efficient tools and algorithms that, grounded in thermodynamics and RNA biophysics, allow us to predict a sequence’s MFES and plastic repertoires [14; 31; 32].

We studied first hypothetical simplified RNA molecules that only produce two structures in their plastic repertoire. We also studied RNA sequences that fold into specific biological or random structures as their MFES and in which only the rules of RNA folding define the properties of their plastic and mutational repertoires. Congruously with our hypothesis, when we consider explicitly ligand binding, fitness still increases monotonically with the thermodynamic stability of the optimal structure. However, a consistent increase in such stability results in diminishing fitness returns: as thermodynamic stability increases, the selective incentive to increase further thermodynamic stability fades. In addition, we identified conditions that enhance this effect. We also evaluated whether decreasing the fitness benefit that an increased RNA stability confers was sufficient to attenuate neutral confinement. To this end, we analysed evolutionary simulations in which selection favors sequences that are able to yield the greater equilibrium concentrations of RNA-ligand complex. In our simulations, evolution occurs in a stabilizing selection regime in which a population starts with the optimal phenotype and selection favors genetic variants that produce that phenotype more easily. The reason is that we considered that avoiding phenotypic evolutionary transitions would allow easier observation and analysis of neutral confinement. Specifically, evolution in our set-up initiates with RNA sequences that yield as their MFES a structure with maximum affinity for the ligand, albeit perhaps not with a very high thermodynamic stability. We found that mutational robustness and thermodynamic stability are evidently lower in populations in which we consider explicitly ligand binding. In contrast, genetic variation within a population and the population’s access to new structures through mutation and thermal fluctuations are greater in those populations. Therefore, our results support one scenario, that we expect to be widely common in nature, in which evolution of RNA secondary structures avoids the harshest effects of neutral confinement.

## Methods

### Sampling of sequences

We analyzed RNA sequences that fold into specific RNA secondary structures as their minimum free energy structures (MFES). We focused on five biologically relevant RNA structures that we obtained from the RNAcentral database [33] and ten additional structures that we identified as the MFES of random nucleotide sequences (Supplementary Tables ST1,2). The biological structures that we study include those of ribozymes, small nucleolar RNA, the transcriptional regulator DsrA and a tRNA. They are coded in mammalian, bacterial or viral genomes. We used the inverse-fold algorithm [34] in the ViennaRNA library [32] to sample randomly sequences that fold into one of these 15 structures as their MFES. For each structure we sampled 10^4^ sequences for our analyses of individual sequences and 200 more to use as initial sequences in our evolutionary simulations. We show in the section 4 of the Supplementary Text that using a different sampling method still yields qualitatively the same results.

Unless noted, results for all the 15 structures are very similar. Therefore, we present in the main text only results for sequences that have the structure of the CPEB3 ribozyme as their MFES. We report results for all other structures in the supplementary material.

### Access to new phenotypes and robustness

To study the ensemble of secondary structures that a sequence *s*_*i*_ yields, we restricted ourselves to the analysis of the set of structures with a free energy in the interval of width *D* kcal mol^−1^ above the sequence’s MFE, as in previous research [18; 19; 35; 36]. We refer to such a set for *s*_*i*_ as the sequence’s *plastic repertoire* Φ_*i*_. We obtained these alternative structures using the ViennaRNA library [32], that applies Wuchty et al’s algorithm [14] for this purpose. Unless noted, *D* = 2 kcal mol^−1^, which is approximately equal to 3.2*kT* at the default temperature of 37 ^*◦*^C. We picked this value of *D* as a compromise between fast computation and finding a considerable number of the structures with non-negligible stability (Supplementary Table ST3).

The *mutational repertoire* Ξ_*i*_ of an *L*-nucleotide sequence *s*_*i*_ is the set of structures that are accessible to that sequence through single mutations. We obtained it by assaying separately the 3*L* possible nucleotide substitutions and reviewing each mutant sequence’s MFES. To evaluate the mutational access to a specific structure, we counted the number of single mutations that lead to a sequence with that structure as its MFES. A sequence may produce two or more structures with the same MFES. In such a situation, we added all the MFES to its mutational repertoire. In this case, we increased the mutational access to each of these structures by one over the number of MFES in that mutant sequence.

We evaluated a sequence’s mutational robustness by assessing the fraction of all possible single mutations that produce the same structure as its MFES. We estimated the partition function *Z*_*i*_ associated to a sequence *s*_*i*_ as the sum of the Boltzmann factors of the structures in the plastic repertoire. Hence, we obtained the Boltzmann probability 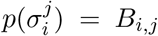 of the plastic repertoire’s *j*-th structure 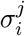 dividing its Boltzmann factor by *Z*_*i*_. We used the Boltzmann probability of a sequence’s MFES as an indicator of its thermodynamic stability.

We assessed a sequence’s intersection between its mutational and plastic repertoires, Ξ_*i*_ and Φ_*i*_, using the overlap coefficient, that equals the cardinality of the intersection of two sets divided by the cardinality of the smaller set. We also used the Pearson correlation coefficient to evaluate the association between, on the one hand, mutational access to the structures in a sequence’s mutational repertoire and, on the other hand, the thermodynamic stability of the structures in the sequence’s plastic repertoire.

We evaluated whether the overlap, or the correlation, between Ξ_*i*_ and Φ_*i*_ were greater than as expected by chance. To do so, we performed paired analyses in which we compared, for each sequence *s*_*i*_ in our sample, the overlap, or correlation, between Ξ_*i*_ and Φ_*i*_ to the one between *s*_*i*_’s mutational repertoire and the plastic repertoire of another sequence *s*_*j*_ picked at random among those sequences that yield the same MFES as *s*_*i*_.

### Ancel-Fontana model

Ancel and Fontana considered that the fitness *ω*_*i*_ associated to a nucleotide sequence *s*_*i*_ equaled the sum of the contributions of the structures in its plastic repertoire Φ_*i*_, i.e. the structures that such sequence could adopt [18]. The contribution of a structure 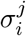 to fitness depended both on its Boltzmann probability 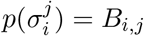 which is associated to its expected abundance in a cell, and on its resemblance 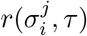 to a pre-defined optimal structure *τ*. More specifically,

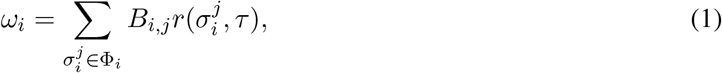

where

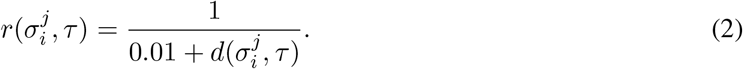

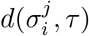 measures the structural distance between structure 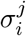 and *τ*. Ancel and Fontana assessed the dissimilarity between two structures using the Hamming distance between the parenthetical notation for RNA secondary structures [18]. Here we instead assess how similar is the arrangement of loops and stacks in the two structures using the tree-edit distance [37], as implemented in the ViennaRNA library [32], but normalized with respect to the sequence’s length.

### RNA-ligand model

In this model we assume the existence of a molecular ligand with a constant concentration ℒ_*i,T*_ in cell *i*. The cell’s fitness depends on the equilibrium concentration of the complex that results from the binding of the cellular copies of an RNA molecule to such ligand. For simplicity and for the purpose of comparison with previous work [18], we consider that an RNA molecule’s ability to bind the ligand only depends on its secondary structure. Thus we assume an optimal structure *τ* with maximum affinity for the ligand. An RNA sequence *s*_*i*_ may adopt any structure 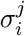 in its plastic repertoire Φ_*i*_, a set with cardinality *M*_*i*_ = |Φ_*i*_|. We consider that ℛ_*i,T*_ , the total amount of copies of the RNA molecule in the cell, remains constant. For simplicity, we also assume that the population of free RNA molecules immediately attains thermodynamic equilibrium, in which the abundance of any such structural configuration will depend on the amount of free RNA molecules and the structure’s Boltzmann probability 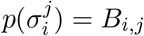. This assumption is irrelevant when the whole system is at equilibrium. Namely, under equilibrium conditions, this model is indistinguishable from a more complicated version that explicitly includes transition rates between the different structural configurations in a sequence’s plastic repertoire (section 1 of the Supplementary Text).

The dynamics of the system in a cell *i* with RNA sequence *s*_*i*_ are described by

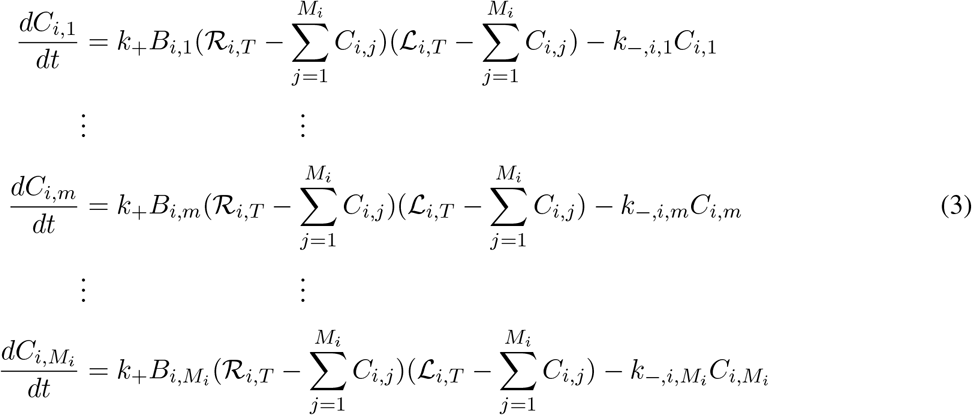

where *C*_*i,m*_ denotes the amount of complex of structure 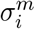 and ligand, 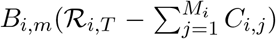 is the amount of free RNA with structure 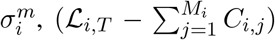 is the amount of non-bound ligand. *k*_+_ is the association constant that we consider the same for all alternative structures. In this model, an RNA molecule’s structure only affects the dissociation rate of their complex with the ligand. This assumption is pertinent not only because it allows a simpler analysis; despite noteworthy exceptions, related molecules that bind the same ligand frequently differ much more on their dissociation rates than on their association rates [38–40]. In the model, the dissociation constant *k*_−,*i,j*_ for complex *C*_*i,j*_ increases with the structural distance between structure 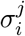 and the optimal structure *τ*. Specifically, we assumed that *k*_−*i,j*_ increased by a constant factor for each structural step away from the optimum structure *τ*: according to

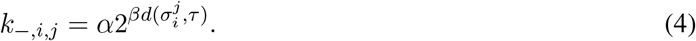

Equations in system (3) are quadratic on several variables. Although there is no general method to solve such systems, we identified properties that allowed us to solve this system exactly (section 2 in Supplementary Text). Moreover, we used previous results on rank-one perturbations of diagonal matrices [41] to show that the system (3) has a unique stable equilibrium point under physically meaningful conditions (section 2 in Supplementary Text). Unless noted, parameter values are ℛ_*i,T*_ = 20 nM, *k*_+_ = 0.001 nM^−1^s^−1^, *α* = 0.001s^−1^, *β* = 0.5.

### Evolutionary simulations

We started a population with *N* identical copies of a nucleotide sequence picked at random among those with the ability to fold into a target secondary structure *τ* as their MFES. Then, we submitted the population to cycles, i.e. generations, of mutation and selection.

Regarding mutation, for each sequence of length *L* we obtained an integer 0 ≤ *u* ≤ *L* from a binomial distribution Bin(*L, µ*), where *µ* is the per-nucleotide mutation rate. Then we picked at random *u* nucleotides from the sequence and replaced them at random. We obtained a sequence’s fitness depending on the model that we followed: Ancel-Fontana, or RNA-ligand, as described above. In all cases, structures more similar to *τ* conferred a greater selective advantage; that is, the populations evolved under stabilizing selection. Next, we applied fitness proportional selection to sample with replacement *N* individuals that constitute the next generation. We repeated these steps for *G* generations.

We analyzed the simulation results using a two-way mixed ANOVA for each of the independent variables that we assessed. We used the optimal structure and scenario as factors, with repeated measures for populations evolving from the same ancestral sequence under different scenarios. For post-hoc tests, we corrected for multiple hypothesis testing using false discovery rate [42]. Our default parameter values are: *N* = 10^3^, *µ* = 5 *×* 10^−5^, *G* = 3 *×* 10^4^.

## Results

### Sequestration of molecules with optimal structures in the RNA-ligand model

To test the validity of our RNA-ligand scenario, we first studied hypothetical RNA molecules that we assume only fold into two structures: an optimal structure *τ* and a non-optimal structure *υ* at a fixed structural distance *d* from *τ*. Because we expected that the behavior of the system varied with the amount of ligand respective to the amount of RNA, we assayed three different concentrations of ligand molecule. We also studied these hypothetical RNA molecules under the perspective of the Ancel-Fontana model. For the Ancel-Fontana model [18] we measured fitness as the average contribution to fitness of each structure, weighted by each structure’s thermodynamic stability (see details in Methods). In the RNA-ligand model, we quantified fitness as the total amount of RNA-ligand complex at thermodynamic equilibrium. In all scenarios, the theoretical maximum fitness value corresponds to a sequence with a plastic repertoire that only contains the optimal structure. We use such maximum fitness as a normalization constant thus allowing us to compare relative fitness fairly among different scenarios.

Fig. 1 shows fitness (blue line) as a function of the thermodynamic stability of the optimal structure. The figure shows that fitness increases linearly with the stability of the optimal structure *τ* in the Ancel-Fontana model. In the RNA-ligand model, fitness also increases monotonically with *τ* ‘s stability. However, in this case fitness is a concave function of *B*_*i*_: as the Boltzmann probability increases, further increase leads to a slighter fitness gain. As we expected, concavity, relative fitness and, consequently, selective proximity to the optimum, are the greatest when the ligand concentration is the highest; in this case the excess molecules of ligand are able to sequester a greater amount of RNA molecules with the optimal structure. For the examples in Fig. 1 we maintained a fixed, relatively low, structural distance *d* = 8 between the optimal structure *τ* and the suboptimal structure. Fig. SF1 in Supplementary Figures shows that the effect is qualitatively the same even when we fix a much higher value for *d*.

**Fig. 1.**
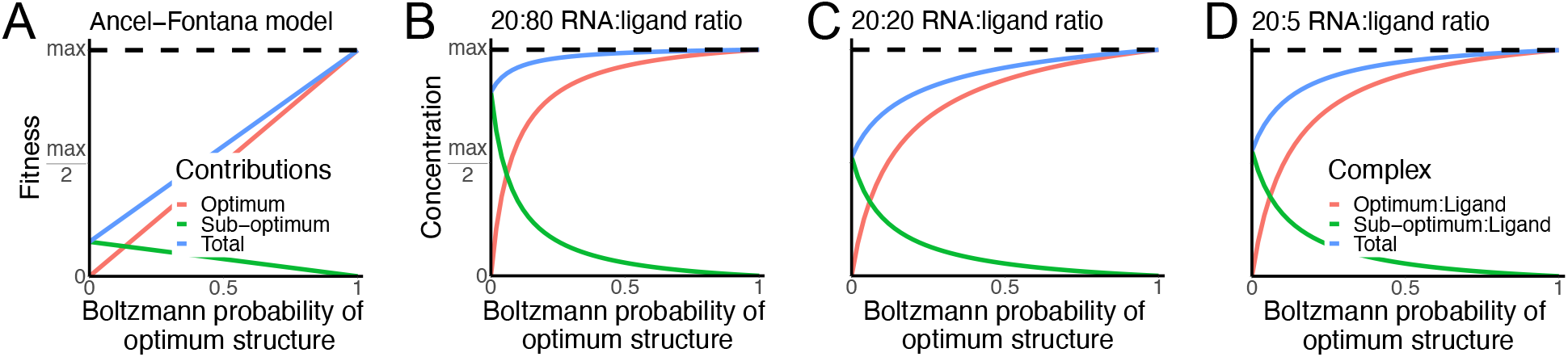
Fitness as a function of thermodynamic stability of a sequence’s MFES. We assume that all sequences have two different structures in their plastic repertoire: an optimal structure *τ* and a second structure *υ* at a fixed structural distance *d* = 8 from *τ*. Green and red lines indicate *υ*’s and *τ* ‘s contribution to fitness, respectively. The blue line refers to the sequence’s total fitness. The horizontal dashed line indicates the maximum fitness value in each scenario.

### Fitness as a concave function of thermodynamic stability in the RNA-ligand model

For Fig. 1 we applied our RNA-ligand model to simplified hypothetical RNA molecules that, non-realistically, always have the same two structures in their plastic repertoires. Now we do the same but for RNA sequences, first, that can fold into complex and diverse ensembles of structures defined by RNA biophysics and, second, in which we have described the association between variation produced by mutation and thermal fluctuations, as shown in section 3 of the Supplementary Text. For comparison, we also study the influence of thermodynamic stability on fitness in the Ancel-Fontana model.

Notwithstanding the intricacies of RNA folding into ensembles of secondary structures, our results are very similar to the ones that we presented for hypothetical simplified RNA molecules. Fig. 2 shows our results for sequences that fold into the structure of the CPEB3 ribozyme as their MFES. The corresponding data for the remaining structures in our study, biological or random, appear in Supplementary Figures SF2,3, respectively. We confirmed that the relationship between the Boltzmann probability of the MFES and fitness is near-linear in the Ancel-Fontana model: mutations that increase thermodynamic stability to the same extent would always produce the same increment in fitness. In contrast, in our RNA-ligand model and according to our expectations, fitness is a concave down function of thermodynamic stability. Hence, adding to a sequence a series of mutations that increase more and more the Boltzmann probability of a structure with a high affinity for the ligand would result in diminishing fitness returns.

**Fig. 2.**
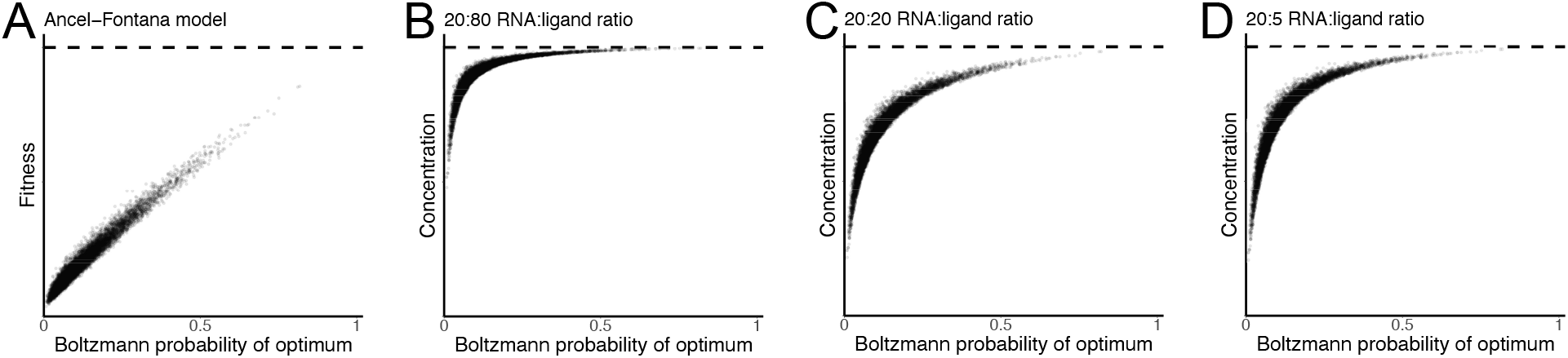
Fitness as a function of thermodynamic stability in sequences that fold into the structure of the CPEB3 ribozyme as their MFES. We assess thermodynamic stability as a sequence’s Boltzmann probability for the structure of the CPEB3 ribozyme. The horizontal dashed line indicates the maximum fitness value in each scenario.

Our hypothesis states that sequestration of optimal and near-optimal structures in RNA-ligand complexes creates a net conversion of unbound molecules to high-affinity structures, replenishing their stock. In this scenario, even with a lower stability, the sequestration-driven accumulation of high-affinity structures would increase complex formation. Thus, we expect that a high concentration of ligand would increase relative fitness and hence its selective proximity to the optimum. It would also intensify fitness concavity as a function of thermodynamic stability. After all, the sequestration of near-optimal of structures would be stronger when there are more ligand molecules. Fig. 2 and Supplementary Figures SF2,3 show that relative fitness and its concavity are indeed greatest in the simulations in which we considered the highest ligand concentration. This is more evident in Fig. 3 and Supplementary Figures SF4,5, that show that a sequence’s relative fitness is always the greatest, and hence more proximate to the maximum, in the ligand-RNA model under high ligand concentrations. The figures also show that relative fitness is always the lowest, farther from the maximum value, under the Ancel-Fontana model. However, the story is not straightforward: concavity and relative fitness do not decrease monotonically as the ligand concentration diminishes. The plots of fitness as a function of thermodynamic stability for intermediate and low concentrations of ligand (Fig. 2 and Supplementary Figures SF2 and SF3) seem very similar. However, the pattern is clearer in the direct comparison of relative fitness between these two models: Relative fitness is slightly but always higher for low than for intermediate concentrations of ligand (Fig. 3 and Supplementary Figures SF4,5). In fact, we also observed this pattern when we studied the hypothetical simplified RNA molecules for Fig. 1. All these observations suggest that ligand binding implies an additional effect, stronger at low ligand concentrations, that increases relative fitness.

**Fig. 3.**
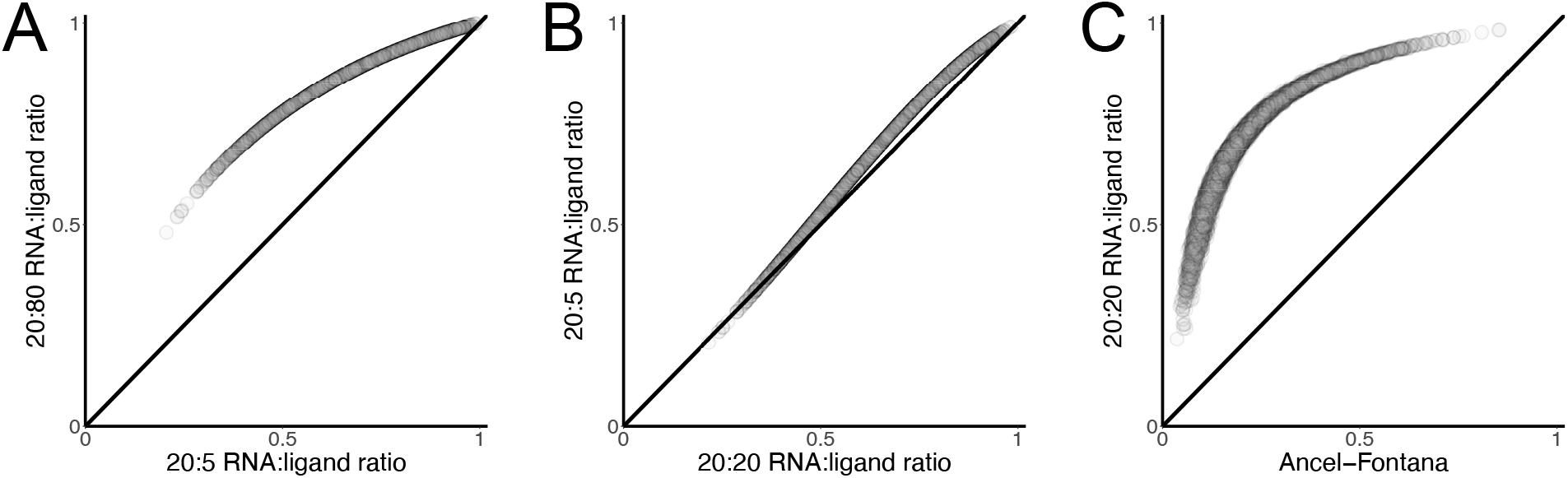
Comparison of relative fitness under different models or ligand concentrations for sequences that fold into the structure of the CPEB3 ribozyme. In all panels, the diagonal indicates the identity line. Points above the diagonal refer to sequences with a higher fitness under the model/ligand concentration that corresponds to the vertical axis than under conditions described in the horizontal axis. Fitness values are normalized with respect to the maximum fitness value for the respective model.

### Additional effects of RNA-ligand binding on relative fitness

One of the additional effects that we expect is that RNA molecules saturate ligand molecules more easily when the latter are scarce, requiring less RNA molecules with a high ligand-affinity structure to approach the maximum amount of RNA-ligand complex. To observe and better study this effect, we developed a new version of our model in which, as a non-realistic and non-natural experiment, we blocked transitions in free RNA molecules from one structure in the plastic repertoire to another, preventing the sequestration-dependent flux towards structures with higher affinity for the ligand. Namely, we consider that the number of RNA molecules that fold into a certain structure *υ*, whether free or ligand-bound, remains constant and proportional to *υ*’s Boltzmann probability. The *m*-th of *M* equations that describe this version of the model is

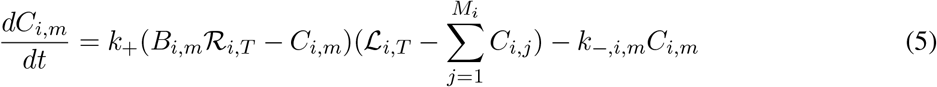

The term (*B*_*i,m*_ℛ_*i,T*_ − *C*_*i,m*_) defines the abundance of free RNA molecules with structure 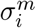. It equals the total amount of RNA molecules with that structure, held constant at a value of *B*_*i,m*_ℛ_*i,T*_ , minus the portion of those molecules that take parte in RNA-ligand complexes. We used Powell’s hybrid method [43] as implemented in the GSL library [44] to assess the complex concentration at equilibrium.

This version of the model shows much more dispersion on fitness values, suggesting that particularities of each sequence *s*_*i*_ and of its plastic repertoire Φ_*i*_ play a more important role in it (Supplementary Figures SF6,7). Supporting that RNA-ligand complexes saturate more easily at low ligand concentrations, we found that, when we prohibit transitions between alternative RNA structures, concavity and relative fitness are greatest in the scenario in which ligand concentration is lowest (Supplementary Figures SF6-9). However, the association between ligand abundance and relative fitness is neither monotonic in this case; relative fitness and, consequently, proximity to the optimum, are slightly greater under high than under intermediate ligand concentrations (Supplementary Figures SF8,9), suggesting that there is a third additional factor that increases relative fitness at high ligand concentrations. The effect is clearer at low Boltzmann probabilities for the optimal structure (Supplementary Figures SF6,7). We hypothesized that this effect occurred because, when ligand is in excess, molecules with a lower affinity structure can also bind the ligand, increasing the concentration of RNA-ligand complex. We reasoned that this effect should be sensitive to the affinity for the ligand of molecules in the plastic repertoire: a very low affinity would hinder it severely. Testing this hypothesis with sequences that fold into large and diverse ensembles of structures following intricate patterns dictated by RNA biophysics would be complicated. We then studied the version of the model in which we block transitions between different structures but now applying it to hypothetical simplified RNA molecules with only two structures in their plastic repertoire. Consistently with our hypothesis, when ligand is in excess, molecules with a lower affinity structure can also bind the ligand, increasing the concentration of RNA-ligand complex. However, if the affinity is too small, ligand abundance is not sufficient to recruit RNA molecules into complexes (Supplementary Figure SF10).

### Attenuation of neutral confinement in evolution under stabilizing selection

We have shown that explicitly considering the binding of RNA molecules to a ligand to form functional complexes reveals fitness as a concave down function of thermodynamic stability in which the relative fitness difference to the optimum is diminished. We designed evolutionary simulations to assess whether weakening the selective incentive to increase thermodynamic stability obstructs the appearance of neutral confinement. We compared populations evolving in either the Ancel-Fontana scenario, under which neutral confinement easily evolves [18], or in the RNA-ligand scenario, with three different levels of ligand abundance. We expected neutral confinement to be strongest in the Ancel-Fontana scenario; within the RNA-ligand scenario, our previous results suggest that neutral confinement would decrease its strength from intermediate to low and high concentrations of ligand. Our simulations occur in a stabilizing selection regime: evolution starts with sequences that already produce the optimum structure as their MFES, although not necessarily with a high thermodynamic stability. We designed our simulations this way to avoid complications in our analysis and to focus on a population’s accumulation of genetic variation and its ability to access different phenotypes. We picked one of the structures in Supplementary Tables ST1 and ST2 as the optimal structure for each of our simulations. In the Ancel-Fontana model, stabilizing selection favors those sequences that yield a stabler optimal structure. In the RNA-ligand models the optimal structure has the highest affinity for the ligand and stabilizing selection favors sequences that produce a higher RNA-ligand complex equilibrium concentration. The results of the statistical analyses of these simulations appear complete in Supplementary Tables ST4-33 and Supplementary Figs. SF11-20.

First we compared the amount of genetic variation that populations hold after 3 *×* 10^4^ generations of evolution under stabilizing selection. We found that the scenario has a significant effect on the amount of genetic variation that populations hold and that this observation is valid regardless the identity of the optimal structure (Supplementary Tables ST4,5). Moreover, as expected, the number of different sequences in the population is, in all cases, significantly lowest in the Ancel-Fontana scenario and significantly highest in the high-ligand RNA-ligand setup (Fig. 4A, Supplementary Table ST6, Supplementary Fig. SF11). We also considered other measures of genetic diversity like heterozygosity, nucleotide diversity [45] and the maximum number of nucleotide differences between any two sequences in the population. They follow the same pattern; however, the results of the post-hoc tests for these other indicators of genetic variation are not as consistent as for the number of different sequences in the population (Supplementary Tables ST7-15 and Supplementary Figs. SF12-14).

**Fig. 4.**
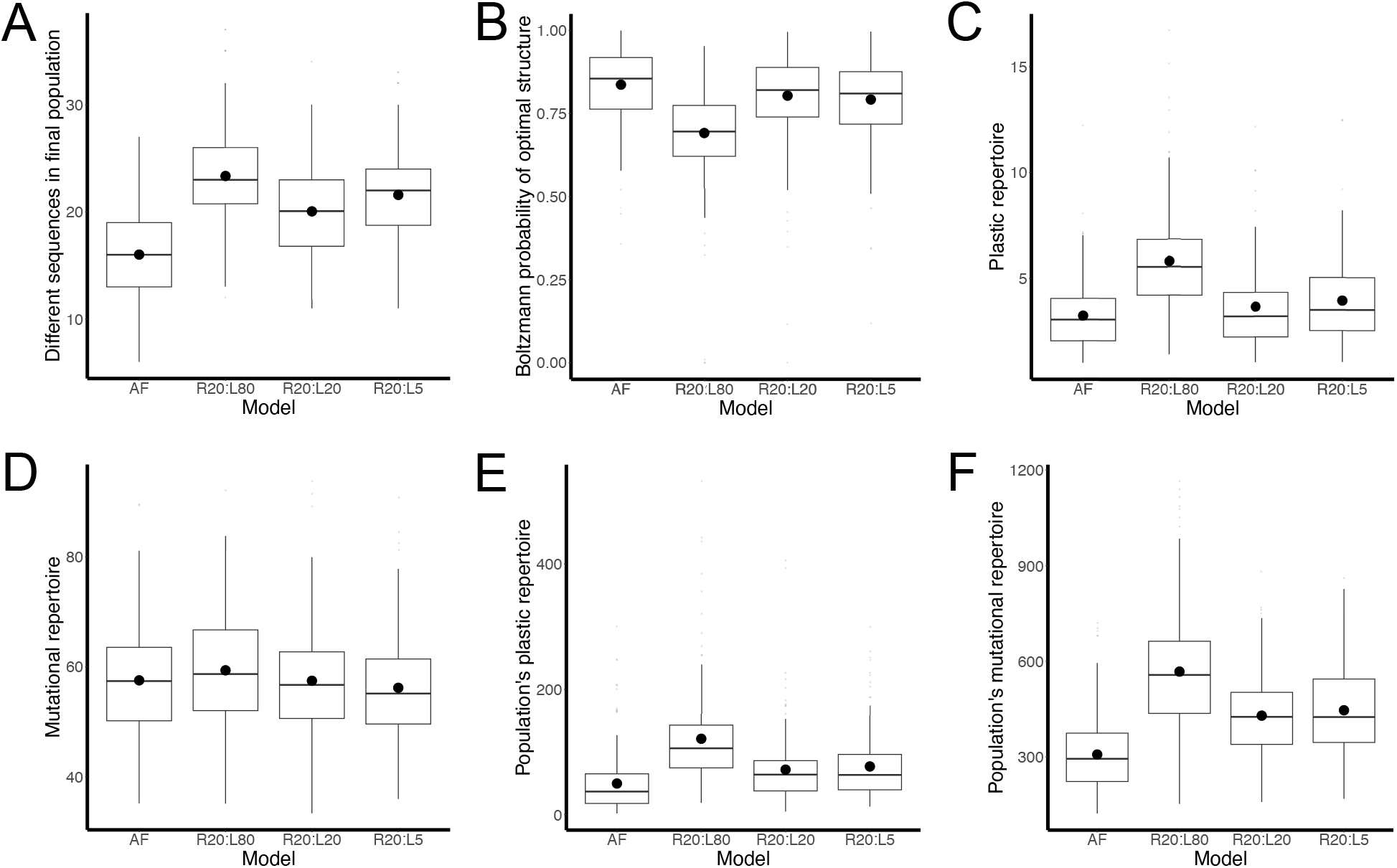
Evolution under stabilizing selection in favor of the CPEB3 ribozyme structure under different scenarios. AF: Ancel-Fontana model; RX:LY: RNA-ligand scenario with X and Y as the constant concentrations of the RNA molecule and the ligand, respectively. A. Number of distinct sequences in the population at the end of evolution. B. Average thermodynamic stability at the end of evolution. C,D. Cardinality of the plastic and mutational repertoires, respectively. E,F. Population’s access to new phenotypes through plasticity and mutation, respectively. We define the population’s plastic, or mutational, repertoire as the union of the plastic, or mutational, repertoires of all the sequences that comprise a population.

As expected, thermodynamic stability (Fig. 4B, Supplementary Tables ST16-18 and Supplementary Fig. SF15) and mutational robustness (Supplementary Fig. SF16, Supplementary Table ST19-21) follow a similar pattern, opposite to that of genetic variation. Despite their similar trend, average differences are much more pronounced in the case of thermodynamic stability. Together, these findings support that the RNA-ligand scenario tends to diminish the concentration of individuals with high robustness genotypes.

In addition, we looked at the access to new secondary structures. Access to new structures through plasticity, as indicated by the size of a sequence’s plastic repertoire, was significantly higher in sequences evolved under the RNA-ligand scenario than in those evolved in the Ancel-Fontana scenario (Fig. 4C, Supplementary Fig. SF17, Supplementary Tables ST22-24). In contrast, a sequence’s average mutational repertoire was not always significantly lower in the Ancel-Fontana than in the RNA-ligand scenario (Fig. 4D, Supplementary Tables ST25-27, Supplementary Fig. SF18). Indeed, for several optimal structures we failed to find a significant effect of the model on the size of the mutational repertoire of individual sequences (Supplementary Table ST26). For example, sequences folding into the structure of the CPEB3 ribozyme that evolved under selection for complex formation have mutational repertoires without significant differences to those of sequences evolved in the Ancel-Fontana scenario (Fig. 4D, Supplementary Table ST27).

While individual sequences evolved under the RNA-ligand scenario do not have easier mutational access to new structures, there may still be differences in the mutational access to new structures at the level of the population. The increased genetic variation in populations that evolved under the RNA-ligand scenario might imply a smaller overlap between the mutational (or plastic) repertoires of different sequences of the population. If this were the case, a more diverse population may have access to a wider range of structural diversity, despite similar access to new structures of individual sequences. To assess this possibility we define a population’s plastic repertoire as the union of all the plastic repertoires of the sequences that comprise a population; it lists all the structures that appear in the plastic repertoire of at least one of the population’s individuals. Analogously, we define the population’s mutational repertoire as the union of the mutational repertoires of all the population’s individuals.

We found that a population’s access to new structures through either mutation or plasticity is clearly greater in populations that evolved under the RNA-ligand scenario (Figs. 4E,F, Supplementary Tables ST28-33 and Supplementary Figs. SF19,20). For example, populations that evolved in the Ancel-Fontana scenario under selection for the structure of the CPEB3 ribozyme could access, on average, 307.9 structures through mutation and 49.8 through plasticity. In contrast, when they evolved in the RNA-ligand scenario under high-ligand conditions, populations could access 568.1 and 121.2 through mutation and plasticity, respectively. Even under conditions of intermediate ligand concentrations, the worst case among the RNA-ligand scenarios, there is much easier access to new structural variation (429.9 through mutation and 71.96 through plasticity) than in populations evolved in the Ancel-Fontana setup (Figs. 4E,F; Supplementary Tables ST30,33). Populations that evolved under selection for any of the structures that we studied clearly follow the same pattern (Supplementary Tables ST28-33 and Supplementary Figs. SF19,20).

## Discussion

Previous research has established that a consistent increase in the stability of one RNA structure *υ* at the expense of others leads to neutral confinement. A population in neutral confinement becomes locked in a cluster of sequences that produce a highly stable *υ* configuration but with scant mutational access to new, perhaps better, structures [15; 18; 19; 24]. One could say that sequences become victims of their success. The support to the idea of neutral confinement rests on three sound and robust premises: first, RNA structure as a strong determinant of biological function [3–5; 7–9];; second, within-cell conformational diversity of the structures of RNA molecules with the same sequence [3; 9–13]; and third, plastogenetic congruence, a positive association between the amount and kind of structural variation produced by thermal fluctuations and mutation [14; 19–23, section 3 of Supplementary Text]. Nevertheless, although all three premises hold, the consequences of neutral confinement are not apparent in nature, as indicated by the diversity of molecular functions that RNA molecules have acquired in all lineages throughout evolution [2]. One way in which RNA molecules may escape neutral confinement is through recombination of different sequences [18; 19; 24]. Ultimately, recombination can merge in one sequence different substructures with independent evolutionary histories. The creation of innovation through recombination in RNA sequences seems to be frequent in RNA viruses [25; 26] and probably was also important during the RNA world [27]. However, to our knowledge, there is no evidence that supports it as a common evolutionary mechanism for RNA structural innovation in modern cellular organisms.

Here we put forward an alternative to avoid neutral confinement in the evolution of RNA. Our alternative does not dispute any of the three important premises that uphold neutral confinement. In our proposal, the function of an RNA molecule depends on specific binding to a ligand to form an RNA-ligand complex and affinity for the ligand depends on the RNA’s structure. Our work identifies three important effects of ligand binding:

- Ligand molecules may function as sinks for RNA molecules with a configuration with higher affinities for the ligand, that they easily entrap into RNA-ligand complexes. As the free RNA molecules wiggle because of thermal fluctuations, they replenish the stock of molecules with a high affinity structure.
- Under low ligand concentrations, most ligand molecules bind high-affinity RNA molecules promptly. Since most ligand molecules are occupied, availability of additional high-affinity RNA molecules produces only marginal increases in the amount of RNA-ligand complex.
- Under high ligand concentrations not only high-affinity molecules bind the ligand; but a significant fraction of RNA molecules with alternative configurations also can bind the abundant free ligand molecules.

Assuming that organismal fitness depends on the amount of RNA-ligand complex, all these factors diminish the fitness difference between organisms with an RNA sequence that produce the same structure but with different Boltzmann probabilities. Thus, populations lack a strong incentive to evolve sequences that produce extraordinarily stable structures but with a meager mutational access to new structures. As our simulations of evolution show, instead of neutral confinement, populations may retain genetic variation within a population, preserve access to new structures through mutation and thermal fluctuations, and maintain the ability to explore genotype space through nearly-neutral mutations.

In our model, the most important way in which ligand-binding weakens neutral confinement is by the sequestration into complexes of high affinity configurations and the consequent restoration of their supply by thermal fluctuations. Outside of the context of RNA evolution, the idea that binding to a ligand can shift the equilibrium distribution of protein conformations is not novel and is well studied [46–48]. For example, *ligand-directed conformational selection* is one of the central ingredients in the Monod-Wyman-Changeux model for allosteric regulation [49–51]. Otherwise, our model hinges on the three same premises for neutral confinement and the reasonable consideration that much of RNA’s function depends on binding to other molecules to form multimeric complexes [2]. Thus, it may be relevant and valid for a wide range of cases.

Our evolutionary simulations show how the RNA-ligand scenario affects a population’s access to new structures through plasticity and to new MFE structures through mutation. Regarding mutations, individual sequences that evolved in the RNA-ligand scenario did not have consistently greater mutational repertoires. However, the whole population’s access to new MFES through mutation was greater when evolution occurred under the RNA-ligand scenario. The reason is that populations in this scenario accrue greater genetic diversity and the mutational repertoires of less similar sequences tend to be more different [52]. Regarding access to new structures through plasticity, individual sequences that evolve in the RNA-ligand scenario typically yield a more diverse plastic repertoire. As our work shows, non-MFE structures may attain significant cellular concentrations if they have a high affinity for a ligand. Therefore, greater individual plastic repertoires may be relevant for the discovery of new functional structures. Moreover, the whole population also has easier access to new structures through plasticity in the RNA-ligand scenario. This is not surprising since, in that scenario, individual plastic repertoires are already greater, the population has more genetic diversity and, as reported previously, a few substitutions can significantly alter a sequence’s plastic repertoire [36].

In this contribution we focused on whether considering ligand-binding explicitly was sufficient to undermine neutral confinement. In our evolutionary simulations, populations evolved in a stabilizing selection regime: the optimal RNA structure was present in the population as a minimum free energy structure from the beginning of our simulations, although perhaps not with a very high thermodynamic stability. Further research is necessary to evaluate how our scenario, and the consequent erosion of neutral confinement, impacts on the evolution of novel structures. Indeed there are interesting observations that suggest that formation of functional RNA-ligand complexes can accelerate evolution of new structures in the face of RNA conformational diversity. First, previous research already shows that a few mutations can change many of the structures in a plastic repertoire [36]. Hence, exploration of those structures accessible through thermal fluctuations may allow faster discovery of structures with enhanced functionality, albeit most often with a low thermodynamic stability. Importantly, our work supports that a structure with high propensity to assemble complexes but with a low thermodynamic stability can confer a non-negligible fitness benefit, facilitating the action of selection. In this case, the few molecules with the new functional structure would form a complex promptly, promoting the conversion of other molecules into the high affinity configuration. Moreover, once a population is enriched with sequences with the high affinity structure in their plastic repertoire, plastogenetic congruence may favor the appearance of sequences that produce the beneficial structure with higher stability or even as their MFES. The existence of clusters of similar sequences that can access the same structure through mutations [53] could facilitate further these transitions. Validation of this prospect by future research would strengthen the case of plasticity in RNA secondary structures as an evolutionary asset instead of an obstacle, as in other kinds of biological systems, including protein structure [17], gene expression patterns [54–56] and morphology [57; 58]. Whether mechanisms analogous to the one that we describe for RNA secondary structures could apply to protein folding or other kinds of traits remains as an open possibility.

## Supplementary material

Supplementary Text, Tables and Figures are available online.

## Acknowledgements

We thank the most valuable technical assistance from JC Sánchez-Leaños and useful comments by L.F. Meraz-Segura, A.G. Cruz-Moreno and H.J. Villasana.

## Authors’ contributions

AL and CE-S developed code and performed research. CE-S, AL, and EU developed the differential equation model, analyzed the data and designed research. CE-S wrote the paper. All authors revised and approved the paper.

