## Supplementary Text for "RNA-ligand complexes and the attenuation of neutral confinement in the evolution of RNA secondary structures"

**3 Plastogenetic congruence** **9**

**4 Analyses of sequences sampled with a random mutational walk subsequent to inverse folding** **14**

**References** **14**

### 1 Model with finite transition rates between alternative RNA secondary structures

In the main text we describe a system in which free RNA molecules attain immediately their equilibrium distribution, as predicted by their Boltzmann probability. Here we relax such assumption, considering a model in which unbound RNA molecules can change their secondary structures at finite rates, as described by the dashed arrows in the illustration of Fig. S1. As in the model in the main text, RNA molecules can also bind reversibly to free ligand molecules to form an RNA-ligand complex, as schematized by solid arrows in Fig. S1. Conversion of unbound RNA molecules from one of its alternative configurations to another occurs with rate constants described in an $M \times M$ square matrix $\mathbf{Z}$. $M$ refers to a sequence's number of distinct alternative RNA secondary structures, that is, the cardinality of the sequence's plastic repertoire $\Phi$. Specifically, an entry $z_{i,j}$ in the matrix $\mathbf{Z}$ represents the rate constant for the transformation of structure $\sigma^j$ into structure $\sigma^i$ (dashed arrows in Fig. S1). Association and dissociation of RNA and ligand molecules (solid arrows in Fig. S1) occurs as in the model described in the main text. Dynamics are described by the system of $2M-1$ equations

$$
\begin{aligned}
\frac{dC_1}{dt} &= k_+ R_1(\mathcal{L}_T - \sum_{l=1}^{M} C_l) - k_{-,1} C_1 \\
&\vdots \\
\frac{dC_i}{dt} &= k_+ R_i(\mathcal{L}_T - \sum_{l=1}^{M} C_l) - k_{-,i} C_i \\
&\vdots \\
\frac{dC_{M-1}}{dt} &= k_+ R_{M-1}(\mathcal{L}_T - \sum_{l=1}^{M} C_l) - k_{-,M-1} C_{M-1} \\
\frac{dC_M}{dt} &= k_+ (\mathcal{R}_T - \sum_{l=1}^{M} C_l - \sum_{l=1}^{M-1} R_l)(\mathcal{L}_T - \sum_{l=1}^{M} C_l) - k_{-,M} C_M \\
\frac{dR_1}{dt} &= k_{-,1} C_1 - k_+ R_1(\mathcal{L}_T - \sum_{l=1}^{M} C_l) + \sum_{j=2}^{M-1} z_{1,j} R_j + z_{1,M}(\mathcal{R}_T - \sum_{l=1}^{M} C_l - \sum_{l=1}^{M-1} R_l) - R_1 \sum_{j=2}^{M} z_{j,1} \\
&\vdots \\
\frac{dR_i}{dt} &= k_{-,i} C_i - k_+ R_i(\mathcal{L}_T - \sum_{l=1}^{M} C_l) + \sum_{j=1;j\neq i}^{M-1} z_{i,j} R_j + z_{i,M}(\mathcal{R}_T - \sum_{l=1}^{M} C_l - \sum_{l=1}^{M-1} R_l) - R_i \sum_{j=1;j\neq i}^{M} z_{j,i} \\
&\vdots \\
\frac{dR_{M-1}}{dt} &= k_{-,M-1} C_{M-1} - k_+ R_{M-1}(\mathcal{L}_T - \sum_{l=1}^{M} C_l) + \sum_{j=1}^{M-2} z_{M-1,j} R_j + z_{M-1,M}(\mathcal{R}_T - \sum_{l=1}^{M} C_l - \sum_{l=1}^{M-1} R_l) - R_{M-1} \sum_{j=1;j\neq M-1}^{M} z_{j,M-1}
\end{aligned}
\tag{1}
$$

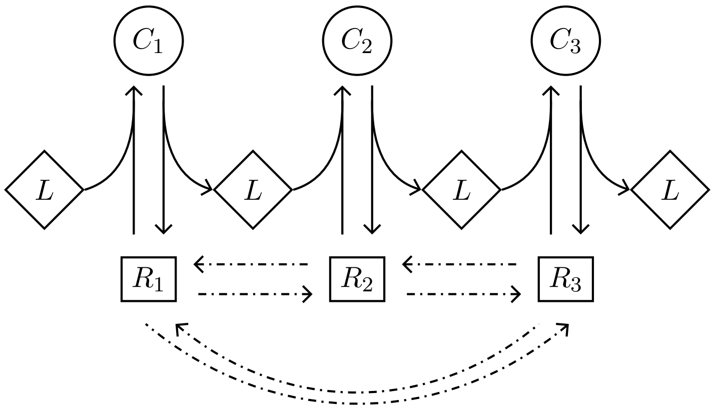

**Fig. S1. Illustration of reactions in the system.** This example considers an RNA sequence that can fold into three different secondary structures, i.e. $M = 3$. Squared nodes ($R_i$) represent free RNA molecules with one of such structures. Diamond nodes ($L$) represent ligand molecules. Circular nodes ($C_i$) stand for RNA-ligand complexes.

Variables and parameters appearing also in the equations in the main text preserve the same meaning. However, here we omit for clarity the subindex that identifies the cell or organism that bears the RNA sequence. $R_i$ represents the amount of free RNA molecule with structure $\sigma^i$. The colored parts in the system of equations (1) represent association and dissociation of RNA molecules and ligand (solid arrows in Fig. S1). The black parts on the right side of the last $M - 1$ equations correspond to the conversion between alternative configurations of unbound RNA molecules (dashed arrows in Fig. S1).

Solving this system of equations would require knowing the values of all the rate constants in matrix $\mathbf{Z}$. While we can obtain the Boltzmann probabilities for each alternative structure using tools available from the ViennaRNA library [1], these probabilities do not allow us to determine unambiguously the rate constants. However, let's consider that the system is at equilibrium and all $2M - 1$ equations equal zero. That the first $M$ equations equal zero implies that the colored parts in the last $M-1$ equations also equal zero, as they have the same magnitude as the correspondingly colored terms in the first $M$ equations. Therefore, the system's equilibrium requires that terms in black on the right side of each of the bottom $M - 1$ equations are also null. This only occurs when transitions between alternative structures in unbound RNA molecules compensate each other, maintaining constant the amount of unbound molecules with each alternative structure; that is, when the concentrations of alternative configurations of free RNA molecules match their Boltzmann probability. Therefore, this more realistic albeit more complicated system is indistinguishable *at equilibrium* from the simpler system, that we describe in the main text, in which we assume that free RNA molecules attain their Boltzmann probabilities instantaneously.

### 2 Analysis of the model in the main text

#### 2.1 Assessment of equilibrium points

In the main text we considered that the rate of change of a complex formed by a ligand molecule and an RNA molecule with structure $\sigma^i$ is given by

$$\frac{dC_i}{dt} = k_+ B_i(\mathcal{R}_T - \sum_{m=1}^{M} C_m)(\mathcal{L}_T - \sum_{m=1}^{M} C_m) - k_{-,i} C_i \tag{2}$$

. All variables and parameters are as defined in the main text. The only notational difference is that here we omit for clarity the subindex that identifies the cell or organism in which dynamics occurs. Expression (2) is valid for any of the $M$ structures in a sequence's plastic repertoire $\Phi$. We may describe the complete system of equations in matrix form, considering the following vectors and matrices:

$$\mathbf{C} = \begin{bmatrix} C_1 \\ \vdots \\ C_i \\ \vdots \\ C_M \end{bmatrix}, \qquad \mathbf{D} = \begin{bmatrix} \frac{k_{-,1}}{k_+B_1} & 0 & \dots & 0 \\ 0 & \frac{k_{-,2}}{k_+B_2} & 0 & \vdots \\ \vdots & 0 & \ddots & \vdots \\ 0 & \dots & 0 & \frac{k_{-,M}}{k_+B_M} \end{bmatrix},$$

$$\mathbf{Q} = \begin{bmatrix} \frac{1}{k_+B_1} & 0 & \dots & 0 \\ 0 & \frac{1}{k_+B_2} & 0 & \vdots \\ \vdots & 0 & \ddots & \vdots \\ 0 & \dots & 0 & \frac{1}{k_+B_M} \end{bmatrix}, \qquad \mathbb{1} = \begin{bmatrix} 1 \\ \vdots \\ 1 \\ \vdots \\ 1 \end{bmatrix}$$

Using the superscript $^\dagger$ to refer to the transpose of a matrix, the whole system at equilibrium is described by

$$G(\mathbf{C}) = \mathbf{Q}\frac{d\mathbf{C}}{dt} = (\mathcal{R}_T - \mathbb{1}^\dagger \mathbf{C})(\mathcal{L}_T - \mathbb{1}^\dagger \mathbf{C})\mathbb{1} - \mathbf{D}\mathbf{C} = \mathbf{0} \tag{3}$$

In order to solve this quadratic equation on several variables, we first partition the solution space $\mathbb{R}^M$ into parallel planes

$$\mathbb{R}^M = \bigcup_{\beta \in \mathbb{R}} \left\{ \mathbf{C} \in \mathbb{R}^M : \mathbb{1}^\dagger \mathbf{C} = \beta \right\}$$

Let's assume that there is a solution $\mathbf{C}$ on the plane $E_\beta := \{\mathbf{C} \in \mathbb{R}^M : \mathbb{1}^\dagger \mathbf{C} = \beta\}$ that satisfies eq. (3). We can find such solution solving the lineal system

$$(\mathcal{R}_T - \beta)(\mathcal{L}_T - \beta)\mathbb{1} = \mathbf{DC}.$$

Because $\mathbf{D}$ is diagonal, the solution is simple:

$$C_i = \frac{(\mathcal{R}_T - \beta)(\mathcal{L}_T - \beta)}{(\mathbf{D})_{i,i}} = \frac{k_+ B_i}{k_{-,i}}(\mathcal{R}_T - \beta)(\mathcal{L}_T - \beta) \tag{4}$$

This will be a solution of eq. (3) if it is true that

$$\sum_{i=1}^{M} C_i = \beta = \sum_{i=1}^{M} \frac{k_+ B_i}{k_{-,i}}(\mathcal{R}_T - \beta)(\mathcal{L}_T - \beta) \tag{5}$$

We can easily solve this quadratic equation to obtain a $\beta$ that satisfies eq. (5), and then substitute this value in eq. (4) to find the values of $C_i$ that satisfy the system (3).

#### 2.2 Uniqueness of the physically meaningful solution

That the solution of the system of equations (3) is physically meaningful requires the conditions

$$0 \le \beta < \mathcal{L}_T \qquad ; \qquad 0 \le \beta < \mathcal{R}_T \tag{6}$$

Eq.(5) defines the function

$$\beta \to F(\beta) := \sum_{i=1}^{M} \frac{k_+ B_i}{k_{-,i}}(\mathcal{R}_T - \beta)(\mathcal{L}_T - \beta) \tag{7}$$

Let's say

$$F(\beta) = \sum_{i=1}^{M} f_i(\beta) \tag{8}$$

, with functions $f_i(\beta)$

$$f_i(\beta) = \frac{k_+ B_i}{k_{-,i}}(\mathcal{R}_T - \beta)(\mathcal{L}_T - \beta)$$

That $k_+$, $k_{-,i}$, $B_i$, $\mathcal{R}_T$ and $\mathcal{L}_T$ are positive implies that, for each $f_i(\beta)$,

$$f_i(0) > 0 \quad ; \quad f_i(\mathcal{L}_T) = 0 \quad ; \quad f_i(\mathcal{R}_T) = 0$$

The first and second derivatives of each $f_i(\beta)$ are, respectively,

$$\frac{df_i(\beta)}{d\beta} = \frac{k_+ B_i}{k_{-,i}}(2\beta - \mathcal{R}_T - \mathcal{L}_T)$$

$$\frac{d^2 f_i(\beta)}{d\beta^2} = \frac{2k_+ B_i}{k_{-,i}}$$

Considering this, eq.(8) and inequalities (6), the first and second derivatives or $F(\beta)$ are negative and positive, respectively. $F$ decreases monotonically in the interval of $\beta$ values that is physically meaningful; therefore, there is a single point in which $\beta = F(\beta)$, as required by eqs. (5) and (7). In other words, the system has a unique physically meaningful equilibrium point.

#### 2.3 Stability of the physically meaningful solution

To analyze the stability of the equilibrium point $\overline{\mathbf{C}} \in \mathbb{R}^M$ that corresponds to $\beta^*$, the only physically meaningful solution of eq. (5), we must analyze the Jacobian of function $G(\mathbf{C})$ (eq. (3)).

$$J_G(\overline{\mathbf{C}}) = \begin{bmatrix} \frac{\partial G_1(\overline{\mathbf{C}})}{\partial C_1} & \cdots & \frac{\partial G_1(\overline{\mathbf{C}})}{\partial C_M} \\ \vdots & \ddots & \vdots \\ \frac{\partial G_M(\overline{\mathbf{C}})}{\partial C_1} & \cdots & \frac{\partial G_M(\overline{\mathbf{C}})}{\partial C_M} \end{bmatrix} \tag{9}$$

There are two kinds of partial derivatives in this Jacobian matrix. On the diagonal, $i = j$:

$$\frac{\partial G_i(\overline{\mathbf{C}})}{\partial C_i} = [2\sum_{m=1}^{M} C_m - \mathcal{R}_T - \mathcal{L}_T] - \frac{k_{-,i}}{k_+ B_i} = [2\beta - \mathcal{R}_T - \mathcal{L}_T)] - \frac{k_{-,i}}{k_+ B_i}$$

; and for all other terms, $i \neq j$:

$$\frac{\partial G_i(\overline{\mathbf{C}})}{\partial C_j} = [2\beta - \mathcal{R}_T - \mathcal{L}_T]$$

Using

$$\mathbf{v} = (2\beta - \mathcal{R}_T - \mathcal{L}_T)\mathbb{1} = \begin{bmatrix} 2\beta - \mathcal{R}_T - \mathcal{L}_T \\ \vdots \\ 2\beta - \mathcal{R}_T - \mathcal{L}_T \end{bmatrix} \quad ; \quad \mathbf{D} = \begin{bmatrix} \frac{k_{-,1}}{k_+ B_1} & 0 & \dots & 0 \\ 0 & \frac{k_{-,2}}{k_+ B_2} & 0 & \vdots \\ \vdots & 0 & \ddots & \vdots \\ 0 & \dots & 0 & \frac{k_{-,M}}{k_+ B_M} \end{bmatrix},$$

we may write the Jacobian (eq. (9)) as

$$J_G(\overline{\mathbf{C}}) = \mathbf{v}\mathbb{1}^\dagger + (-\mathbf{D}) \tag{10}$$

In other words, the Jacobian matrix is a rank-one perturbation of a diagonal matrix. We then adapt a result from Ionascu [2] on this class of systems, as follows.

**Theorem 1.** *Let* $\mathbf{H} = \mathrm{diag}(h_i)_{i=1}^N$ *be a diagonal matrix and* $\mathbf{w}$ *any vector. Consider matrix* $\mathbf{T} = \mathbf{H} + \mathbf{w}\mathbb{1}^\dagger$ *that we obtain as a combination of the diagonal matrix* $\mathbf{H}$ *and the matrix* $\mathbf{w}\mathbb{1}^\dagger$*, that has all its columns equal to* $\mathbf{w}$*. If* $\mu \in \mathbb{R}$ *is an eigenvalue of* $\mathbf{T}$ *and* $\mu \notin \{h_1, h_2, \dots, h_N\}$*, then* $\mu$ *satisfies the equation*

$$\sum_{i=1}^{N} \frac{w_i}{\mu - h_i} = 1$$

*Proof.* Let's assume that $\mu \in \mathbb{R}$ is an eigenvalue of $\mathbf{T}$ and $\mathbf{x}$ is its associated eigenvector. Then,

$$\mathbf{T}\mathbf{x} = \mu\mathbf{x} = (\mathbf{H} + \mathbf{w}\mathbb{1}^\dagger)\mathbf{x} = \mathbf{H}\mathbf{x} + (\mathbb{1}^\dagger\mathbf{x})\mathbf{w}.$$

Thus,

$$(\mu I - \mathbf{D})\mathbf{x} = (\mathbb{1}^\dagger\mathbf{x})\mathbf{w} \tag{11}$$

, with $I$ referring to the $N \times N$ identity matrix. The $i$-th coordinate in eq. (11) is

$$(\mu - h_i)x_i = (\mathbb{1}^\dagger\mathbf{x})w_i$$

Then we have

$$\frac{w_i}{\mu - h_i} = \frac{x_i}{\mathbb{1}^\dagger\mathbf{x}}.$$

Summing over $i$ and considering that $\mathbb{1}^\dagger \mathbf{x} = \sum_{i=1}^{N} x_i$, we have

$$\sum_{i=1}^{N} \frac{w_i}{\mu - h_i} = \frac{1}{\mathbb{1}^\dagger \mathbf{x}} \sum_{i=1}^{N} x_i = 1$$

, which is what the theorem says. $\square$

To assess the stability of the unique physically meaningful equilibrium point $\overline{\mathbf{C}}$, we need to assess the eigenvalues of the Jacobian matrix $J_G(\overline{\mathbf{C}})$. We refer to the eigenvector that corresponds to eigenvalue $\mu$ as $\mathbf{x}$. Using $\gamma = 2\beta - \mathcal{R}_T - \mathcal{L}_T$, eq. (10) and following the eigenvalue equation, we have

$$J_G(\overline{\mathbf{C}}))\mathbf{x} = (\mathbf{v}\mathbb{1}^\dagger - \mathbf{D})\mathbf{x} = (\gamma\mathbb{1}\mathbb{1}^\dagger - \mathbf{D})\mathbf{x} = \mu\mathbf{x} \tag{12}$$

$$\implies \gamma\mathbb{1}(\mathbb{1}^\dagger\mathbf{x}) - \mathbf{D}\mathbf{x} = \mu\mathbf{x} \tag{13}$$

Calling $d_i$ the $i$-th element of the diagonal of matrix $\mathbf{D}$. For any element $j$ of $\mathbf{x}$ we have

$$\gamma(\sum_{i=1}^{M} x_i) - d_j x_j = \mu x_j$$
$$\implies \gamma(\sum_{i=1}^{M} x_i) = x_j(\mu + d_j)$$

We now have two cases:

i) $(\mu + d_j) = 0$ for at least one of the diagonal entries $d_j$

ii) $(\mu + d_j) \neq 0$ for all the diagonal entries $d_j$

##### 2.3.1 Case i): $(\mu + d_j) = 0$ for at least one $d_j$

In this case, $\mu = -d_j$. Since all the values in the diagonal of $\mathbf{D}$ are positive, this implies that $\mu < 0$.

##### 2.3.2 Case ii): $(\mu + d_j) \neq 0$ for all entries $d_j$

Applying theorem (1) to the Jacobian matrix as described in eq. (10), we get:

$$\sum_{i=1}^{M} \frac{v_i}{\mu + (\mathbf{D})_{i,i}} = \sum_{i=1}^{M} \frac{\gamma}{\mu + d_i} = 1$$

Under physically valid conditions and for all $1 \leq i \leq M$, $\gamma < 0$ and $d_i > 0$. Hence, theorem (1) requires that $\mu < 0$. Therefore, all the eigenvalues of the Jacobian $J_G(\overline{\mathbf{C}})$ are negative and the unique physically meaningful equilibrium point is stable.

### 3 Plastogenetic congruence

Regarding RNA secondary structures, plastogenetic congruence is the association between the structural variants produced by thermal fluctuations and mutation. We assessed how strong is this association for sequences that fold into the structures, biological and random, that we opted to study. The issue is important because plastogenetic congruence is an important condition for the appearance of neutral confinement [3]. First we found that RNA molecules with a strong robustness to thermal fluctuations tend to have a high mutational robustness (Fig. S2a, Table S1). Moreover, sequences that can access many new structures through single mutations also usually have many different structures in their plastic repertoire (Fig. S2b, Table S2). In addition, we assessed whether mutation and thermal fluctuations tend to produce the same phenotypes. We did so by evaluating the overlap coefficient between the mutational and plastic repertoires, $\Xi_i$ and $\Phi_i$, for each sequence $s_i$ in our sample. This coefficient equals the number of structures that appear in both

**Table S1. Correlation between a sequence's mutational robustness and its thermodynamic stability.**

| Structure | Mutational robustness | Thermodynamic stability* | Correlation |
|---|---|---|---|
| | Median (1st, 3rd quartiles) | Median (1st, 3rd quartiles) | Pearson's $r$; $p$-value |
| CPEB3 ribozyme | 0.275 (0.227, 0.329) | 0.105 (0.066, 0.17) | 0.69; $< 10^{-6}$ |
| DsrA | 0.263 (0.226, 0.292) | 0.129 (0.078, 0.213) | 0.522; $< 10^{-6}$ |
| Hepatitis ribozyme | 0.262 (0.217, 0.307) | 0.066 (0.039, 0.112) | 0.622; $< 10^{-6}$ |
| snoRNA | 0.277 (0.227, 0.33) | 0.056 (0.033, 0.096) | 0.669; $< 10^{-6}$ |
| tRNAphe | 0.265 (0.219, 0.311) | 0.152 (0.092, 0.253) | 0.685; $< 10^{-6}$ |
| ran0 | 0.283 (0.239, 0.328) | 0.165 (0.102, 0.263) | 0.637; $< 10^{-6}$ |
| ran1 | 0.211 (0.167. 0.25) | 0.107 (0.067, 0.173) | 0.63; $< 10^{-6}$ |
| ran2 | 0.289 (0.239, 0.344) | 0.14 (0.086, 0.227) | 0.663; $< 10^{-6}$ |
| ran3 | 0.239 (0.2, 0.283) | 0.159 (0.1, 0.252) | 0.654; $< 10^{-6}$ |
| ran4 | 0.317 (0.261, 0.367) | 0.184 (0.114, 0.296) | 0.712; $< 10^{-6}$ |
| ran5 | 0.261 (0.217, 0.317) | 0.143 (0.092, 0.222) | 0.656; $< 10^{-6}$ |
| ran6 | 0.156 (0.128, 0.194) | 0.088 (0.055, 0.141) | 0.615; $< 10^{-6}$ |
| ran7 | 0.261 (0.217, 0.311) | 0.155 (0.096, 0.253) | 0.624; $< 10^{-6}$ |
| ran8 | 0.222 (0.178, 0.261) | 0.103 (0.065, 0.164) | 0.591; $< 10^{-6}$ |
| ran9 | 0.233 (0.194, 0.272) | 0.134 (0.085, 0.211) | 0.68; $< 10^{-6}$ |

* We evaluate thermodynamic stability as the Boltzmann probability of a sequence's MFES.

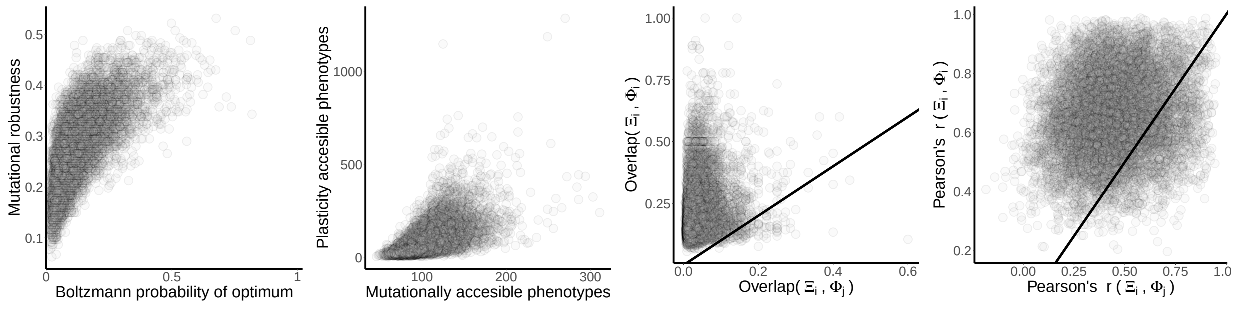

(a) (b) (c) (d)

**Fig. S2. Plastogenetic congruence in sequences that fold into the structure of the CPEB3 ribozyme as their MFES.** (a) Thermodynamic stability is positively associated to a molecule's ability to withstand mutations. Pearson's $r = 0.69$; $p < 10^{-6}$. (b) The number of phenotypes that a sequence can access through mutation is positively correlated with the number of phenotypes in its plastic repertoire. $r = 0.6$; $p < 10^{-6}$. (c) and (d) Points above the identity line (black diagonal) indicate sequences with a greater value on the vertical axis than on the horizontal axis. (c) The overlap between a sequence $s_i$'s mutational and plastic repertoires, $\Xi_i$ and $\Phi_i$, is most often greater than the one between $\Xi_i$ and the plastic repertoire $\Phi_j$ of a random sequence $s_j$ that yields the same MFES as $s_i$. Wilcoxon's signed rank test. $W = 49,890,458$; $p < 10^{-6}$. (d) Structures that have greater mutational access from a sequence $s_i$ tend to have a greater Boltzmann probability in $s_i$'s plastic repertoire. This correlation is greater than the one between mutational access from $s_i$ and the Boltzmann probability for structures in the plastic repertoire of a random sequence $s_j$ yielding the same MFES as $s_i$. $W = 45,534,700$; $p < 10^{-6}$.

**Table S2. Correlation between the number of structures in the mutational and plastic repertoires.**

| Structure | Mutational repertoire $\Xi$ | Plastic repertoire $\Phi$ | Correlation |
|---|---|---|---|
| | Median (1st, 3rd quartiles) | Median (1st, 3rd quartiles) | Pearson's $r$; $p$-value |
| CPEB3 ribozyme | 104 (91, 121) | 73 (43, 122) | 0.598; $< 10^{-6}$ |
| DsrA | 117 (100,142) | 52 (29,92) | 0.452; $< 10^{-6}$ |
| Hepatitis ribozyme | 148 (126, 181) | 124 (70, 220) | 0.556; $< 10^{-6}$ |
| snoRNA | 172 (148, 205) | 167 (92, 292) | 0.508; $< 10^{-6}$ |
| tRNAphe | 107 (94, 123) | 48 (26, 83) | 0.503; $< 10^{-6}$ |
| ran0 | 79 (69, 91) | 40 (23, 69) | 0.498; $< 10^{-6}$ |
| ran1 | 98 (87, 113) | 67 (40, 111) | 0.557; $< 10^{-6}$ |
| ran2 | 89 (77, 105) | 48 (27, 81) | 0.582; $< 10^{-6}$ |
| ran3 | 84 (74, 95) | 42 (25, 71) | 0.5; $< 10^{-6}$ |
| ran4 | 78 (68, 90) | 35 (19, 60) | 0.592; $< 10^{-6}$ |
| ran5 | 87 (76, 99) | 50 (30, 80) | 0.567; $< 10^{-6}$ |
| ran6 | 107 (93, 132) | 85 (51, 139) | 0.59; $< 10^{-6}$ |
| ran7 | 88 (78, 101) | 43 (25, 74) | 0.527; $< 10^{-6}$ |
| ran8 | 99 (86, 116) | 69 (41, 114) | 0.554; $< 10^{-6}$ |
| ran9 | 88 (79, 99) | 52 (31, 85) | 0.512; $< 10^{-6}$ |

**Table S3. Overlap coefficient between mutational and plastic repertoires.**

| Structure | Overlap($\Xi_i$,$\Phi_i$) (same sequence) | Overlap($\Xi_i$,$\Phi_j$) (different sequence) | Wilcoxon signed-rank test |
|---|---|---|---|
| | Median (1st, 3rd quartiles) | Median (1st, 3rd quartiles) | $W$; $p$-value |
| CPEB3 ribozyme | 0.188 (0.147, 0.267) | 0.033 (0.02, 0.053) | 49,890,458; $< 10^{-6}$ |
| DsrA | 0.3 (0.206,0.435) | 0.102 (0.063, 0.167) | 47,562,378; $< 10^{-6}$ |
| Hepatitis ribozyme | 0.17 (0.14, 0.23) | 0.035 (0.023, 0.055) | 49,623,530; $< 10^{-6}$ |
| snoRNA | 0.134 (0.112, 0.176) | 0.017 (0.011, 0.028) | 49,939,505; $< 10^{-6}$ |
| tRNAphe | 0.292 (0.198, 0.429) | 0.065 (0.037, 0.115) | 49,063,250; $< 10^{-6}$ |
| ran0 | 0.283 (0.202, 0.41) | 0.077 (0.047, 0.0125) | 48,913,171; $< 10^{-6}$ |
| ran1 | 0.22 (0.169, 0.316) | 0.045 (0.028, 0.074) | 49,725,388; $< 10^{-6}$ |
| ran2 | 0.267 (0.193, 0.387) | 0.06 (0.035, 0.103) | 49,284,242; $< 10^{-6}$ |
| ran3 | 0.3 (0.213, 0.435) | 0.07 (0.043, 0.116) | 49,201,618; $< 10^{-6}$ |
| ran4 | 0.318 (0.219, 0.467) | 0.075 (0.044, 0.13) | 48,654,930; $< 10^{-6}$ |
| ran5 | 0.25 (0.182, 0.364) | 0.061 (0.036, 0.1) | 49,289,944; $< 10^{-6}$ |
| ran6 | 0.203 (0.163, 0.279) | 0.045 (0.029, 0.07) | 49,709,868; $< 10^{-6}$ |
| ran7 | 0.282 (0.196, 0.414) | 0.056 (0.033, 0.1) | 49,383,348; $< 10^{-6}$ |
| ran8 | 0.216 (0.168, 0.306) | 0.043 (0.026, 0.069) | 49,806,970; $< 10^{-6}$ |
| ran9 | 0.233 (0.176, 0.333) | 0.05 (0.032, 0.082) | 49,430,214; $< 10^{-6}$ |

**Table S4. Correlation between mutational access and Boltzmann probability.**

| Structure | Pearson's $r$($\Xi_i$,$\Phi_i$) (same sequence) | Pearson's $r$($\Xi_i$,$\Phi_j$) (different sequence) | Wilcoxon signed-rank test |
|---|---|---|---|
| | Median (1st, 3rd quartiles) | Median (1st, 3rd quartiles) | $W$; $p$-value |
| CPEB3 ribozyme | 0.658 (0.561, 0.76) | 0.441 (0.319, 0.578) | 45,534,700; $< 10^{-6}$ |
| DsrA | 0.684 (0.579, 0.787) | 0.442 (0.317, 0.581) | 45,777,783; $< 10^{-6}$ |
| Hepatitis ribozyme | 0.586 (0.49, 0.69) | 0.343 (0.23, 0.473) | 46,704,526; $< 10^{-6}$ |
| snoRNA | 0.547 (0.453, 0.649) | 0.362 (0.257, 0.488) | 43,916,914; $< 10^{-6}$ |
| tRNAphe | 0.716 (0.608, 0.821) | 0.531 (0.401, 0.678) | 42,997,799; $< 10^{-6}$ |
| ran0 | 0.712 (0.606, 0.818) | 0.519 (0.39, 0.665) | 43,713,816; $< 10^{-6}$ |
| ran1 | 0.649 (0.548, 0.749) | 0.342 (0.22, 0.486) | 48,431,622; $< 10^{-6}$ |
| ran2 | 0.709 (0.606, 0.812) | 0.437 (0.307, 0.588) | 46,938,612; $< 10^{-6}$ |
| ran3 | 0.712 (0.611, 0.81) | 0.492 (0.366, 0.629) | 45,820,950; $< 10^{-6}$ |
| ran4 | 0.753 (0.644, 0.856) | 0.568 (0.433, 0.713) | 43,260,356; $< 10^{-6}$ |
| ran5 | 0.692 (0.6, 0.789) | 0.504 (0.39, 0.632) | 44,618,696; $< 10^{-6}$ |
| ran6 | 0.605 (0.506, 0.702) | 0.236 (0.122, 0.369) | 49,684,096; $< 10^{-6}$ |
| ran7 | 0.718 (0.613, 0.819) | 0.464 (0.334, 0.612) | 46,397,126; $< 10^{-6}$ |
| ran8 | 0.656 (0.552, 0.753) | 0.32 (0.2, 0.459) | 48,891,286; $< 10^{-6}$ |
| ran9 | 0.68 (0.576, 0.781) | 0.462 (0.338, 0.595) | 45,928,684 $< 10^{-6}$ |

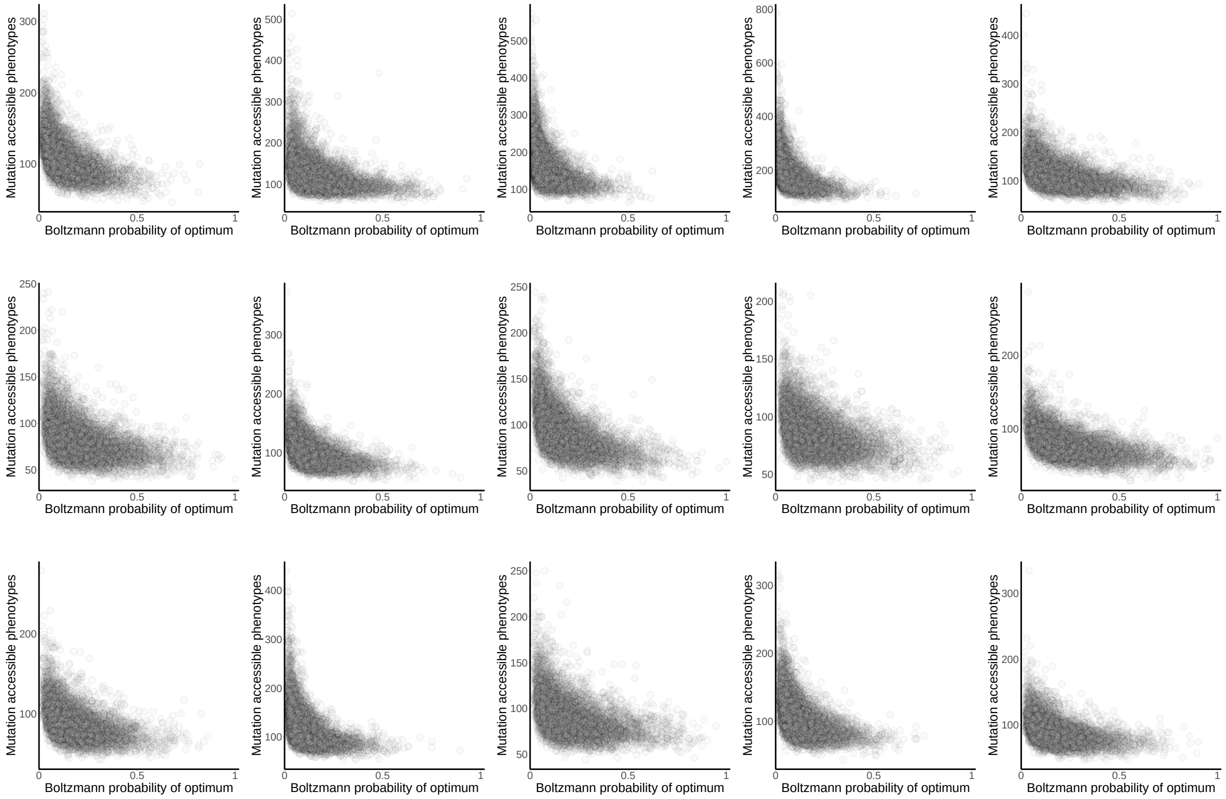

**Fig. S3. Thermodynamic stability and mutational access to new structures are negatively correlated.** We assess thermodynamic stability as a sequence's Boltzmann probability for a specific secondary structure picked a priori. We evaluated mutational access as the cardinality of a sequence's mutational repertoire. The first row presents data for sequences that fold into biological structures in the following order: CPEB3 ribozyme, DsrA, Hepatitis ribozyme, snoRNA, tRNAphe. The second and third row show data for sequences folding into structures ran0-9.

repertoires divided by the cardinality of the smaller set. To assess the significance of such an overlap, we compared it to the overlap between the sequence's mutational repertoire $\Xi_i$ and the plastic repertoire $\Phi_j$ of a different random sequence $s_j$ that folds into the same MFES as $s_i$. The overlap is clearly beyond random expectation for all the structures that we studied. For example, the median for this measure of plastogenetic congruence in sequences that fold into the structure of the CPEB3 ribozyme is 0.188. In contrast, the overlap for the corresponding mutational and plastic repertoires of different sequences has a median of 0.033. Fig. S2c and Table S3 show that the difference is highly significant for sequences folding into, respectively, the CPEB3 ribozyme structure and any of the structures that we study.

Even though the overlap between $\Xi_i$ and $\Phi_i$ is highly statistically significant, the magnitude of its median indicates that it often happens that an appreciable number of the structures in one of these repertoires is absent from the other. We hypothesized that plastogenetic congruence is specially relevant for those structures easily accessible through mutations or thermal fluctuations. In this case, structures present in only one of the repertoires would be the ones with a low mutational access (if in $\Xi_i$), or a low Boltzmann probability (if in $\Phi_i$). We thus studied whether there are more mutations that lead from a sequence $s_i$ to sequences that have a structure $\upsilon$ as its MFES when $\upsilon$'s Boltzmann probability is high in $s_i$'s plastic repertoire. For each sequence $s_i$ in our sample we performed a Pearson correlation analysis between mutational access in $s_i$'s mutational repertoire and Boltzmann probability in $s_i$'s plastic repertoire. We found that the correlation was positive in all $10^4$ sequences for each of the 15 structures in our study. Mutational and plastic repertoires' alignment in the production of variation has a strong sequence specific component: mutational access and access through plasticity are significantly better correlated when assessed in the mutational ($\Xi_i$) and plastic ($\Phi_i$) repertoires of the same sequence $s_i$ than when evaluated in repertoires $\Xi_i$ and $\Phi_j$, pertaining to different sequences $s_i$ and $s_j$ that yield the same MFES (Fig. S2d and Table S4).

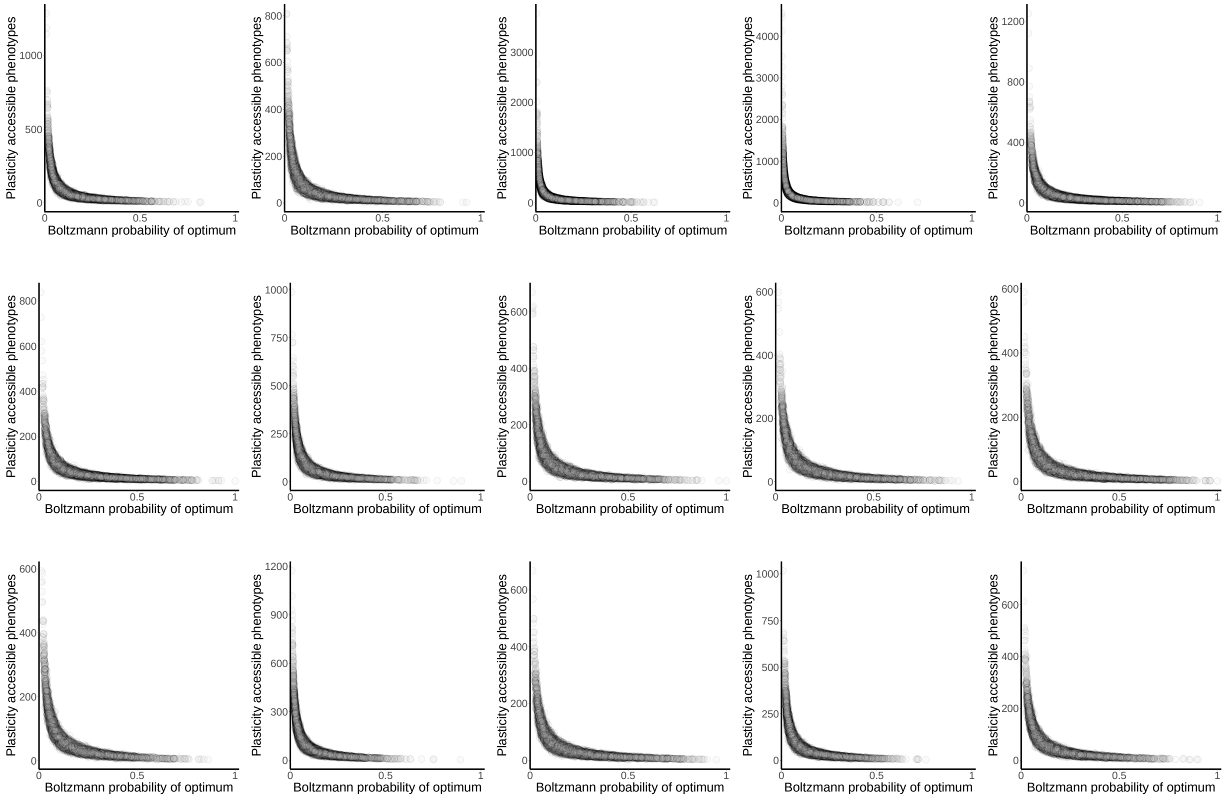

**Fig. S4. Thermodynamic stability and access through plasticity to new structures are negatively correlated.** We assess thermodynamic stability as a sequence's Boltzmann probability for a specific secondary structure picked a priori. We evaluated access through plasticity as the cardinality of a sequence's plastic repertoire. The first row presents data for sequences that fold into biological structures in the following order: CPEB3 ribozyme, DsrA, Hepatitis ribozyme, snoRNA, tRNAphe. The second and third row show data for sequences folding into structures ran0-9.

We also assessed the association between thermodynamic stability and access to new phenotypes through mutation or through plasticity. We found that the size of the mutational and plastic repertoires $\Xi_i$ and $\Phi_i$ tends to decrease with a sequence's Boltzmann probability of the optimal structure (Figs. S3 and S4).

### 4 Analyses of sequences sampled with a random mutational walk subsequent to inverse folding

The many RNA sequences that can fold into the same MFES often arrange in large mutationally connected sets. One could traverse any such set through single mutations without losing the ability to yield the same MFES [4–6]. The inverse-fold algorithm [7] is a fast, computationally effective method that allows obtaining random samples of sequences that produce a specific MFES $\tau$ from all the different mutationally connected sets associated to it. Notwithstanding its many advantages, using the inverse-fold algorithm to sample sequences that yield a specific MFES $\tau$ deviates from uniformity [8; 9], albeit slightly [10].

In reference [9] the authors propose a random mutational walk after inverse-folding as a method to avoid the algorithm's bias. We followed such proposal to assess the possibility that our results were an artifact due to the sampling method. For each sequence in our sample we performed an initial inverse-fold followed by $50L$ mutational steps that preserved the sequence's MFES, $L$ corresponding to the sequence's length. Figures S5 and S6 indeed show that results for sequences that fold into the structure of the CPEB3 ribozyme are qualitatively the same in this case as for sequences sampled only with inverse fold, reported in the main text. This observation supports that our findings are not a spurious artifact of the sampling method.

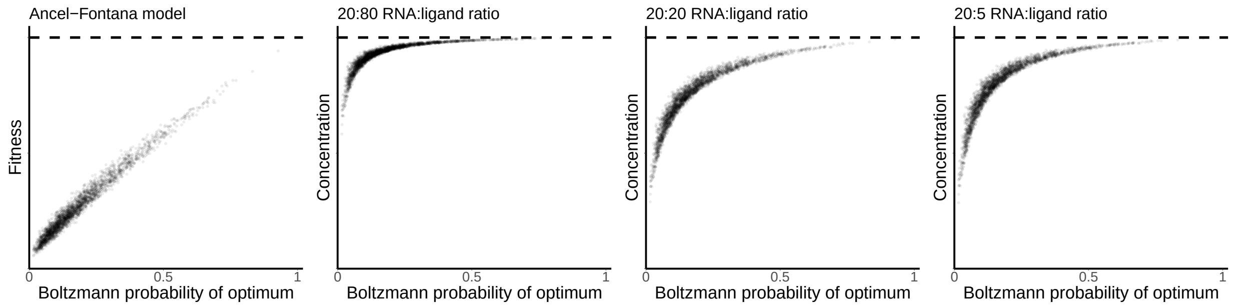

**Fig. S5. Fitness as a function of thermodynamic stability in sequences that fold into the structure of the CPEB3 ribozyme as their MFES and that were sampled by a random mutational walk after inverse-folding.** We assess thermodynamic stability as a sequence's Boltzmann probability for the structure of the CPEB3 ribozyme. The horizontal dashed line indicates the maximum fitness value in each scenario.

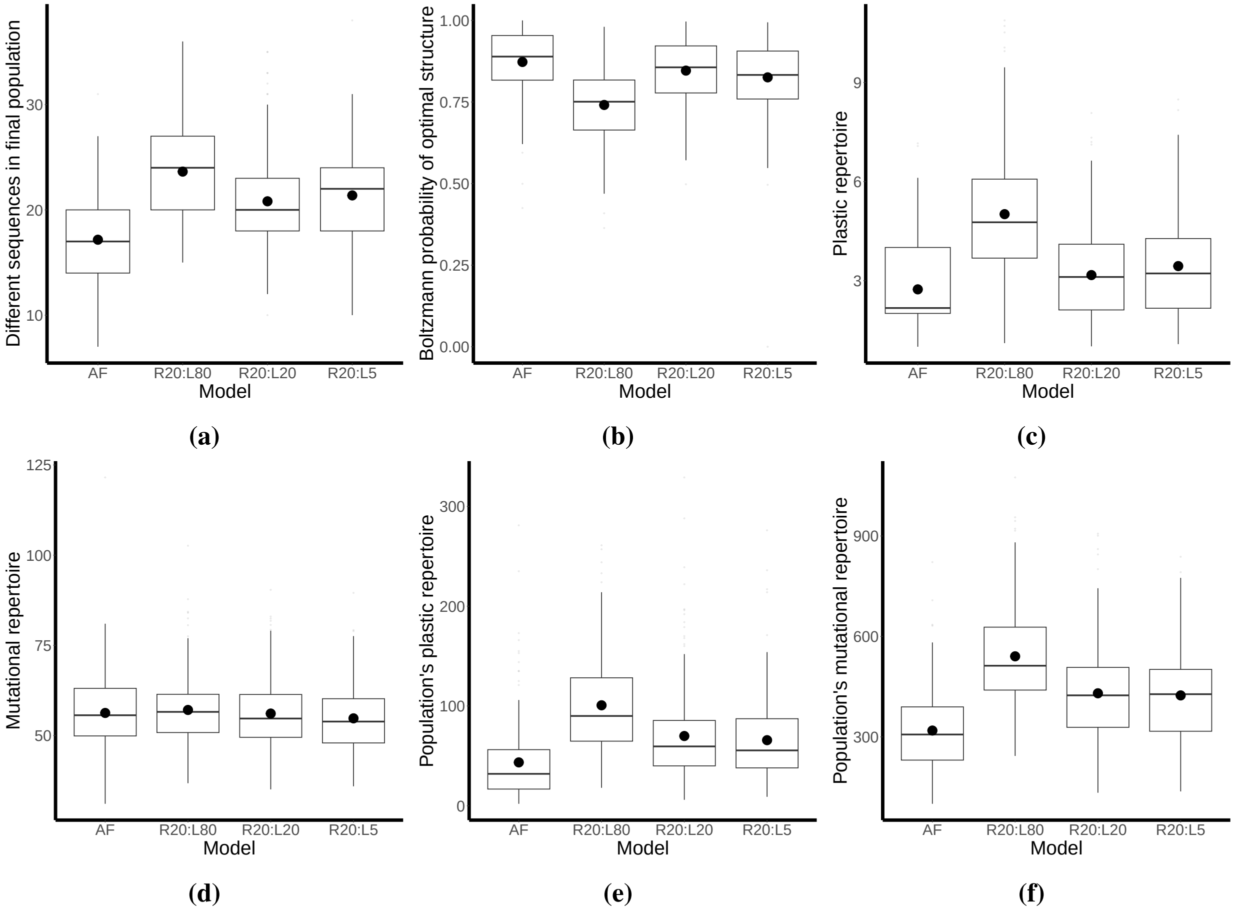

**Fig. S6. Evolution under stabilizing selection in favor of the CPEB3 ribozyme structure under different scenarios.** Initial sequences were sampled using inverse-folding followed by a long random mutational walk. AF: Ancel-Fontana model; RX:LY: RNA-ligand scenario with X and Y as the constant concentrations of the RNA molecule and the ligand, respectively. (a) Number of distinct sequences in the population at the end of evolution. (b) Average thermodynamic stability at the end of evolution. (c,d). Cardinality of the plastic and mutational repertoires, respectively. (e,f). Population's access to new phenotypes through plasticity and mutation, respectively. We define the population's plastic, or mutational, repertoire as the union of the plastic, or mutational, repertoires of all the sequences that comprise a population.
