## Supplementary Figures for "RNA-ligand complexes and the attenuation of neutral confinement in the evolution of RNA secondary structures"

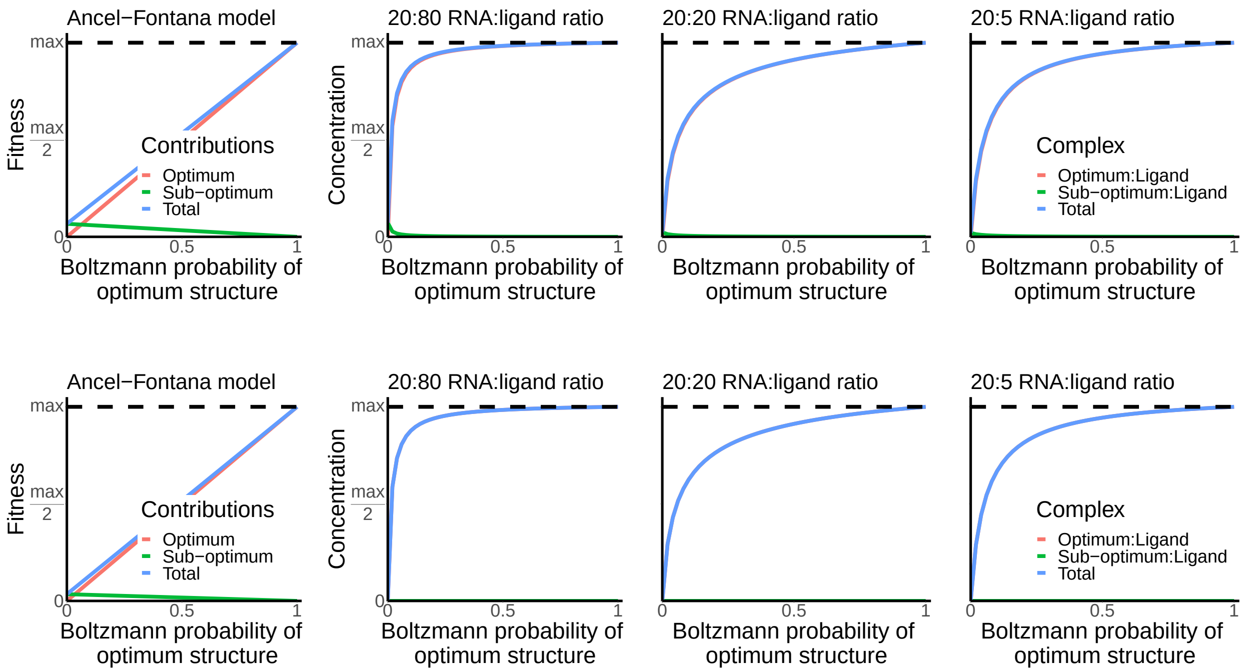


**Fig. SF1. Fitness as a function of thermodynamic stability of a sequence's MFES with an increased structural distance between optimal and alternative structures.** We assume that all sequences have two different structures in their plastic repertoire: an optimal structure $\tau$ and a second structure $\upsilon$ at a fixed structural distance $d$ from $\tau$. Green and red lines indicate $\upsilon$'s and $\tau$'s contribution to fitness, respectively. The blue line refers to the sequence's total fitness. $d = 20$ and $d = 40$ in the top and bottom rows, respectively.

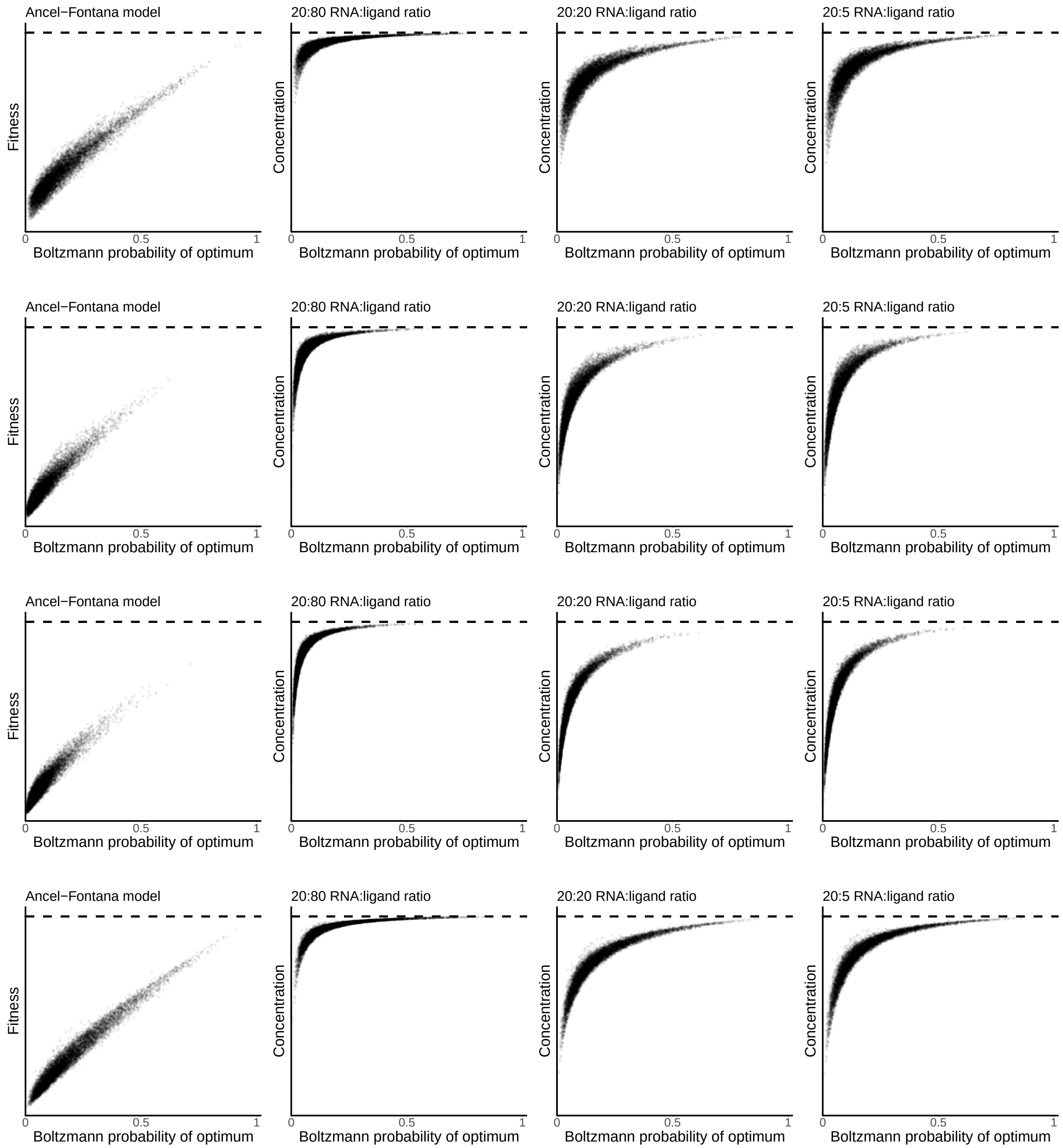


**Fig. SF2. Fitness as a function of thermodynamic stability in sequences that fold into biological structures as their MFES.** We assess thermodynamic stability as a sequence's Boltzmann probability for a biological structure. The horizontal dashed line indicates the maximum fitness value in each scenario. In the Ancel-Fontana model the relationship between thermodynamic stability and fitness is near-linear. In the RNA-ligand model, fitness is a concave function of thermodynamic stability. Each row presents data for sequences that fold into one biological structure. In order, the structures are: DsrA, Hepatitis ribozyme, snoRNA, tRNAphe.

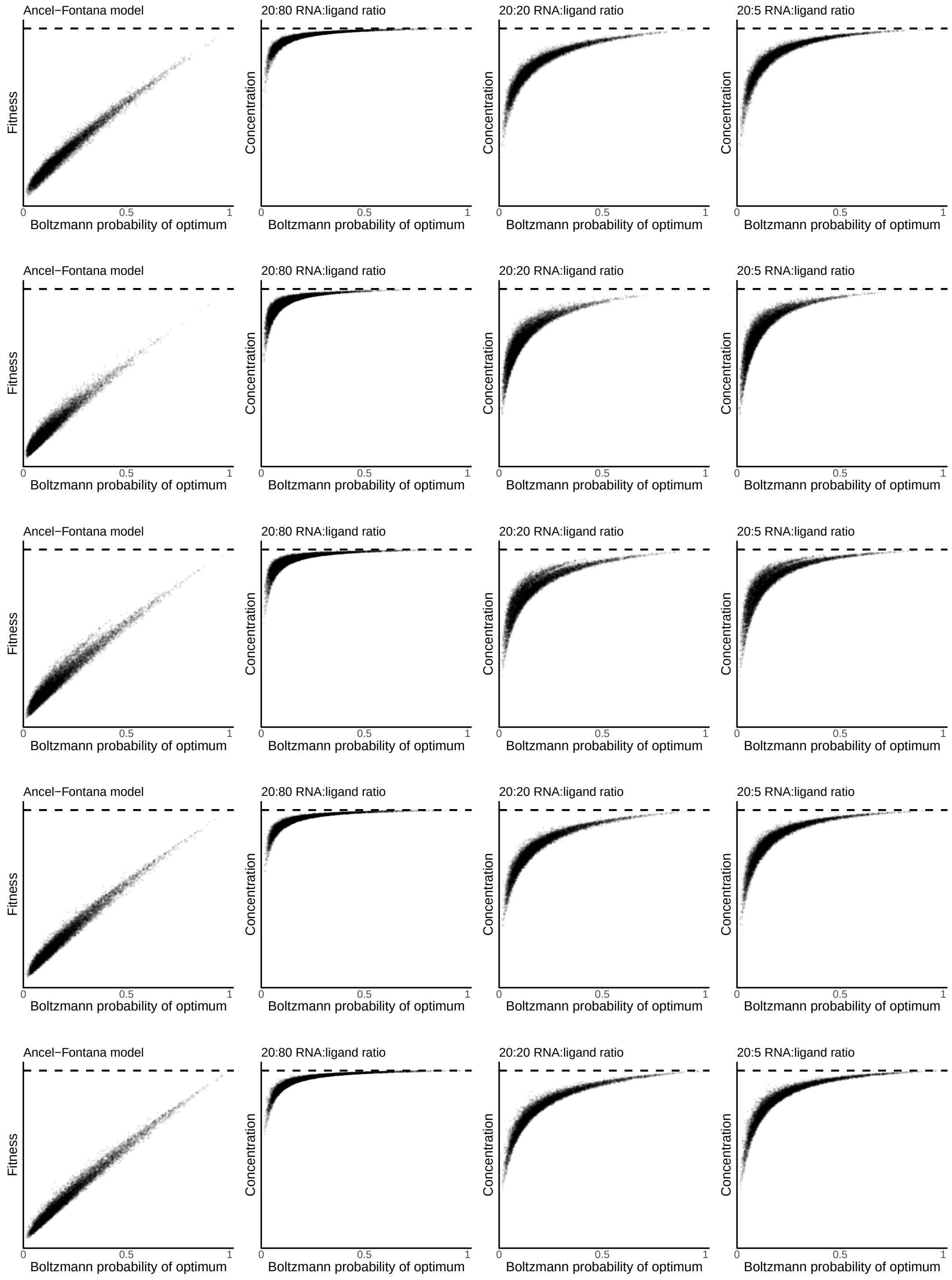
Ancel−Fontana model
20:80 RNA:ligand ratio
20:20 RNA:ligand ratio
20:5 RNA:ligand ratio
Fitness
Concentration
0
0.5
1
Boltzmann probability of optimum

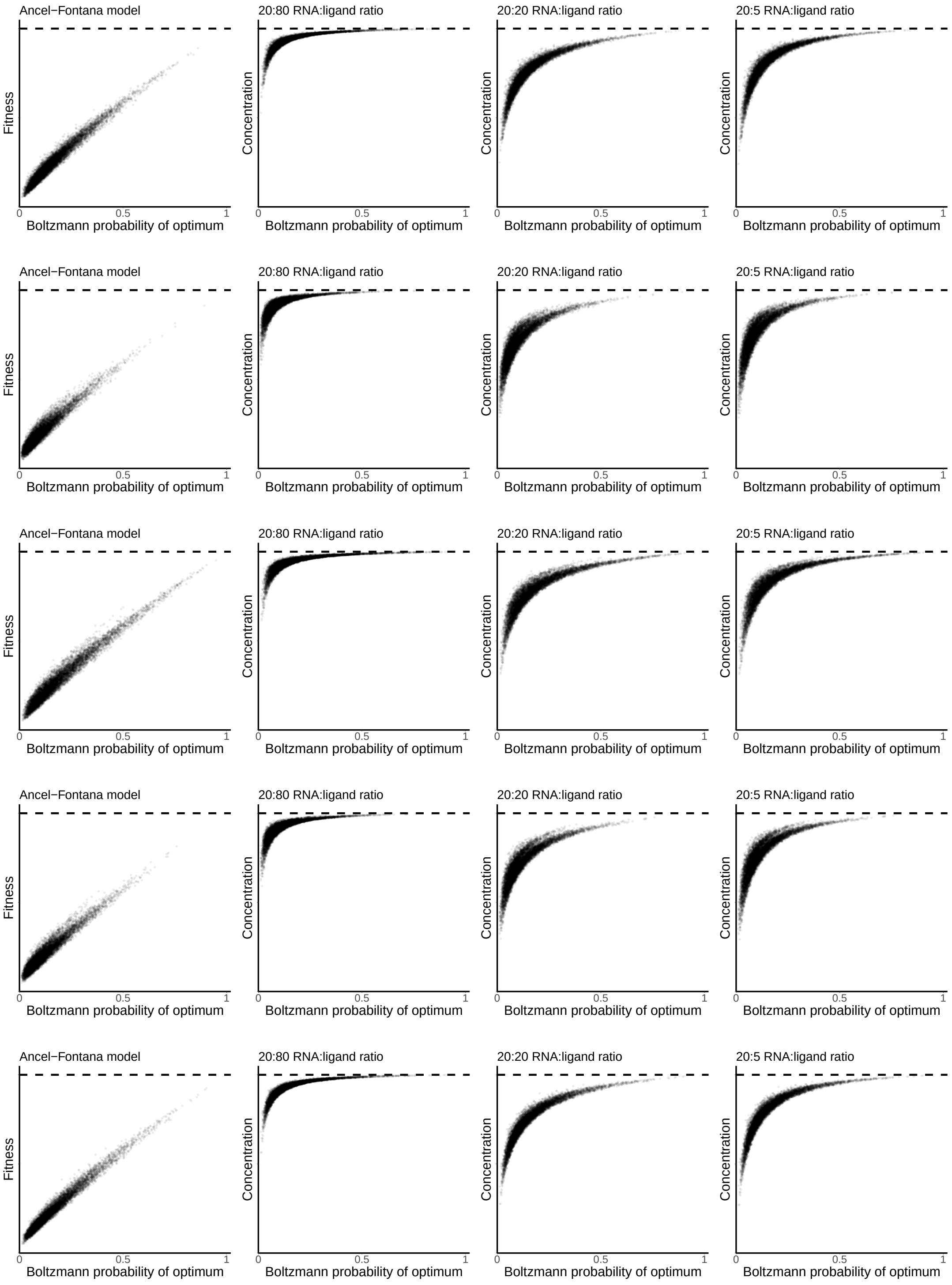


**Fig. SF3. Fitness as a function of thermodynamic stability in sequences that fold into specific random structures as their MFES.** Each row presents data for sequences that fold into one of the structures in Table ST2, appearing in the same order as in that table.

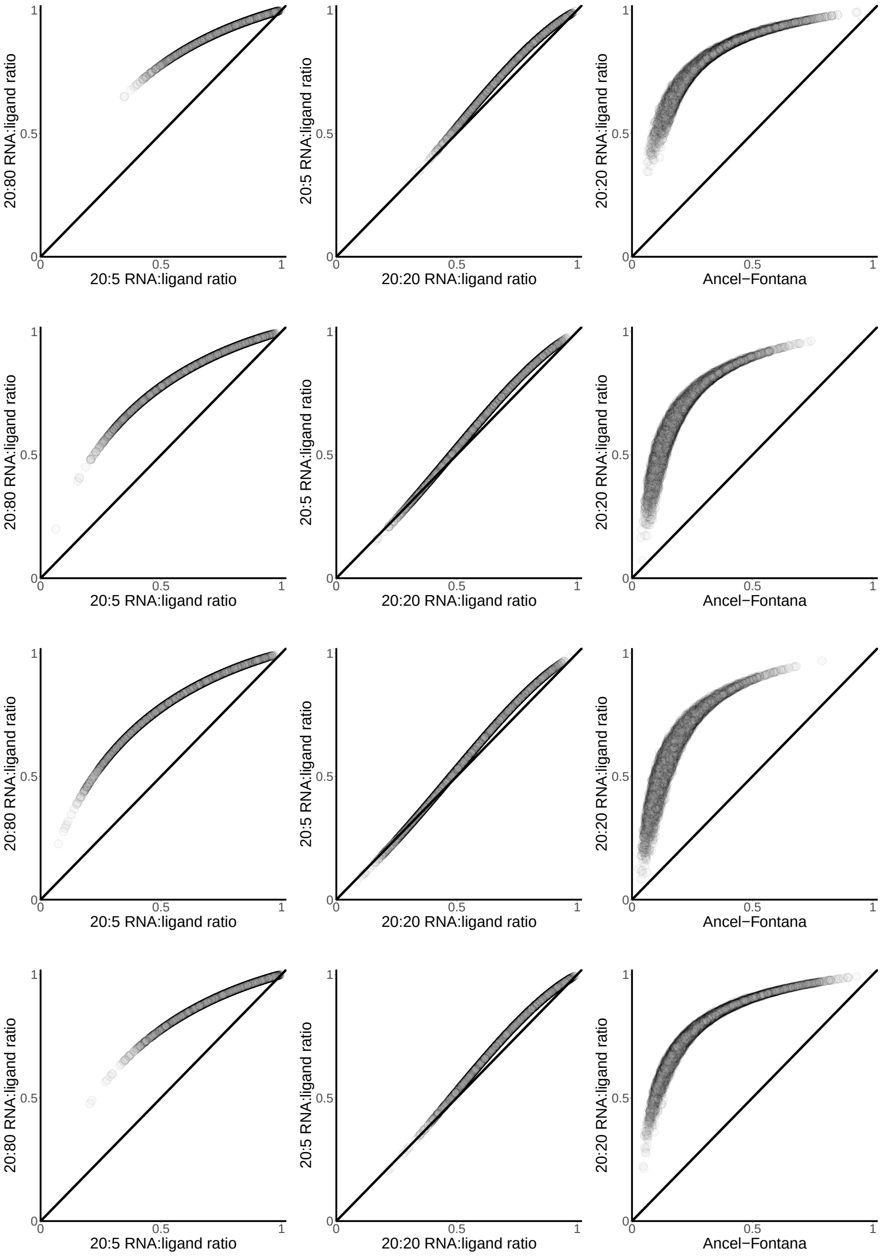


**Fig. SF4. Comparison of relative fitness of sequences that fold into specific biological structures under different models or ligand concentrations.** In all panels, the diagonal indicates the identity line. Points above the diagonal refer to sequences with a higher fitness under the model/ligand concentration that corresponds to the vertical axis than under conditions described in the horizontal axis. Fitness values are normalized with respect to the maximum fitness value for the respective model. Each row presents data for sequences that fold into one structure. In order, the structures are: DsrA, Hepatitis ribozyme, snoRNA, tRNAphe.

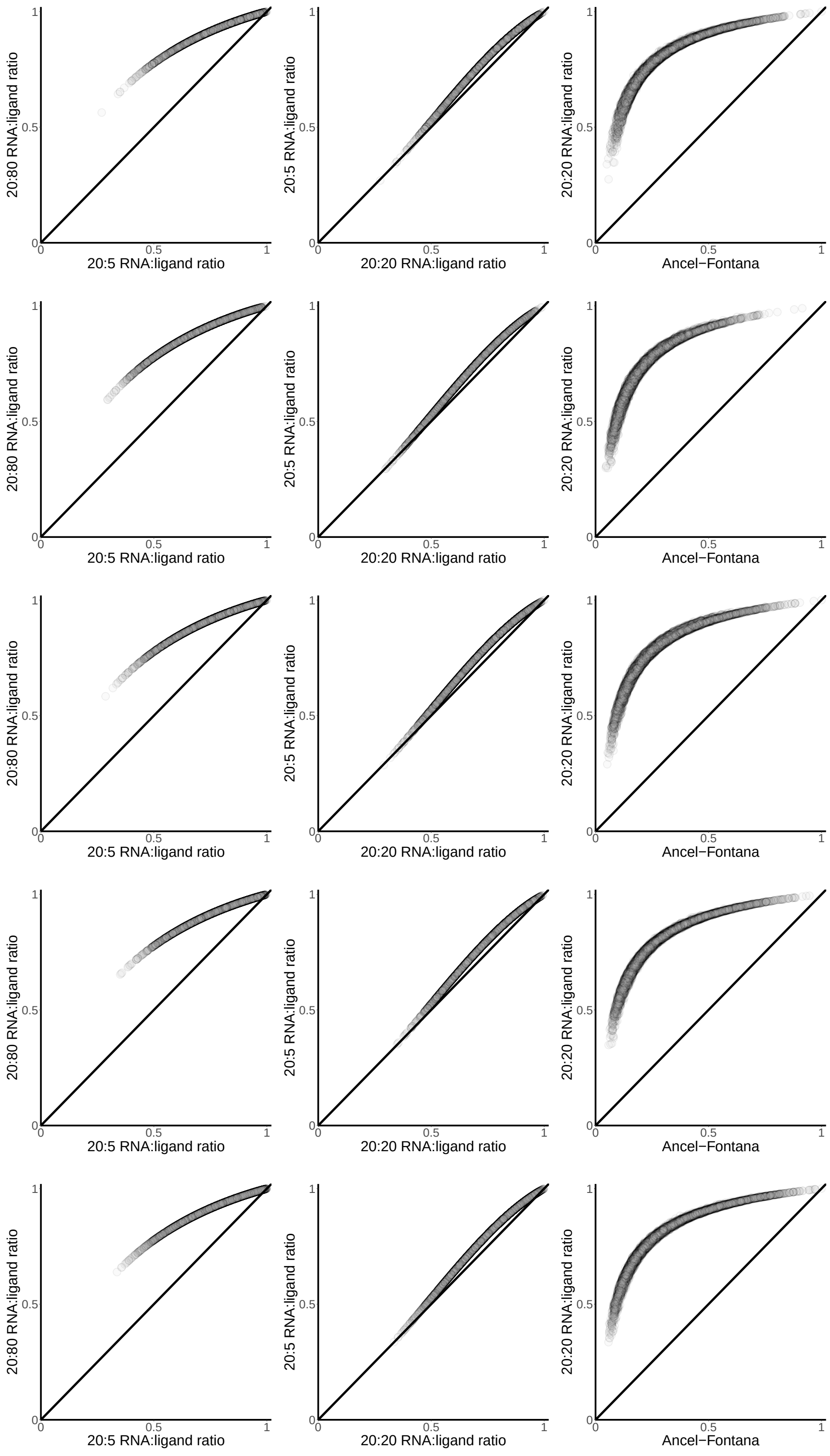
20:80 RNA:ligand ratio
20:5 RNA:ligand ratio
20:5 RNA:ligand ratio
20:20 RNA:ligand ratio
20:20 RNA:ligand ratio
Ancel−Fontana
0
0.5
1

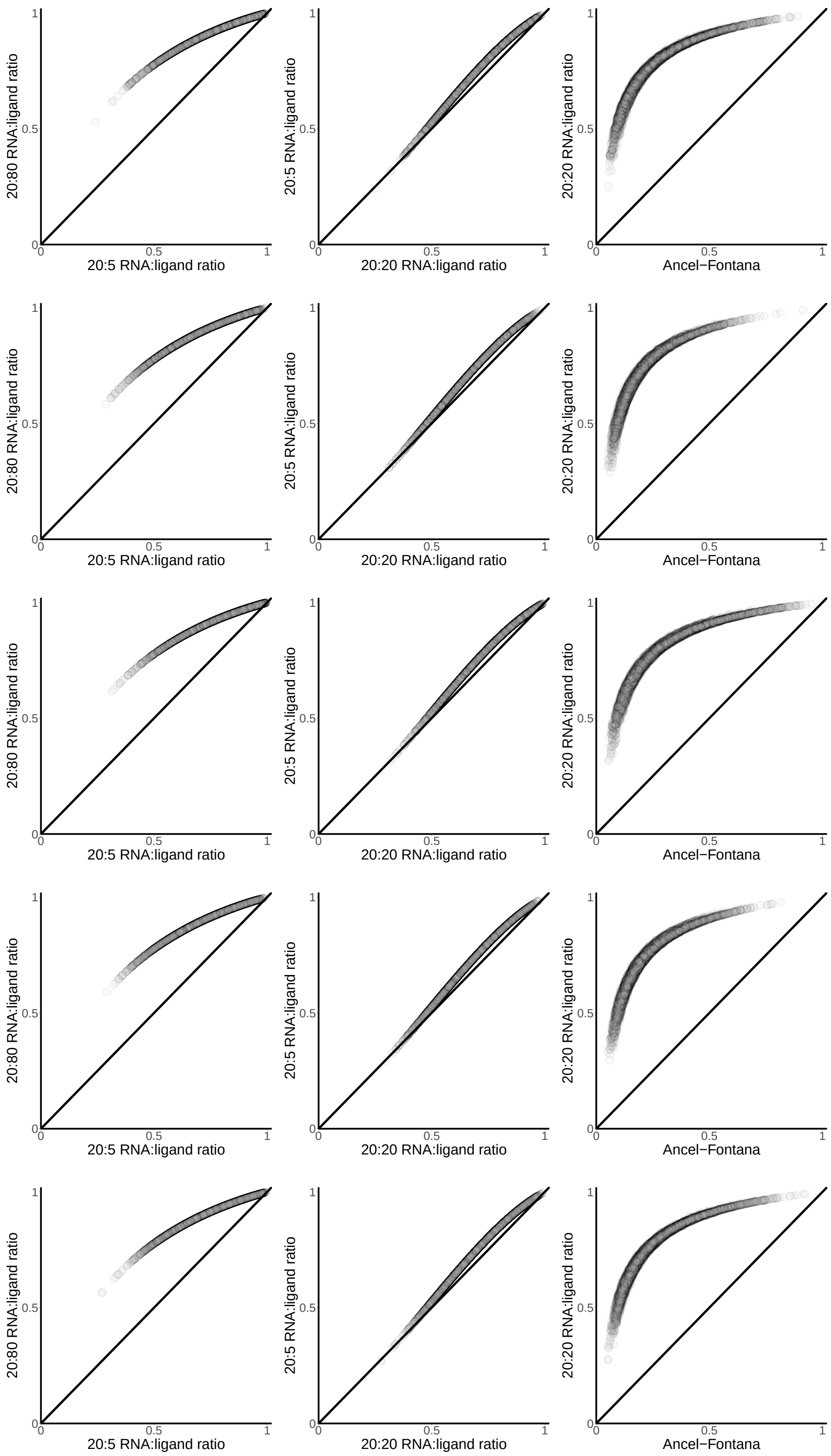


**Fig. SF5. Comparison of relative fitness of sequences that fold into our randomly-picked structures under different models or ligand concentrations.** Fitness values are normalized with respect to the maximum fitness value for the respective model. Each row presents data for sequences that fold into one of the structures in Table ST2, appearing in the same order as in that table.

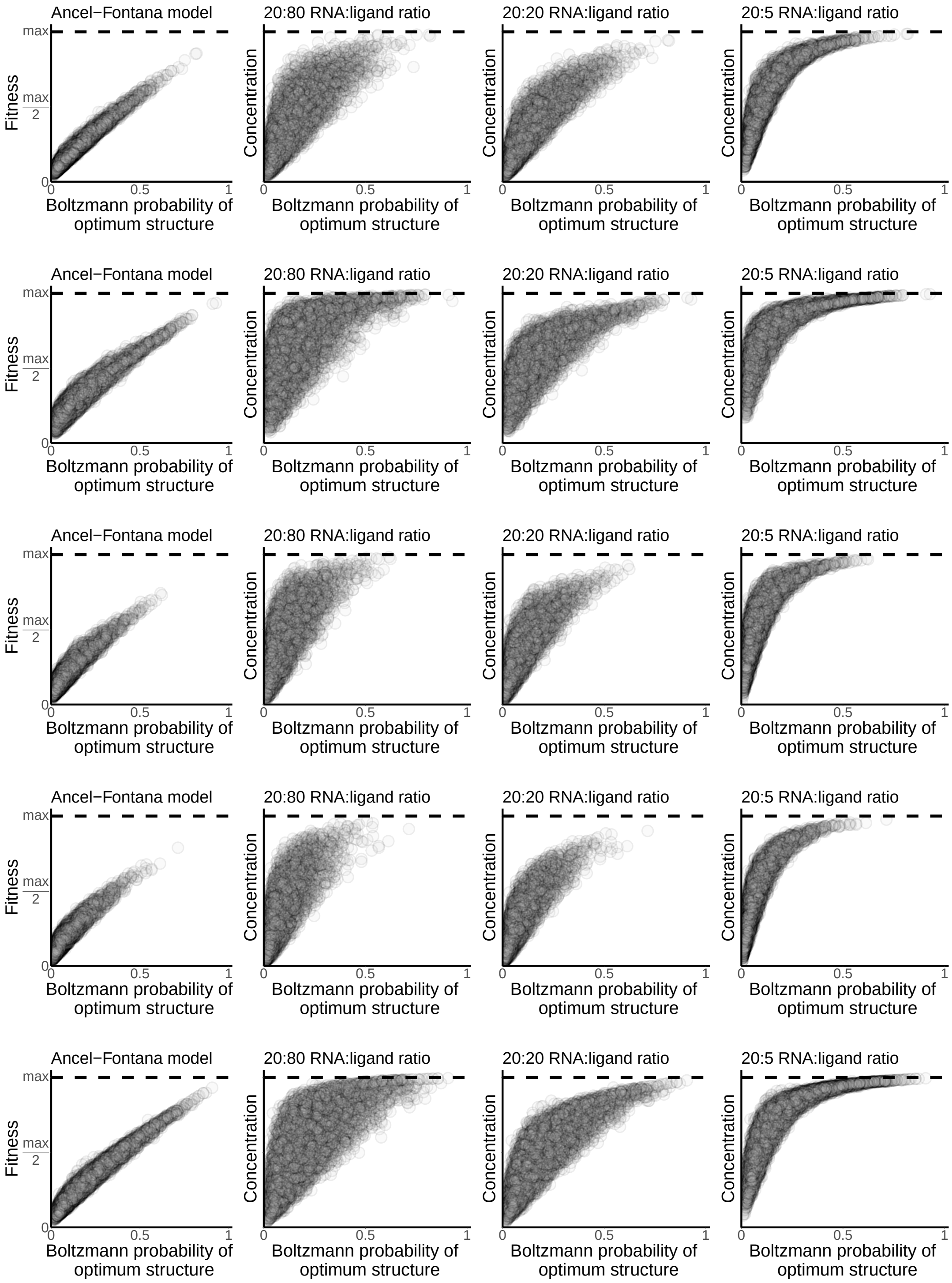


**Fig. SF6. Fitness as a function of thermodynamic stability in sequences that fold into a biological structure as their MFES when transitions between alternative structures are prohibited.** We assess thermodynamic stability as a sequence's Boltzmann probability for a specific biological structure. Each row presents data for sequences that fold into one structure. In order, the structures are: CPEB3 ribozyme, DsrA, Hepatitis ribozyme, snoRNA, tRNAphe.

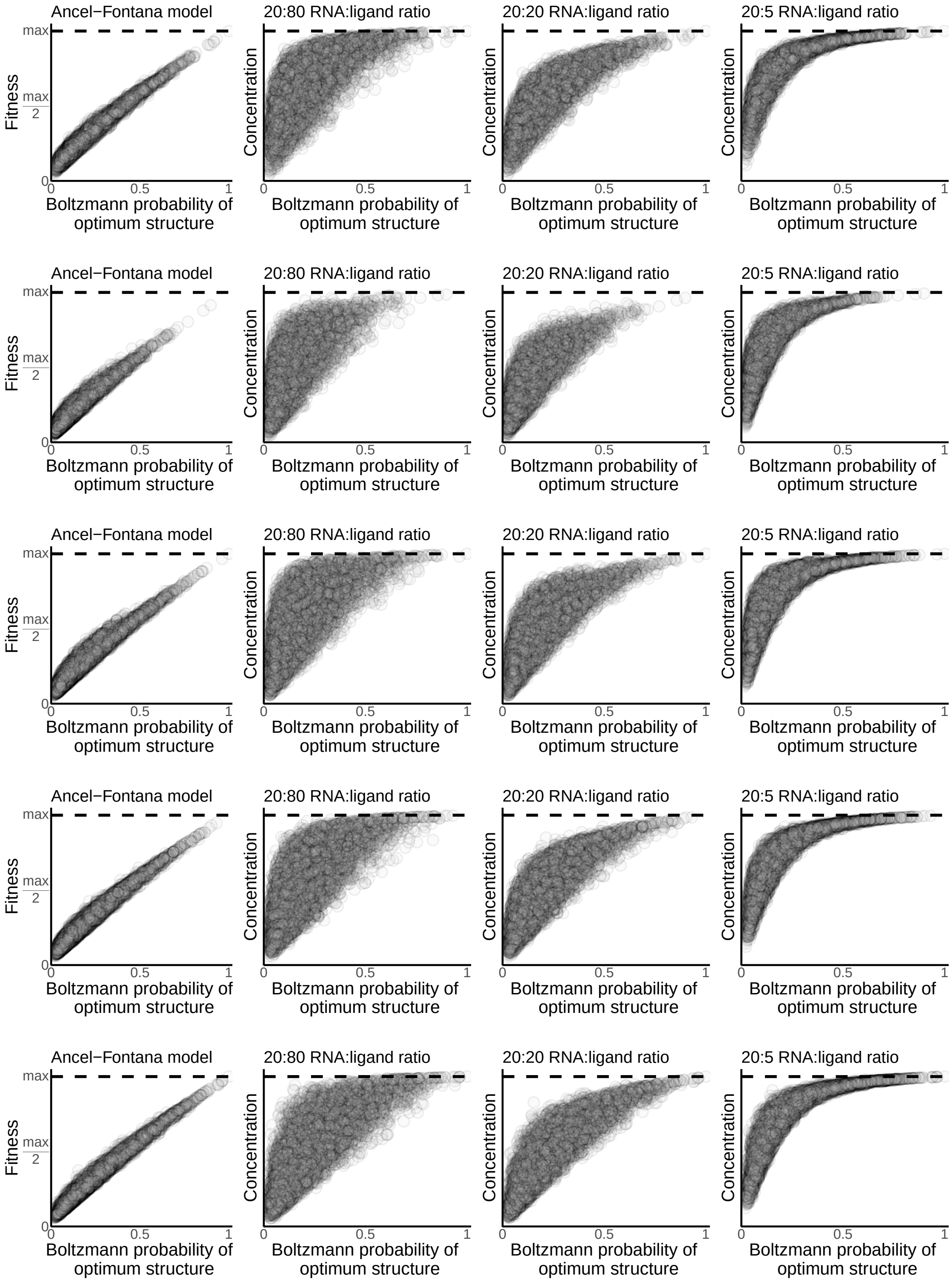
Ancel−Fontana model
20:80 RNA:ligand ratio
20:20 RNA:ligand ratio
20:5 RNA:ligand ratio
Fitness
Concentration
max
max/2
0
0.5
1
Boltzmann probability of optimum structure

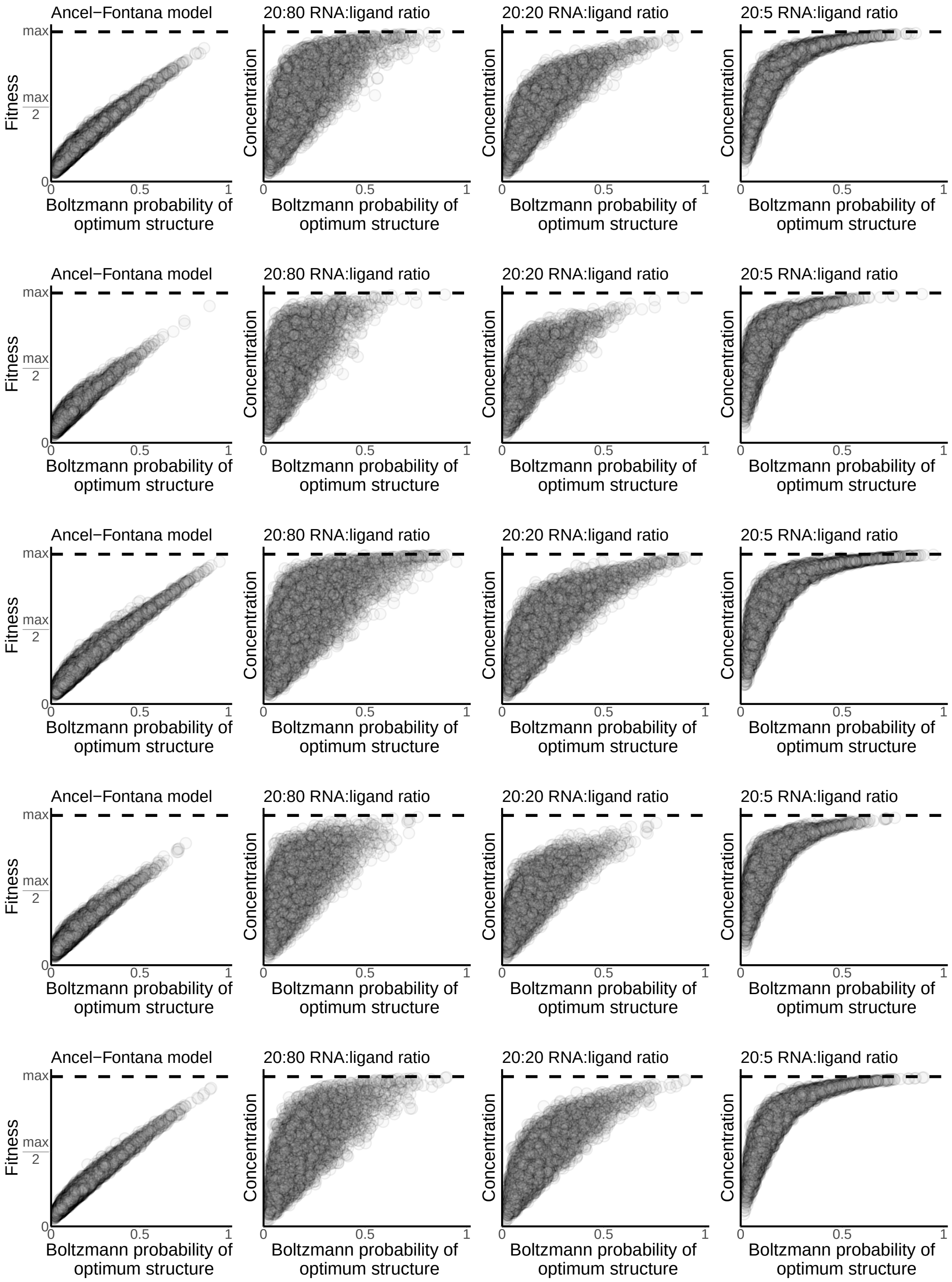


**Fig. SF7. Fitness as a function of thermodynamic stability in sequences that fold into pre-specified random structures as their MFES when transitions between alternative structures are prohibited.** We assess thermodynamic stability as a sequence's Boltzmann probability for a specific structure. Each row presents data for sequences that fold into one of the structures in Table ST2, appearing in the same order as in that table.

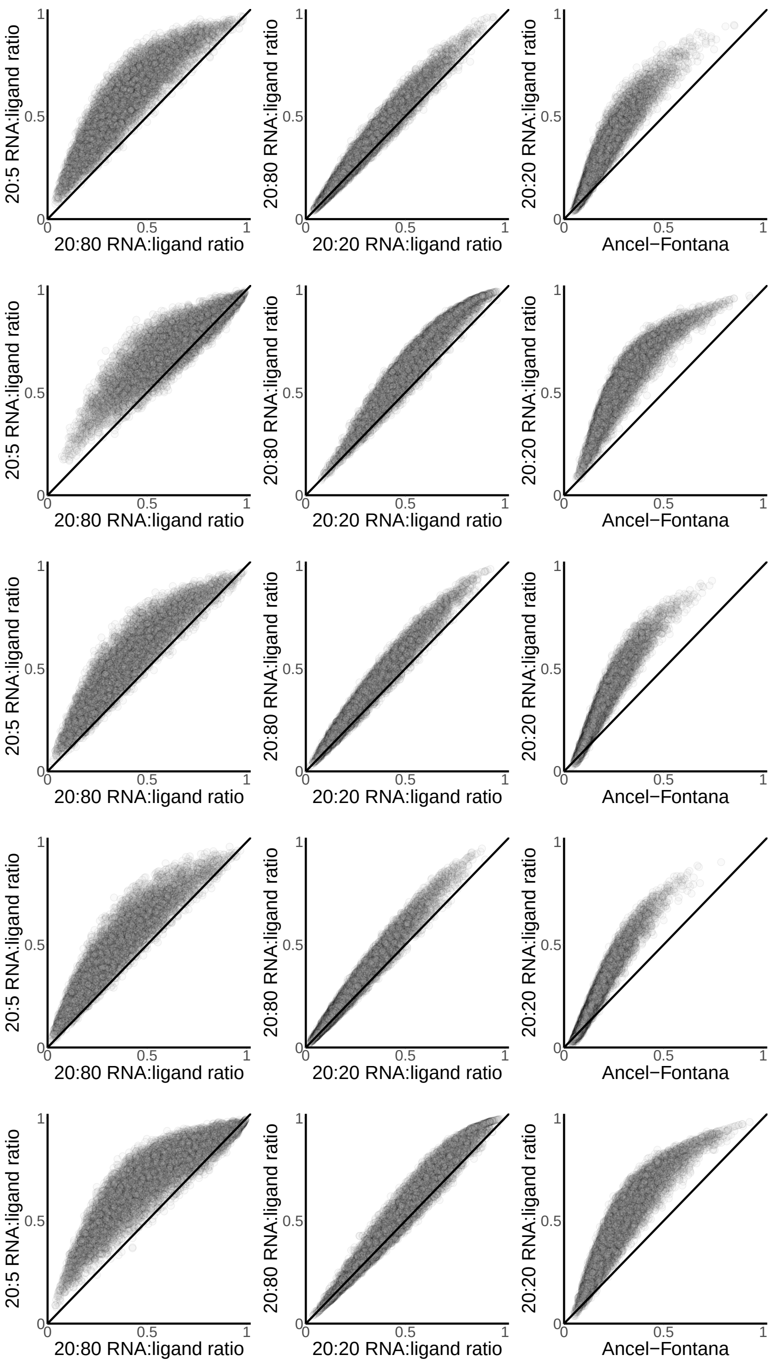


**Fig. SF8. Comparison of relative fitness of sequences that fold into specific biological structures under different models or ligand concentrations when transitions between alternative structures are prohibited.** In all panels, the diagonal indicates the identity line. Points above the diagonal refer to sequences with a higher fitness under the model/ligand concentration that corresponds to the vertical axis than under conditions described in the horizontal axis. Fitness values are normalized with respect to the maximum fitness value for the respective model. Each row presents data for sequences that fold into one structure. In order, the structures are: CPEB3 ribozyme, DsrA, Hepatitis ribozyme, snoRNA, tRNAphe.

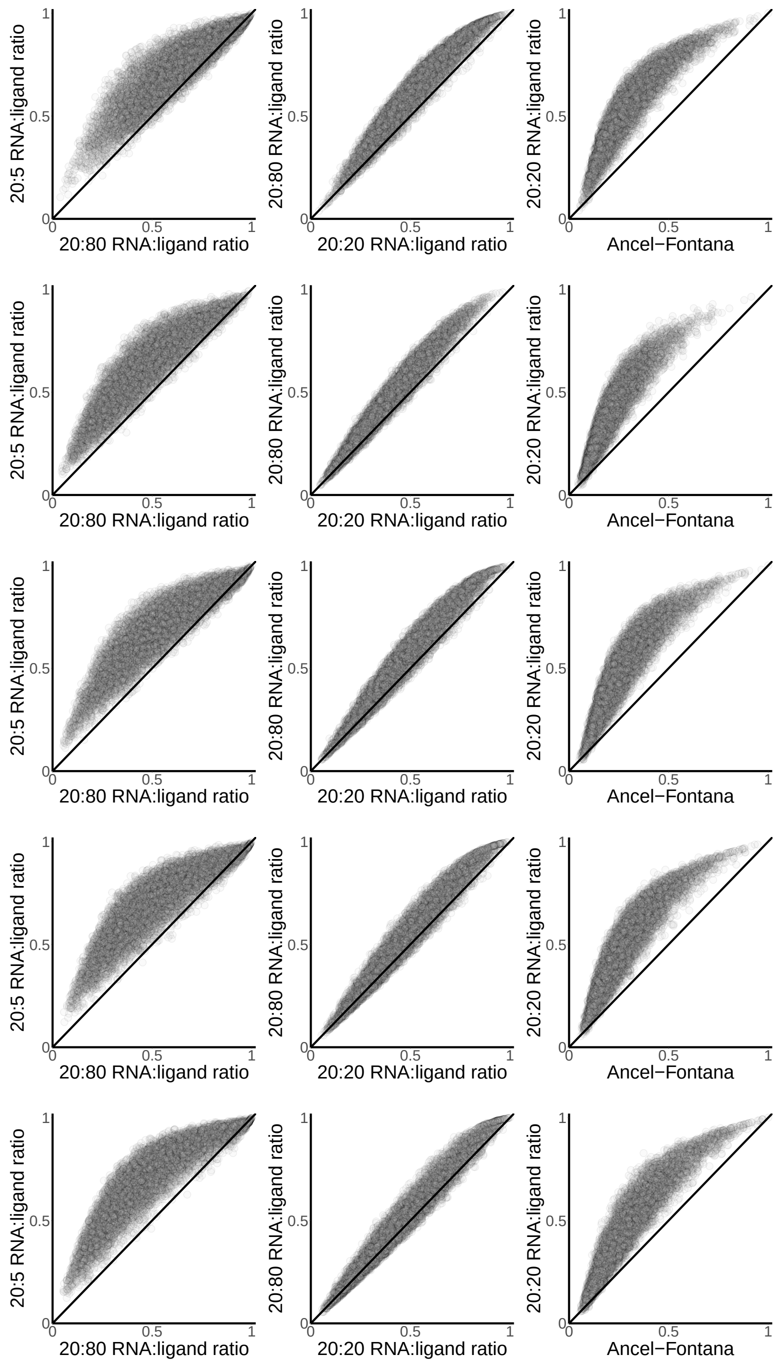
20:5 RNA:ligand ratio
20:80 RNA:ligand ratio
20:80 RNA:ligand ratio
20:20 RNA:ligand ratio
20:20 RNA:ligand ratio
Ancel−Fontana
0
0.5
1

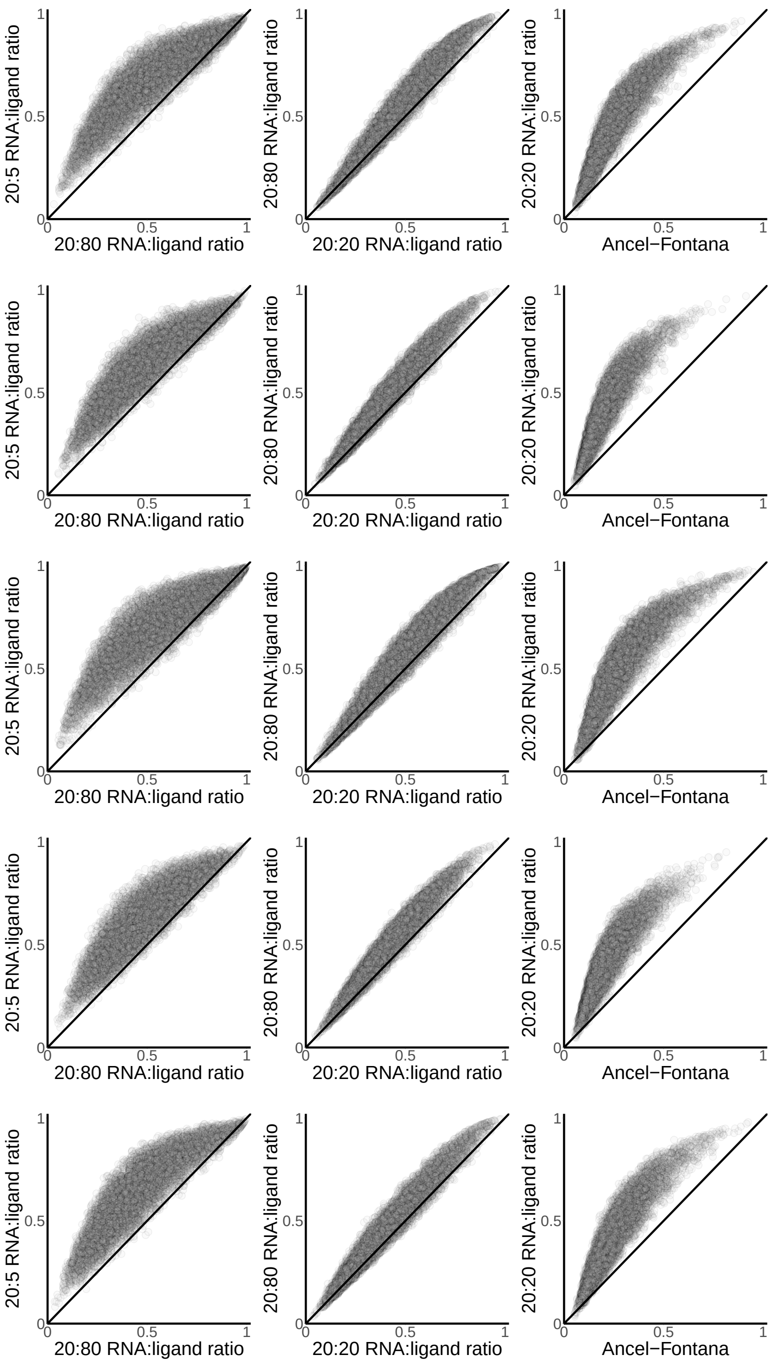


**Fig. SF9. Comparison of relative fitness of sequences that fold into specific random structures under different models or ligand concentrations when transitions between alternative structures are prohibited.** In all panels, the diagonal indicates the identity line. Points above the diagonal refer to sequences with a higher fitness under the model/ligand concentration that corresponds to the vertical axis than under conditions described in the horizontal axis. Fitness values are normalized with respect to the maximum fitness value for the respective model. Each row presents data for sequences that fold into one of the structures in Table ST2, appearing in the same order as in that table.

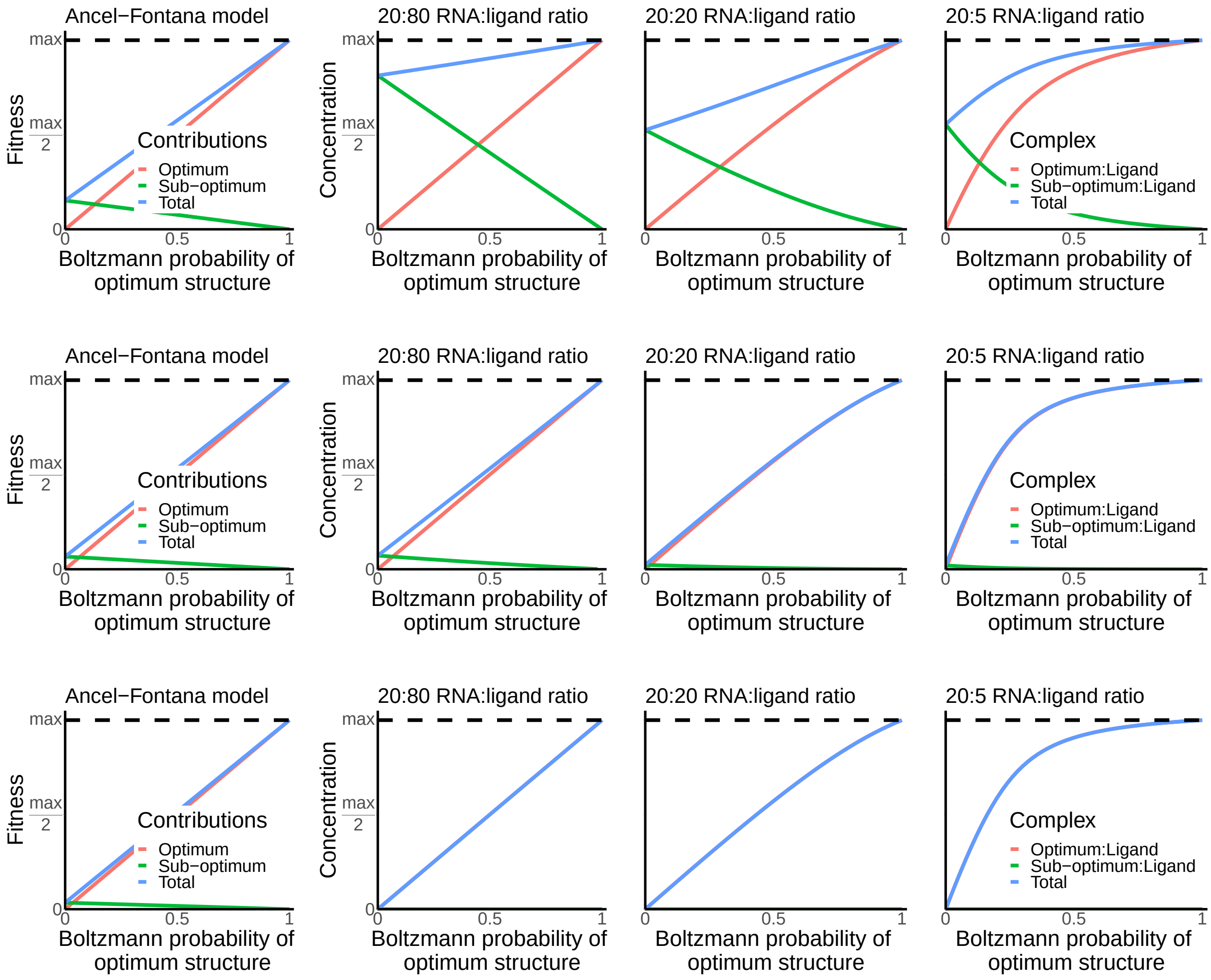


**Fig. SF10. Fitness as a function of thermodynamic stability of a sequence's MFES when transitions between alternative structures are prohibited.** We assume that all sequences have two different structures in their plastic repertoire: an optimal structure $\tau$ and a second structure $\upsilon$ at a fixed structural distance $d$ from $\tau$. Each row corresponds to a different fixed value of $d$: 8, 20, and 40 from top to bottom. Green and red lines indicate $\upsilon$'s and $\tau$'s contribution to fitness, respectively. The blue line refers to the sequence's total fitness.

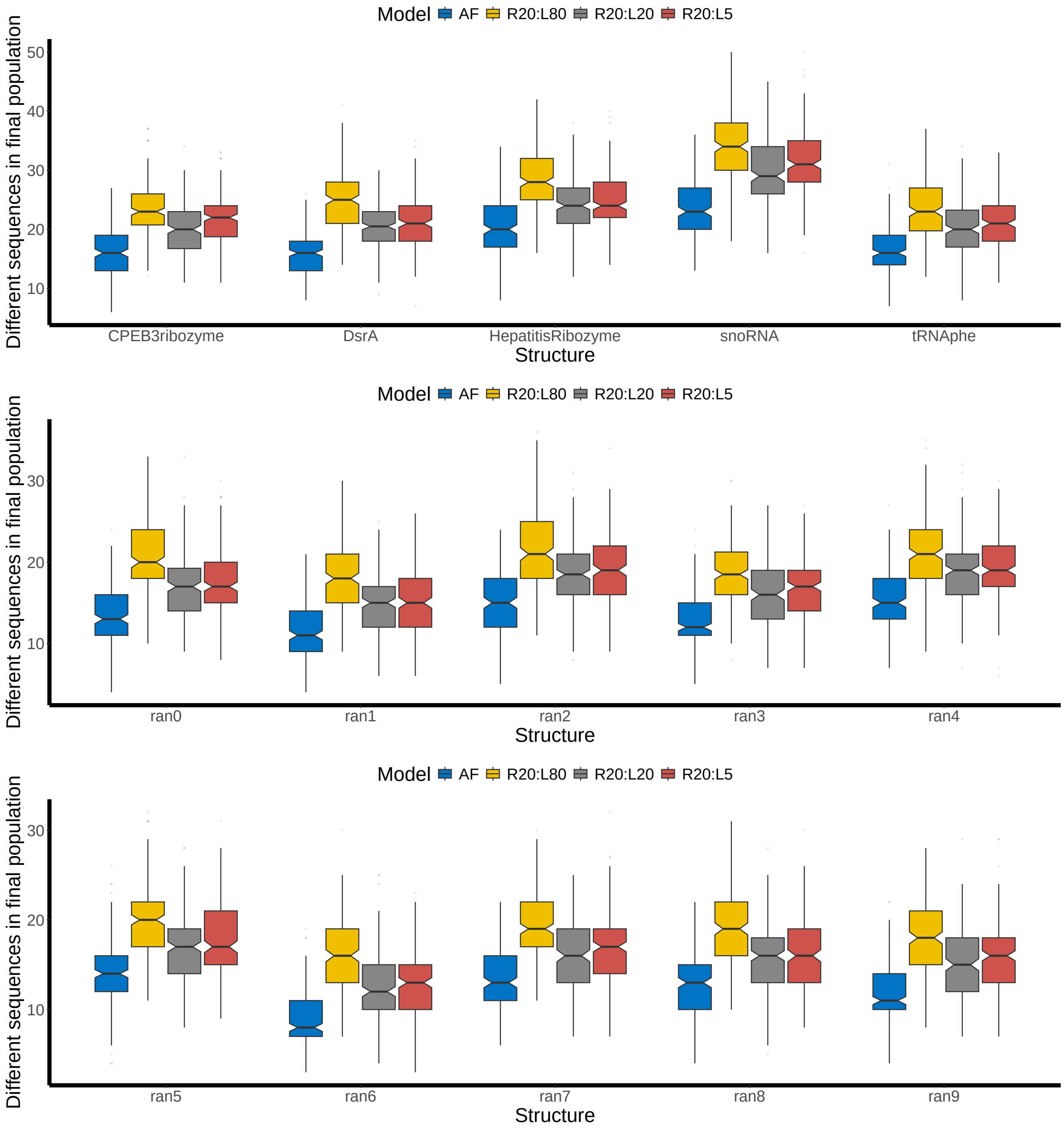


**Fig. SF11. Number of different sequences at the end of evolution.**

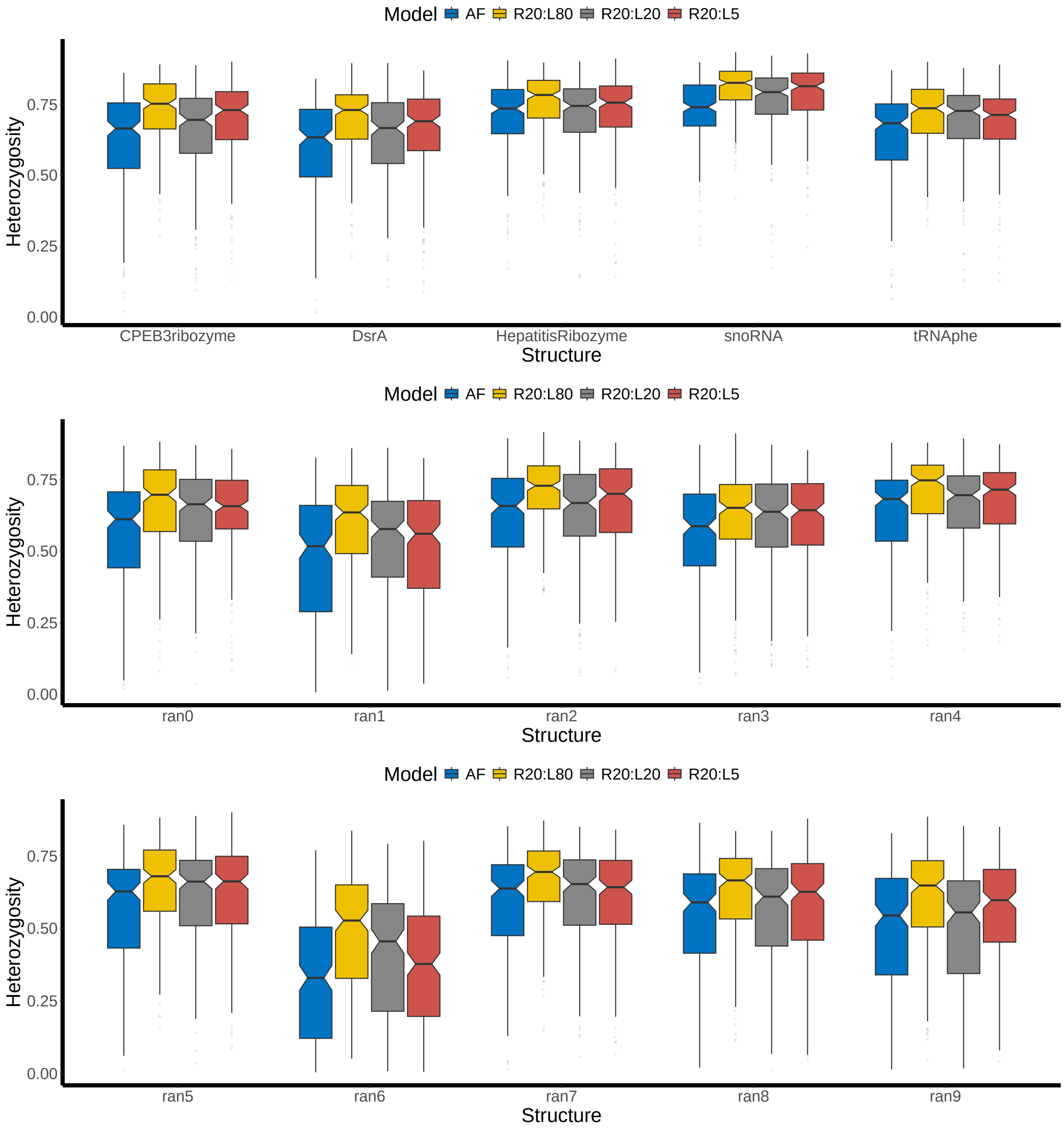


**Fig. SF12. Heterozygosity at the end of evolution.**

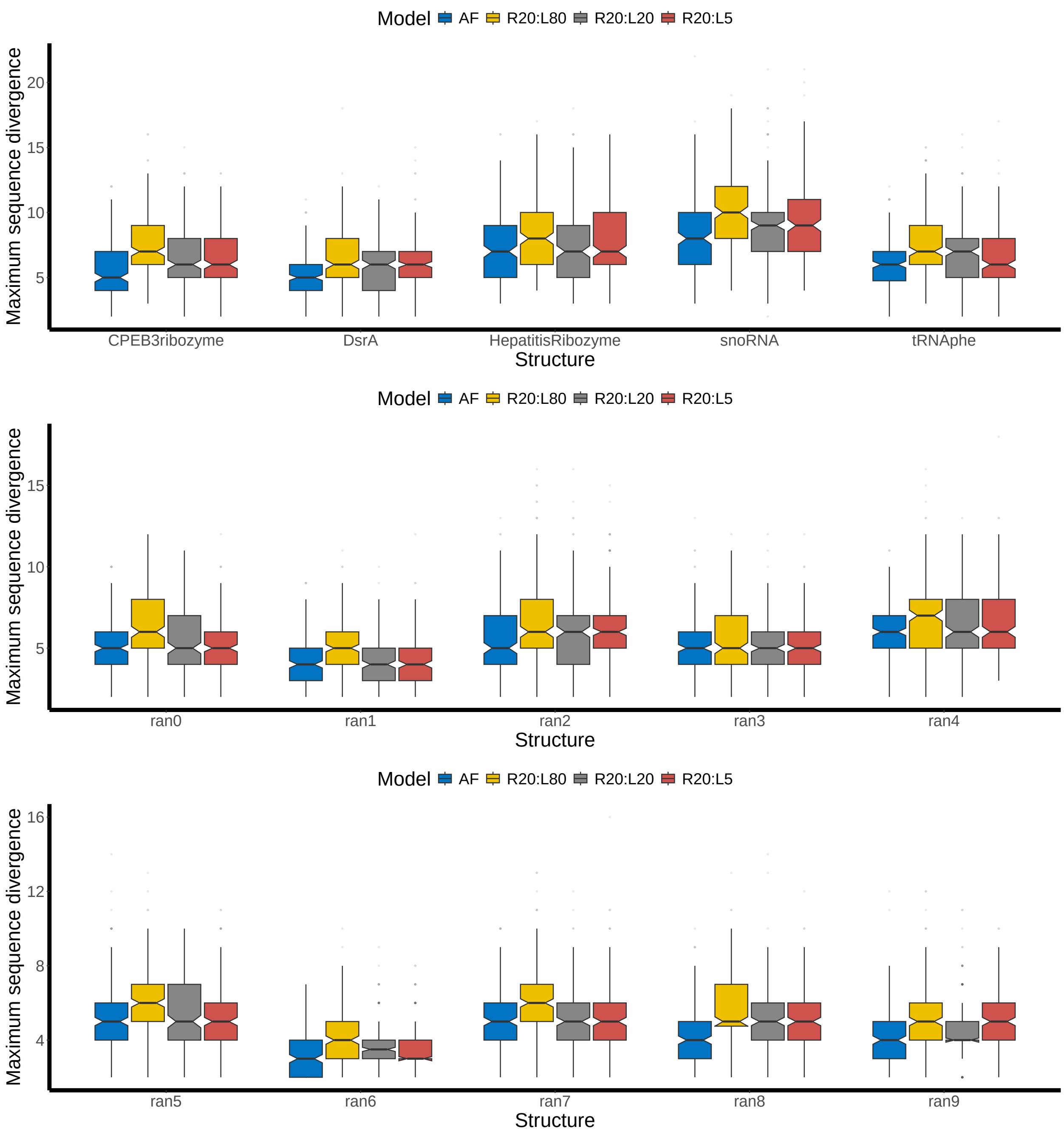


**Fig. SF13. Maximum sequence divergence at the end of evolution.**

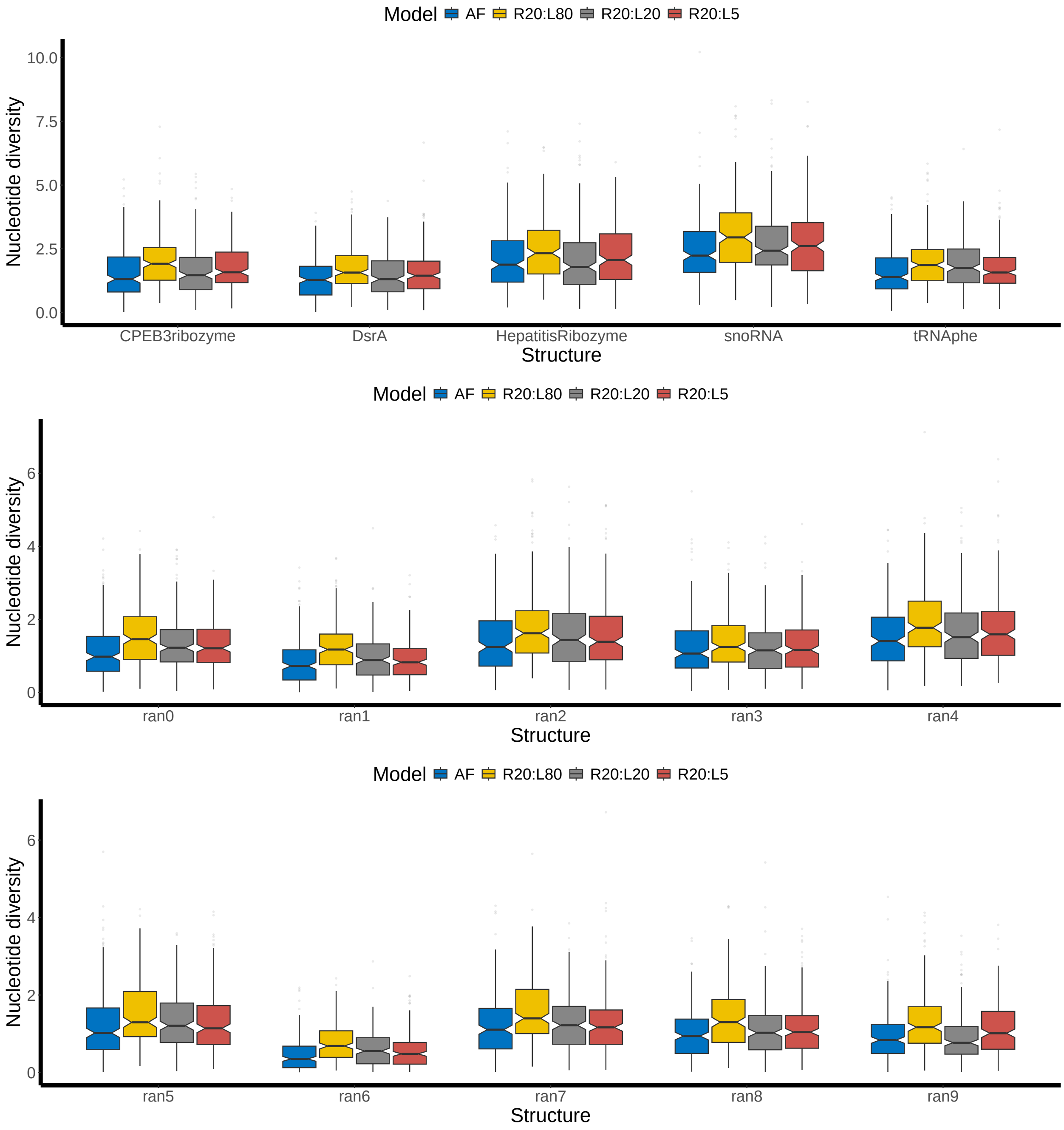


**Fig. SF14. Nucleotide diversity at the end of evolution.**

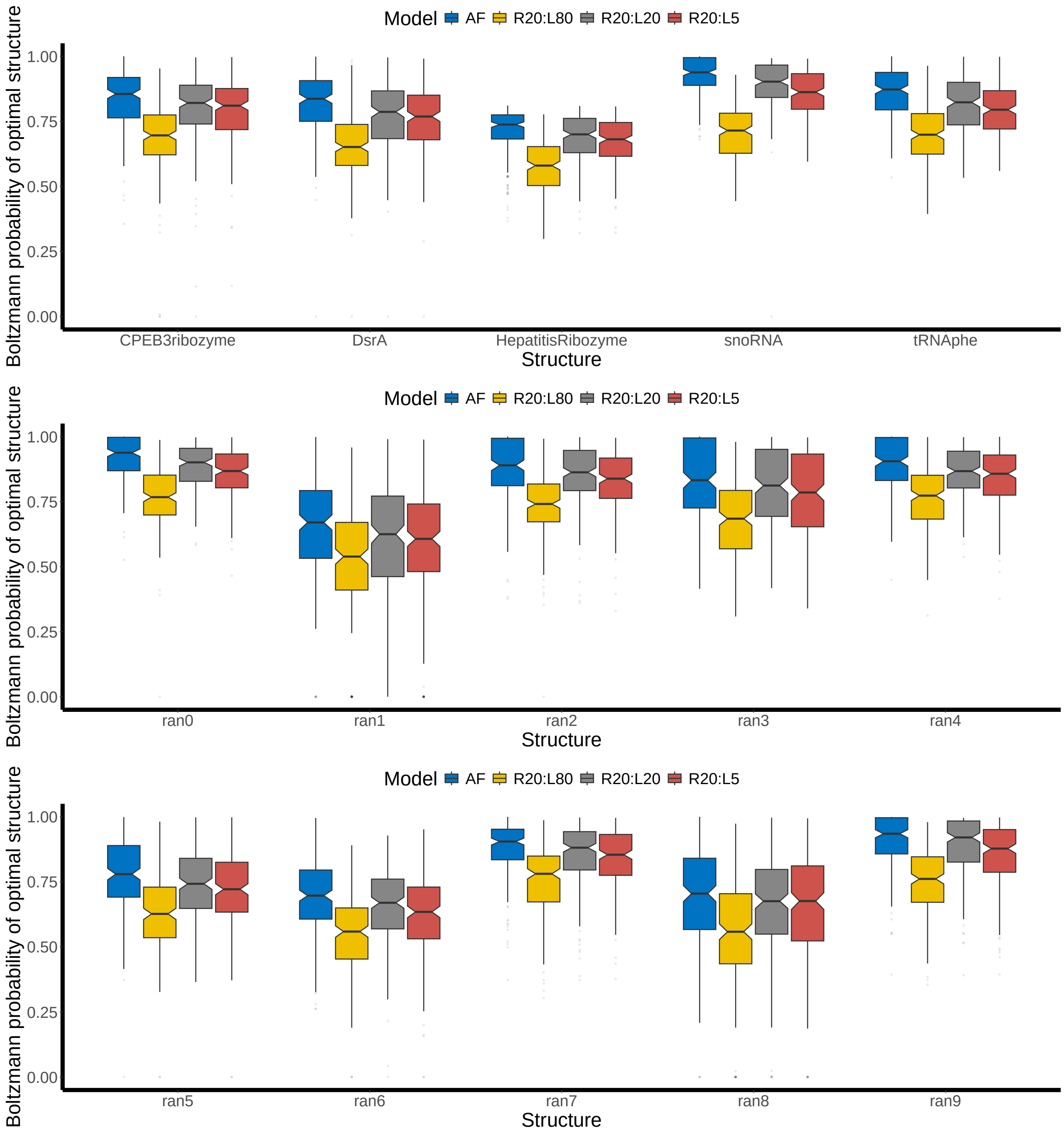


**Fig. SF15. Thermodynamic stability at the end of evolution.**

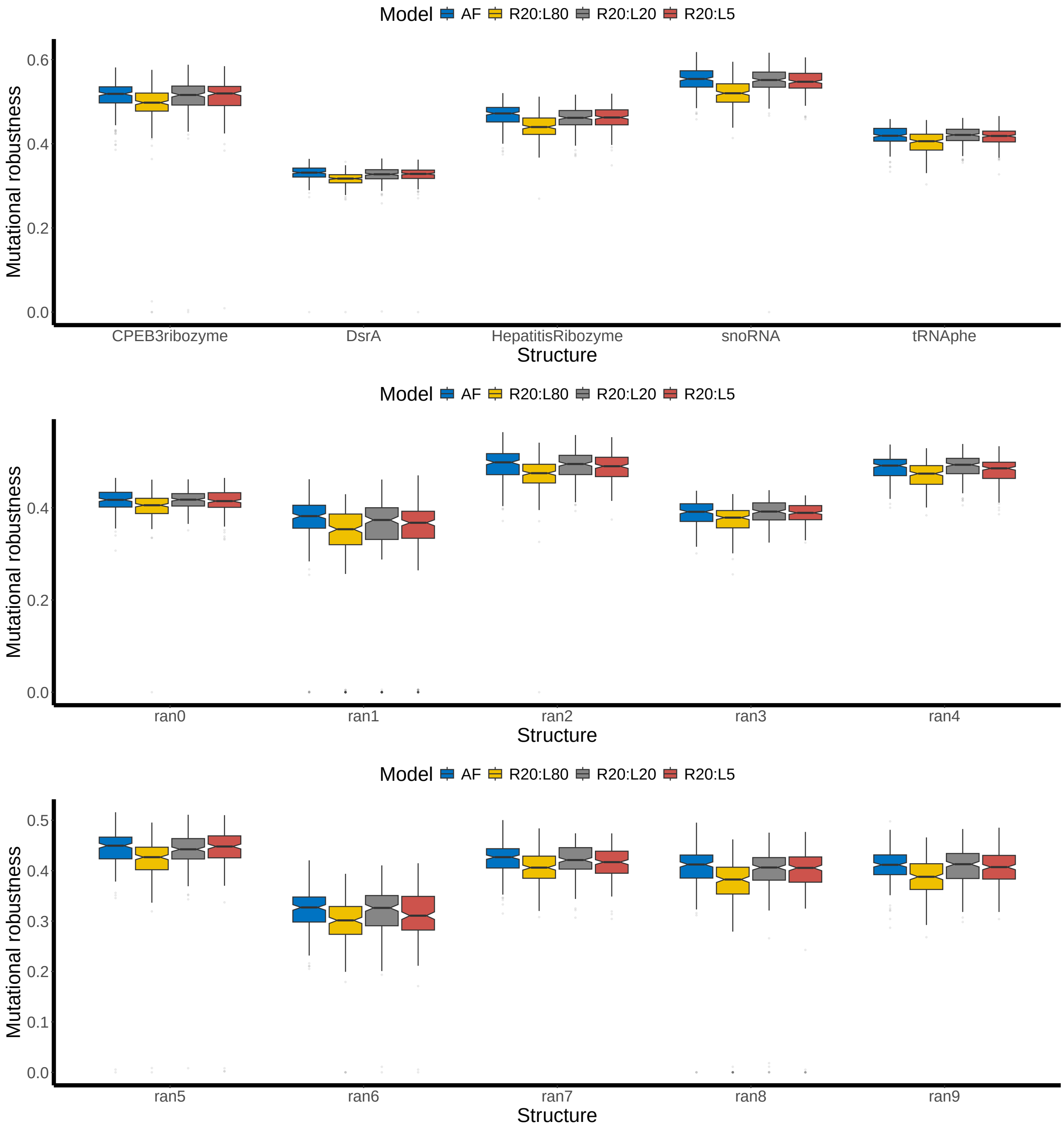


**Fig. SF16. Mutational robustness at the end of evolution.**

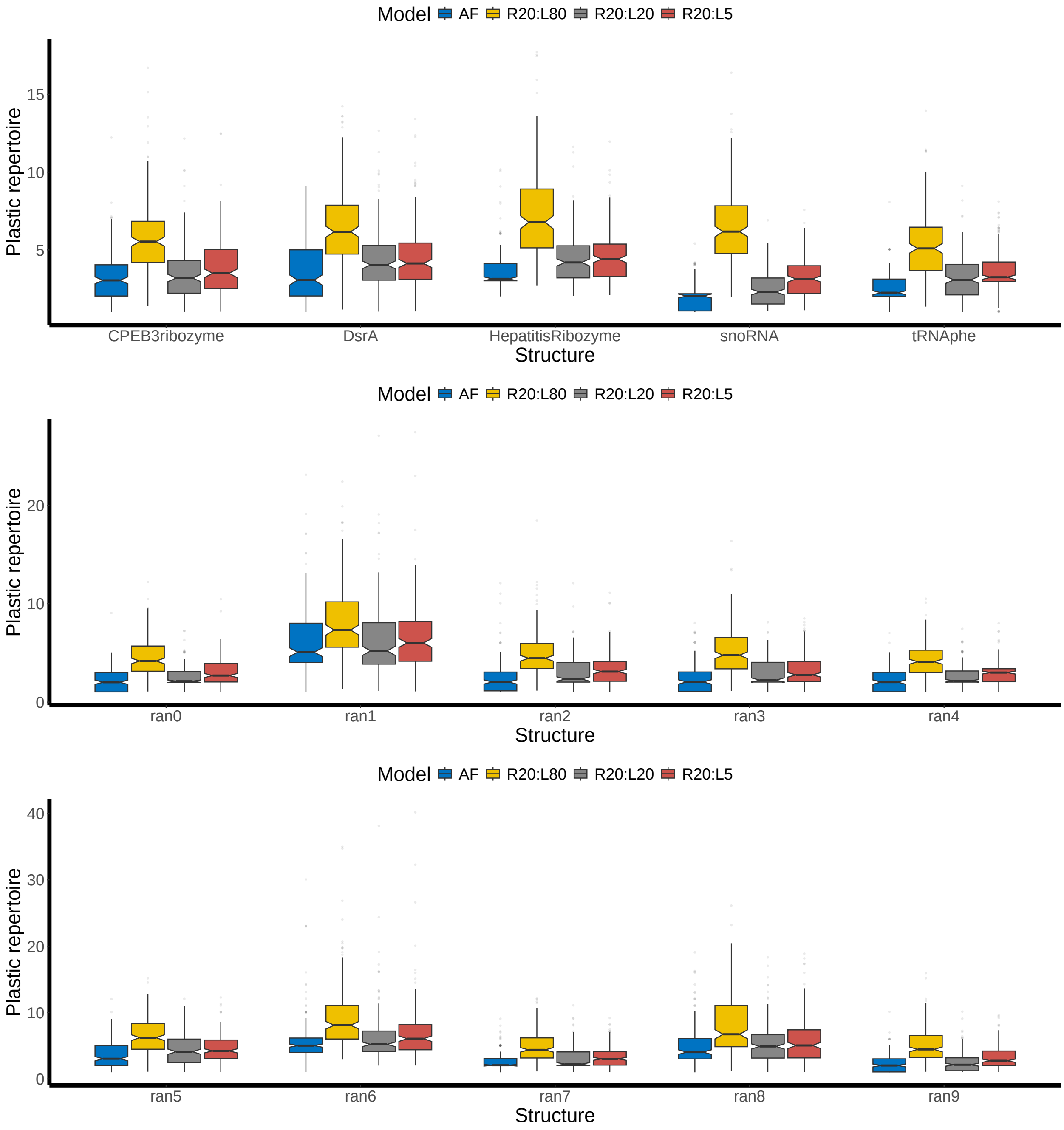


**Fig. SF17. Average plastic repertoire at the end of evolution.**

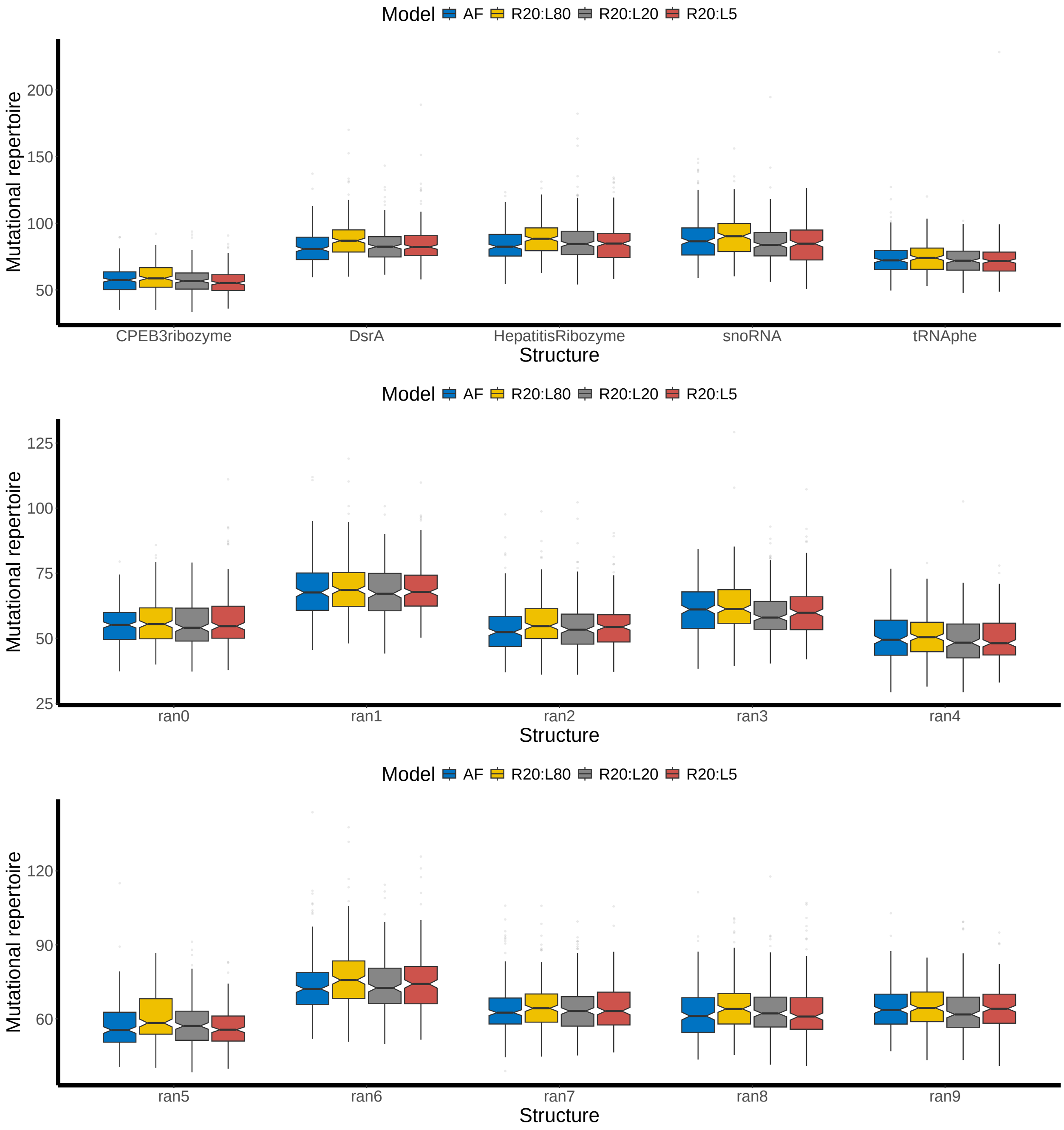


**Fig. SF18. Average mutational repertoire at the end of evolution.**

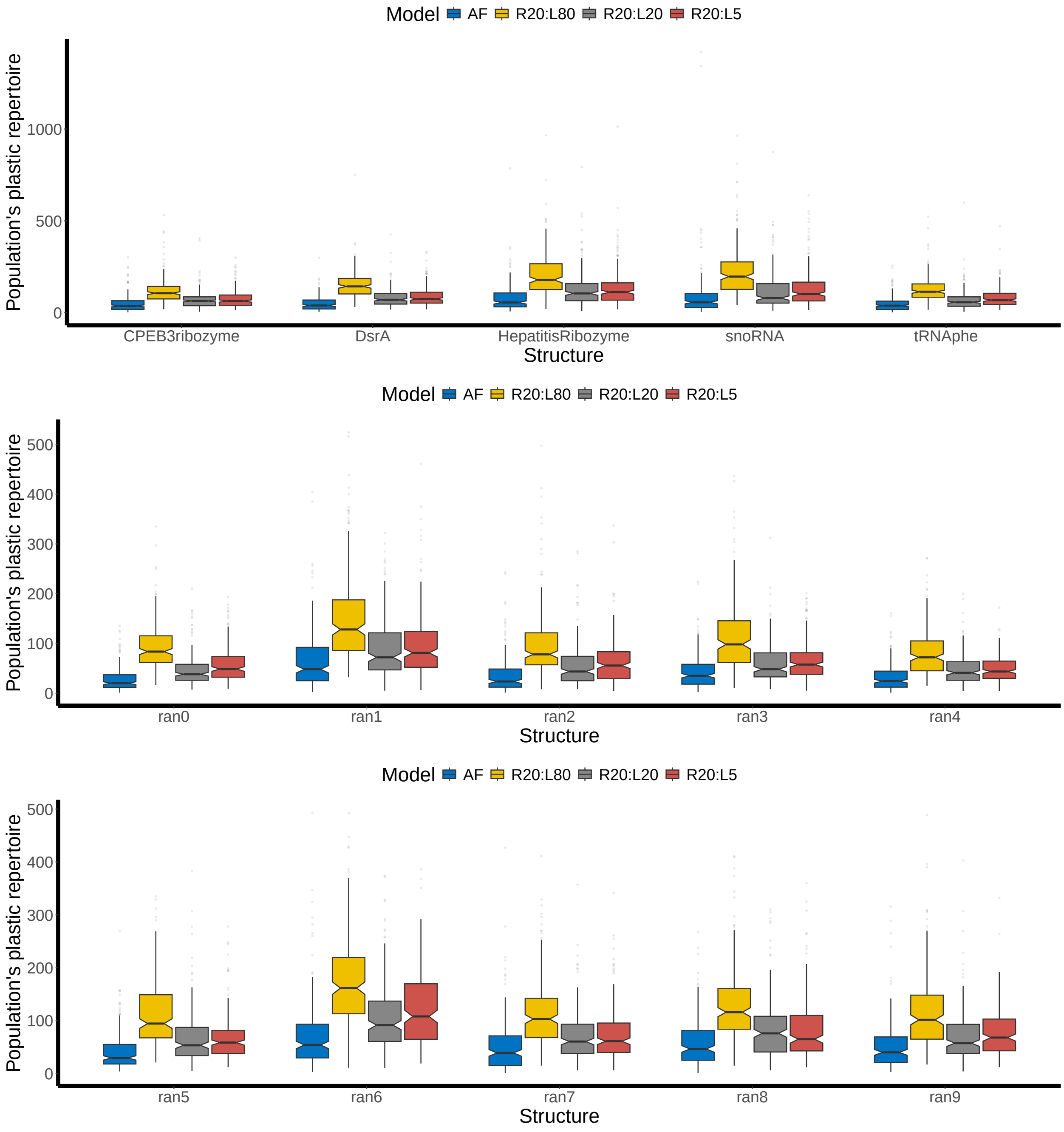


**Fig. SF19. Population's plastic repertoire at the end of evolution.**

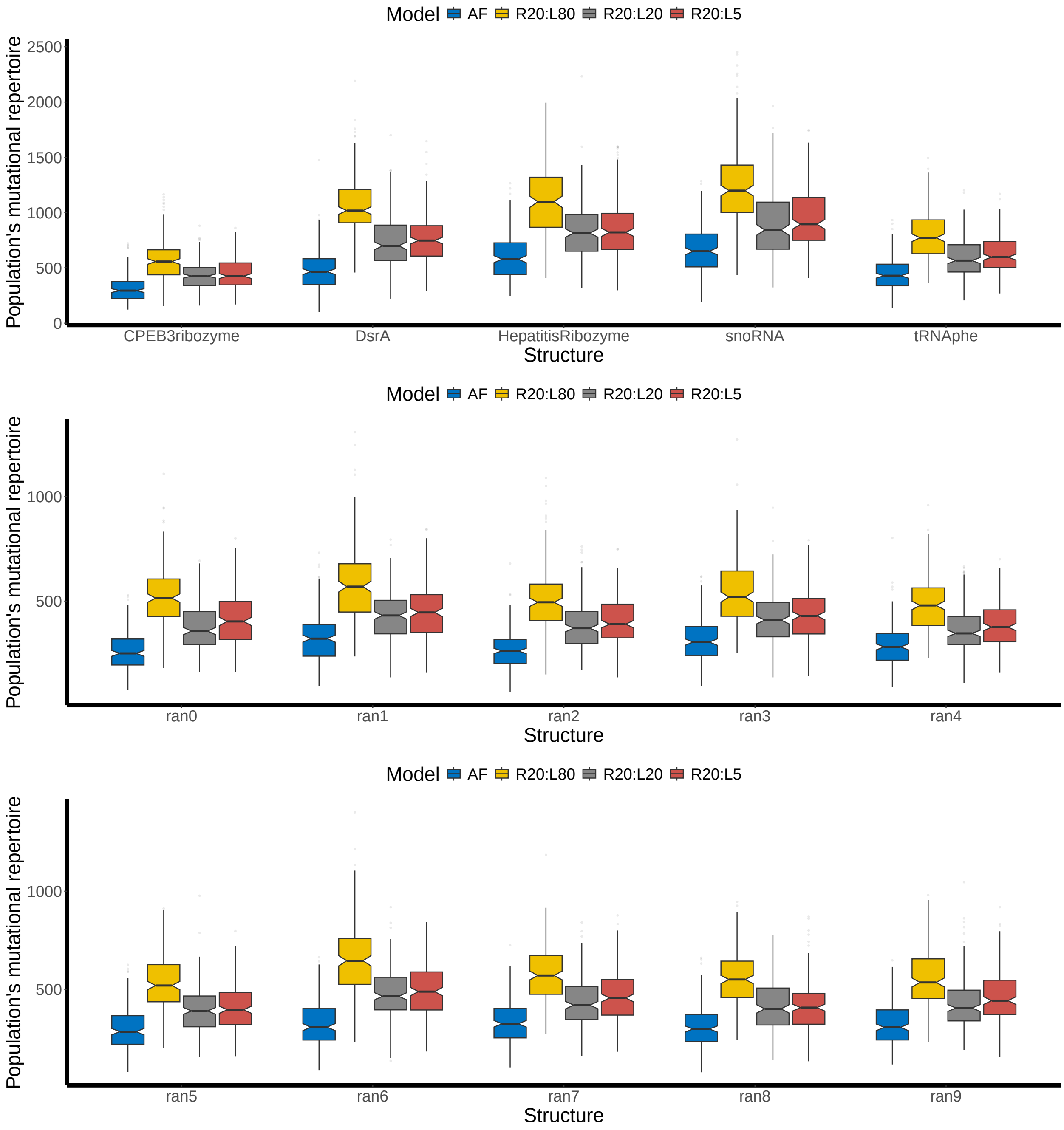


**Fig. SF20. Population's mutational repertoire at the end of evolution.**
