## Supplementary Tables for "RNA-ligand complexes and the attenuation of neutral confinement in the evolution of RNA secondary structures"

**Table ST1. Biological RNA structures considered in this work.**

| RNA | Species | Structure | Length |
|---|---|---|---|
| CPEB3 ribozyme | *Homo sapiens* | .(((((..........(((.......))).)))))...(((((..........)))))........... | 69 |
| DsrA RNA | *Salmonella enterica* subsp. enterica serovar Typhi str. Ty2 | .(((((((.....)))))))...(((((((((((((.....)))))))).)).))).((((((.((....))..)))))). | 81 |
| HDV ribozyme | Hepatitis D virus | ..(((((((.........(((........))))))))))......(((((((((((.(...).))))))).))))............. | 89 |
| Small nucleolar RNA, H/ACA box | *Macaca fascicularis* | .(((.(((............((((((...........))))))..........))).)))........(((((.............)))))....... | 101 |
| tRNA-Phe (GAA) | *Escherichia coli* str. K-12 substr. MG1655 | (((((((..((((........)))).(((((.......))))).....(((((.......)))))))))))). | 76 |

**Table ST2. Random RNA structures considered in this work.**

| Id | Structure | Length |
|---|---|---|
| Ran0 | ....((((....))))....((((((((....((.((......)).))..)))))))).. | 60 |
| Ran1 | ((((..((((((.(((((...........)))))......).))).))).)).))....... | 60 |
| Ran2 | .............((((.(.(((((.(((......))).)).))).).))))....... | 60 |
| Ran3 | .......(((.((((((.((((...)))).)))...))).)))..(((((.....))))) | 60 |
| Ran4 | ((((....))))........(((((((......))))))).((((......))))...... | 60 |
| Ran5 | ............((((...((((....((((((.....))))..))....))))..)))) | 60 |
| Ran6 | ..((((((((((.(((...((....(....)....))...))).....)))).)).)))) | 60 |
| Ran7 | .........((((((...((..(.((.((((..........)))))).)..)))))))). | 60 |
| Ran8 | ...(((.(((((((..............)))))...)))))..(((((......)).))) | 60 |
| Ran9 | ..((((....(((........(((((((..(....)..))))))))))......)))).. | 60 |

**Table ST3. Number of alternative structures in a sequence's plastic repertoire, considering an energy band of $D = 2$kcal mol$^{-1}$**

| Structure | Median | 1st quartile ($Q_1$) | 3rd quartile ($Q_3$) | QCD* | Mean±SD |
|---|---|---|---|---|---|
| CPEB3 ribozyme | 73 | 43 | 122 | 0.48 | $95.2 \pm 79.7$ |
| DsrA | 52 | 29 | 92 | 0.52 | $73 \pm 69.4$ |
| Hepatitis ribozyme | 124 | 70 | 220 | 0.52 | $177.4 \pm 183.1$ |
| snoRNA | 167 | 92 | 292 | 0.52 | $235.9 \pm 246$ |
| tRNAphe | 48 | 26 | 83 | 0.52 | $65.8 \pm 63.8$ |
| ran0 | 40 | 23 | 69 | 0.5 | $54.2 \pm 48.4$ |
| ran1 | 67 | 40 | 111 | 0.47 | $86.4 \pm 69.8$ |
| ran2 | 48 | 27 | 81 | 0.5 | $63.1 \pm 54.4$ |
| ran3 | 42 | 25 | 71 | 0.48 | $55.6 \pm 46.7$ |
| ran4 | 35 | 19 | 60 | 0.52 | $47.2 \pm 43.3$ |
| ran5 | 50 | 30 | 80 | 0.45 | $63.7 \pm 51.9$ |
| ran6 | 85 | 51 | 139 | 0.46 | $109.8 \pm 91.4$ |
| ran7 | 43 | 25 | 74 | 0.49 | $56.8 \pm 48.7$ |
| ran8 | 69 | 41 | 114 | 0.47 | $89.2 \pm 72.4$ |
| ran9 | 52 | 31 | 85 | 0.47 | $65.7 \pm 51.5$ |

* Quartile coefficient of dispersion: $\frac{Q_3 - Q_1}{Q_3 + Q_1}$

**Table ST4. Two-way ANOVA for the effect of structure and model on different sequences in final population.** Repeated measures under different models.

| Effect | DFn | DFd | $F$ | $p$ | $p < 0.05$ | G. $\eta^2$ |
|---|---|---|---|---|---|---|
| Structure | 14 | 2985 | 487.7 | 0 | * | 0.49 |
| Model | 2.95 | 8801.5 | 1853 | 0 | * | 0.265 |
| Structure:Model | 41.28 | 8801.5 | 5.059 | 6.65e-24 | * | 0.014 |

**Table ST5. One-way ANOVA for the effect of model on different sequences in final population.** Repeated measures under different models.

| Structure | DFn | DFd | $F$ | $p$ | $p < 0.05$ | G. $\eta^2$ | Adjusted $p$ |
|---|---|---|---|---|---|---|---|
| CPEB3ribozyme | 3 | 597 | 124.2 | 1.68e-62 | * | 0.269 | 3.6e-62 |
| DsrA | 2.84 | 564.78 | 192.5 | 8.99e-83 | * | 0.359 | 6.742e-82 |
| HepatitisRibozyme | 3 | 597 | 111.4 | 2.77e-57 | * | 0.253 | 4.617e-57 |
| ran0 | 3 | 597 | 129.8 | 9.54e-65 | * | 0.291 | 2.862e-64 |
| ran1 | 3 | 597 | 150 | 1.97e-72 | * | 0.302 | 7.387e-72 |
| ran2 | 3 | 597 | 107.6 | 1.05e-55 | * | 0.235 | 1.432e-55 |
| ran3 | 3 | 597 | 102.9 | 1.06e-53 | * | 0.245 | 1.325e-53 |
| ran4 | 3 | 597 | 88.17 | 3.09e-47 | * | 0.202 | 3.09e-47 |
| ran5 | 3 | 597 | 98.06 | 1.3e-51 | * | 0.222 | 1.393e-51 |
| ran6 | 2.87 | 571.76 | 208.3 | 1.19e-88 | * | 0.337 | 1.785e-87 |
| ran7 | 3 | 597 | 108.3 | 5.61e-56 | * | 0.244 | 8.415e-56 |
| ran8 | 3 | 597 | 114.8 | 1.11e-58 | * | 0.239 | 2.081e-58 |
| ran9 | 2.87 | 571.62 | 131.1 | 1.4e-62 | * | 0.273 | 3.5e-62 |
| snoRNA | 2.89 | 575.06 | 166.8 | 9.24e-76 | * | 0.33 | 4.62e-75 |
| tRNAphe | 3 | 597 | 100.5 | 1.13e-52 | * | 0.223 | 1.304e-52 |

**Table ST6. Pair-wise post hoc tests for the effect of structure and model on different sequences in final population.** Repeated measures under different models.

| Structure | Model M1 | Model M2 | statistic | $p$ | Adjusted $p$ | Signif. | Mean M1 | Mean M2 | Difference |
|---|---|---|---|---|---|---|---|---|---|
| CPEB3 ribozyme | AF | R20:L20 | -11.31 | 3.2e-23 | 6.4e-23 | **** | 16.02 | 20.05 | -4.035 |
| CPEB3 ribozyme | AF | R20:L80 | -17.57 | 2.48e-42 | 1.49e-41 | **** | 16.02 | 23.35 | -7.335 |
| CPEB3 ribozyme | AF | R20:L5 | -14.52 | 4.75e-33 | 1.43e-32 | **** | 16.02 | 21.58 | -5.57 |
| CPEB3 ribozyme | R20:L20 | R20:L80 | -8.18 | 3.32e-14 | 4.98e-14 | **** | 20.05 | 23.35 | -3.3 |
| CPEB3 ribozyme | R20:L20 | R20:L5 | -3.794 | 0.000197 | 0.000197 | *** | 20.05 | 21.58 | -1.535 |
| CPEB3 ribozyme | R20:L80 | R20:L5 | 4.271 | 3.01e-05 | 3.61e-05 | **** | 23.35 | 21.58 | 1.765 |
| DsrA | AF | R20:L20 | -14.16 | 6.24e-32 | 1.25e-31 | **** | 15.88 | 20.56 | -4.685 |
| DsrA | AF | R20:L80 | -24.45 | 7.18e-62 | 4.31e-61 | **** | 15.88 | 24.76 | -8.885 |
| DsrA | AF | R20:L5 | -15.24 | 2.98e-35 | 8.94e-35 | **** | 15.88 | 21.21 | -5.33 |
| DsrA | R20:L20 | R20:L80 | -10.11 | 1.17e-19 | 1.76e-19 | **** | 20.56 | 24.76 | -4.2 |
| DsrA | R20:L20 | R20:L5 | -1.785 | 0.076 | 0.076 | ns | 20.56 | 21.21 | -0.645 |
| DsrA | R20:L80 | R20:L5 | 8.78 | 7.52e-16 | 9.02e-16 | **** | 24.76 | 21.21 | 3.555 |
| Hepatitis ribozyme | AF | R20:L20 | -9.344 | 1.93e-17 | 3.86e-17 | **** | 20.29 | 24.32 | -4.03 |
| Hepatitis ribozyme | AF | R20:L80 | -17.74 | 7.31e-43 | 4.39e-42 | **** | 20.29 | 28.17 | -7.88 |
| Hepatitis ribozyme | AF | R20:L5 | -11.8 | 1.03e-24 | 3.09e-24 | **** | 20.29 | 24.98 | -4.69 |
| Hepatitis ribozyme | R20:L20 | R20:L80 | -8.7 | 1.25e-15 | 1.88e-15 | **** | 24.32 | 28.17 | -3.85 |
| Hepatitis ribozyme | R20:L20 | R20:L5 | -1.612 | 0.108 | 0.108 | ns | 24.32 | 24.98 | -0.66 |
| Hepatitis ribozyme | R20:L80 | R20:L5 | 6.732 | 1.75e-10 | 2.1e-10 | **** | 28.17 | 24.98 | 3.19 |
| ran0 | AF | R20:L20 | -10.46 | 1.1e-20 | 2.2e-20 | **** | 13.38 | 17.07 | -3.69 |
| ran0 | AF | R20:L80 | -19.93 | 2.58e-49 | 1.55e-48 | **** | 13.38 | 20.56 | -7.18 |
| ran0 | AF | R20:L5 | -12.46 | 1.02e-26 | 3.06e-26 | **** | 13.38 | 17.55 | -4.175 |
| ran0 | R20:L20 | R20:L80 | -8.7 | 1.25e-15 | 1.88e-15 | **** | 17.07 | 20.56 | -3.49 |
| ran0 | R20:L20 | R20:L5 | -1.36 | 0.175 | 0.175 | ns | 17.07 | 17.55 | -0.485 |
| ran0 | R20:L80 | R20:L5 | 7.844 | 2.61e-13 | 3.13e-13 | **** | 20.56 | 17.55 | 3.005 |
| ran1 | AF | R20:L20 | -10.85 | 7.38e-22 | 1.11e-21 | **** | 11.53 | 14.52 | -2.99 |
| ran1 | AF | R20:L80 | -20.74 | 1.26e-51 | 7.56e-51 | **** | 11.53 | 18.04 | -6.505 |
| ran1 | AF | R20:L5 | -11.31 | 3.22e-23 | 9.66e-23 | **** | 11.53 | 14.97 | -3.44 |
| ran1 | R20:L20 | R20:L80 | -10.96 | 3.45e-22 | 6.9e-22 | **** | 14.52 | 18.04 | -3.515 |
| ran1 | R20:L20 | R20:L5 | -1.482 | 0.14 | 0.14 | ns | 14.52 | 14.97 | -0.45 |
| ran1 | R20:L80 | R20:L5 | 9.446 | 9.85e-18 | 1.18e-17 | **** | 18.04 | 14.97 | 3.065 |
| ran2 | AF | R20:L20 | -10.36 | 2.1e-20 | 4.2e-20 | **** | 14.88 | 18.66 | -3.785 |
| ran2 | AF | R20:L80 | -17.74 | 7.45e-43 | 4.47e-42 | **** | 14.88 | 21.55 | -6.675 |
| ran2 | AF | R20:L5 | -12.19 | 6.87e-26 | 2.06e-25 | **** | 14.88 | 19.32 | -4.44 |
| ran2 | R20:L20 | R20:L80 | -7.475 | 2.4e-12 | 3.6e-12 | **** | 18.66 | 21.55 | -2.89 |
| ran2 | R20:L20 | R20:L5 | -1.706 | 0.09 | 0.09 | ns | 18.66 | 19.32 | -0.655 |
| ran2 | R20:L80 | R20:L5 | 5.7 | 4.28e-08 | 5.14e-08 | **** | 21.55 | 19.32 | 2.235 |
| ran3 | AF | R20:L20 | -10.73 | 1.78e-21 | 3.56e-21 | **** | 12.78 | 16.3 | -3.515 |
| ran3 | AF | R20:L80 | -18.34 | 1.25e-44 | 7.5e-44 | **** | 12.78 | 18.8 | -6.02 |
| ran3 | AF | R20:L5 | -11.14 | 1e-22 | 3e-22 | **** | 12.78 | 16.66 | -3.875 |
| ran3 | R20:L20 | R20:L80 | -6.707 | 2.01e-10 | 3.02e-10 | **** | 16.3 | 18.8 | -2.505 |
| ran3 | R20:L20 | R20:L5 | -1.025 | 0.307 | 0.307 | ns | 16.3 | 16.66 | -0.36 |
| ran3 | R20:L80 | R20:L5 | 6.039 | 7.48e-09 | 8.98e-09 | **** | 18.8 | 16.66 | 2.145 |

| Structure | Model M1 | Model M2 | statistic | $p$ | Adjusted $p$ | Signif. | Mean M1 | Mean M2 | Difference |
|---|---|---|---|---|---|---|---|---|---|
| ran4 | AF | R20:L20 | -10.02 | 2.22e-19 | 4.44e-19 | **** | 15.49 | 18.93 | -3.44 |
| ran4 | AF | R20:L80 | -15.29 | 2e-35 | 1.2e-34 | **** | 15.49 | 21.26 | -5.77 |
| ran4 | AF | R20:L5 | -11.29 | 3.76e-23 | 1.13e-22 | **** | 15.49 | 19.19 | -3.7 |
| ran4 | R20:L20 | R20:L80 | -6.234 | 2.67e-09 | 4e-09 | **** | 18.93 | 21.26 | -2.33 |
| ran4 | R20:L20 | R20:L5 | -0.7564 | 0.45 | 0.45 | ns | 18.93 | 19.19 | -0.26 |
| ran4 | R20:L80 | R20:L5 | 5.303 | 3.02e-07 | 3.62e-07 | **** | 21.26 | 19.19 | 2.07 |
| ran5 | AF | R20:L20 | -9.158 | 6.52e-17 | 1.3e-16 | **** | 13.98 | 16.87 | -2.89 |
| ran5 | AF | R20:L80 | -16.77 | 6.15e-40 | 3.69e-39 | **** | 13.98 | 19.85 | -5.87 |
| ran5 | AF | R20:L5 | -10.5 | 8.57e-21 | 2.57e-20 | **** | 13.98 | 17.8 | -3.82 |
| ran5 | R20:L20 | R20:L80 | -8.65 | 1.72e-15 | 2.58e-15 | **** | 16.87 | 19.85 | -2.98 |
| ran5 | R20:L20 | R20:L5 | -2.624 | 0.009 | 0.009 | ** | 16.87 | 17.8 | -0.93 |
| ran5 | R20:L80 | R20:L5 | 5.713 | 4.01e-08 | 4.81e-08 | **** | 19.85 | 17.8 | 2.05 |
| ran6 | AF | R20:L20 | -14.11 | 8.67e-32 | 1.73e-31 | **** | 8.86 | 12.58 | -3.725 |
| ran6 | AF | R20:L80 | -24.43 | 8.38e-62 | 5.03e-61 | **** | 8.86 | 15.98 | -7.125 |
| ran6 | AF | R20:L5 | -16.08 | 7.97e-38 | 2.39e-37 | **** | 8.86 | 12.9 | -4.035 |
| ran6 | R20:L20 | R20:L80 | -11.28 | 3.91e-23 | 5.86e-23 | **** | 12.58 | 15.98 | -3.4 |
| ran6 | R20:L20 | R20:L5 | -1.059 | 0.291 | 0.291 | ns | 12.58 | 12.9 | -0.31 |
| ran6 | R20:L80 | R20:L5 | 9.976 | 2.9e-19 | 3.48e-19 | **** | 15.98 | 12.9 | 3.09 |
| ran7 | AF | R20:L20 | -8.433 | 6.79e-15 | 8.15e-15 | **** | 13.49 | 16.21 | -2.72 |
| ran7 | AF | R20:L80 | -18.25 | 2.21e-44 | 1.33e-43 | **** | 13.49 | 19.53 | -6.04 |
| ran7 | AF | R20:L5 | -9.393 | 1.4e-17 | 2.8e-17 | **** | 13.49 | 16.6 | -3.105 |
| ran7 | R20:L20 | R20:L80 | -9.775 | 1.12e-18 | 3.36e-18 | **** | 16.21 | 19.53 | -3.32 |
| ran7 | R20:L20 | R20:L5 | -1.108 | 0.269 | 0.269 | ns | 16.21 | 16.6 | -0.385 |
| ran7 | R20:L80 | R20:L5 | 8.541 | 3.45e-15 | 5.18e-15 | **** | 19.53 | 16.6 | 2.935 |
| ran8 | AF | R20:L20 | -8.908 | 3.29e-16 | 4.94e-16 | **** | 12.7 | 15.6 | -2.9 |
| ran8 | AF | R20:L80 | -19.12 | 5.92e-47 | 3.55e-46 | **** | 12.7 | 18.92 | -6.225 |
| ran8 | AF | R20:L5 | -11.71 | 2.03e-24 | 6.09e-24 | **** | 12.7 | 16.44 | -3.745 |
| ran8 | R20:L20 | R20:L80 | -8.922 | 3.01e-16 | 4.94e-16 | **** | 15.6 | 18.92 | -3.325 |
| ran8 | R20:L20 | R20:L5 | -2.502 | 0.013 | 0.013 | * | 15.6 | 16.44 | -0.845 |
| ran8 | R20:L80 | R20:L5 | 7.099 | 2.18e-11 | 2.62e-11 | **** | 18.92 | 16.44 | 2.48 |
| ran9 | AF | R20:L20 | -10.41 | 1.57e-20 | 3.14e-20 | **** | 11.81 | 15.02 | -3.205 |
| ran9 | AF | R20:L80 | -20.62 | 2.78e-51 | 1.67e-50 | **** | 11.81 | 18.24 | -6.435 |
| ran9 | AF | R20:L5 | -14.07 | 1.18e-31 | 3.54e-31 | **** | 11.81 | 16.06 | -4.25 |
| ran9 | R20:L20 | R20:L80 | -9.253 | 3.5e-17 | 5.25e-17 | **** | 15.02 | 18.24 | -3.23 |
| ran9 | R20:L20 | R20:L5 | -3.058 | 0.003 | 0.003 | ** | 15.02 | 16.06 | -1.045 |
| ran9 | R20:L80 | R20:L5 | 5.962 | 1.12e-08 | 1.34e-08 | **** | 18.24 | 16.06 | 2.185 |
| snoRNA | AF | R20:L20 | -13.33 | 2.16e-29 | 4.32e-29 | **** | 23.78 | 29.56 | -5.77 |
| snoRNA | AF | R20:L80 | -22.4 | 2.69e-56 | 1.61e-55 | **** | 23.78 | 34.1 | -10.31 |
| snoRNA | AF | R20:L5 | -16.55 | 2.94e-39 | 8.82e-39 | **** | 23.78 | 31.27 | -7.485 |
| snoRNA | R20:L20 | R20:L80 | -8.639 | 1.85e-15 | 2.78e-15 | **** | 29.56 | 34.1 | -4.54 |
| snoRNA | R20:L20 | R20:L5 | -3.578 | 0.000435 | 0.000435 | *** | 29.56 | 31.27 | -1.715 |
| snoRNA | R20:L80 | R20:L5 | 5.629 | 6.11e-08 | 7.33e-08 | **** | 34.1 | 31.27 | 2.825 |
| tRNAphe | AF | R20:L20 | -10.51 | 7.62e-21 | 1.52e-20 | **** | 16.57 | 20.3 | -3.735 |
| tRNAphe | AF | R20:L80 | -18.01 | 1.21e-43 | 7.26e-43 | **** | 16.57 | 23.28 | -6.71 |
| tRNAphe | AF | R20:L5 | -11.51 | 7.69e-24 | 2.31e-23 | **** | 16.57 | 20.97 | -4.4 |
| tRNAphe | R20:L20 | R20:L80 | -7.336 | 5.45e-12 | 8.18e-12 | **** | 20.3 | 23.28 | -2.975 |
| tRNAphe | R20:L20 | R20:L5 | -1.629 | 0.105 | 0.105 | ns | 20.3 | 20.97 | -0.665 |
| tRNAphe | R20:L80 | R20:L5 | 5.402 | 1.87e-07 | 2.24e-07 | **** | 23.28 | 20.97 | 2.31 |

**Table ST7. Two-way ANOVA for the effect of structure and model on heterozygosity.** Repeated measures under different models.

| Effect | DFn | DFd | $F$ | $p$ | $p < 0.05$ | G. $\eta^2$ |
|---|---|---|---|---|---|---|
| Structure | 14 | 2985 | 205.5 | 0 | * | 0.197 |
| Model | 2.96 | 8848.37 | 145.5 | 5.04e-91 | * | 0.035 |
| Structure:Model | 41.5 | 8848.37 | 1.69 | 0.004 | * | 0.006 |

**Table ST8. One-way ANOVA for the effect of model on heterozygosity.** Repeated measures under different models.

| Structure | DFn | DFd | $F$ | $p$ | $p < 0.05$ | G. $\eta^2$ | Adjusted $p$ |
|---|---|---|---|---|---|---|---|
| CPEB3ribozyme | 2.83 | 562.73 | 17.5 | 2.1e-10 | * | 0.06 | 1.05e-09 |
| DsrA | 2.75 | 546.55 | 11.83 | 4.45e-07 | * | 0.044 | 9.536e-07 |
| HepatitisRibozyme | 2.87 | 571.4 | 5.903 | 7e-04 | * | 0.022 | 0.00075 |
| ran0 | 3 | 597 | 10.24 | 1.39e-06 | * | 0.038 | 2.367e-06 |
| ran1 | 3 | 597 | 13.49 | 1.58e-08 | * | 0.047 | 5.925e-08 |
| ran2 | 2.82 | 562.06 | 13.24 | 5.17e-08 | * | 0.046 | 1.551e-07 |
| ran3 | 3 | 597 | 2.99 | 0.031 | * | 0.011 | 0.031 |
| ran4 | 2.86 | 568.89 | 11.53 | 4.15e-07 | * | 0.039 | 9.536e-07 |
| ran5 | 2.9 | 576.45 | 6.999 | 0.000154 | * | 0.026 | 0.0001777 |
| ran6 | 3 | 597 | 20.34 | 1.46e-12 | * | 0.07 | 2.19e-11 |
| ran7 | 2.87 | 570.78 | 8.965 | 1.2e-05 | * | 0.034 | 1.636e-05 |
| ran8 | 3 | 597 | 8.449 | 1.66e-05 | * | 0.03 | 2.075e-05 |
| ran9 | 3 | 597 | 10.23 | 1.42e-06 | * | 0.038 | 2.367e-06 |
| snoRNA | 3 | 597 | 18.13 | 2.85e-11 | * | 0.064 | 2.138e-10 |
| tRNAphe | 3 | 597 | 9.123 | 6.54e-06 | * | 0.033 | 9.81e-06 |

**Table ST9. Pair-wise post hoc tests for the effect of structure and model on heterozygosity.** Repeated measures under different models.

| Structure | Model M1 | Model M2 | statistic | $p$ | Adjusted $p$ | Signif. | Mean M1 | Mean M2 | Difference |
|---|---|---|---|---|---|---|---|---|---|
| CPEB3 ribozyme | AF | R20:L20 | -1.554 | 0.122 | 0.122 | ns | 0.62 | 0.648 | -0.028 |
| CPEB3 ribozyme | AF | R20:L80 | -6.571 | 4.28e-10 | 2.57e-09 | **** | 0.62 | 0.724 | -0.104 |
| CPEB3 ribozyme | AF | R20:L5 | -4.808 | 3e-06 | 6e-06 | **** | 0.62 | 0.69 | -0.07 |
| CPEB3 ribozyme | R20:L20 | R20:L80 | -5.185 | 5.28e-07 | 1.58e-06 | **** | 0.648 | 0.724 | -0.076 |
| CPEB3 ribozyme | R20:L20 | R20:L5 | -2.682 | 0.008 | 0.012 | * | 0.648 | 0.69 | -0.042 |
| CPEB3 ribozyme | R20:L80 | R20:L5 | 2.4 | 0.017 | 0.021 | * | 0.724 | 0.69 | 0.034 |
| DsrA | AF | R20:L20 | -2.039 | 0.043 | 0.051 | ns | 0.596 | 0.631 | -0.035 |
| DsrA | AF | R20:L80 | -7.215 | 1.11e-11 | 6.66e-11 | **** | 0.596 | 0.691 | -0.095 |
| DsrA | AF | R20:L5 | -2.626 | 0.009 | 0.014 | * | 0.596 | 0.644 | -0.048 |
| DsrA | R20:L20 | R20:L80 | -3.85 | 0.000159 | 0.000477 | *** | 0.631 | 0.691 | -0.06 |
| DsrA | R20:L20 | R20:L5 | -0.7506 | 0.454 | 0.454 | ns | 0.631 | 0.644 | -0.013 |
| DsrA | R20:L80 | R20:L5 | 2.979 | 0.003 | 0.006 | ** | 0.691 | 0.644 | 0.047 |
| Hepatitis ribozyme | AF | R20:L20 | -0.2896 | 0.772 | 0.772 | ns | 0.7 | 0.704 | -0.004 |
| Hepatitis ribozyme | AF | R20:L80 | -4.098 | 6.07e-05 | 0.000364 | *** | 0.7 | 0.752 | -0.052 |
| Hepatitis ribozyme | AF | R20:L5 | -1.2 | 0.231 | 0.347 | ns | 0.7 | 0.716 | -0.016 |
| Hepatitis ribozyme | R20:L20 | R20:L80 | -3.52 | 0.000535 | 0.002 | ** | 0.704 | 0.752 | -0.048 |
| Hepatitis ribozyme | R20:L20 | R20:L5 | -0.7797 | 0.436 | 0.523 | ns | 0.704 | 0.716 | -0.012 |
| Hepatitis ribozyme | R20:L80 | R20:L5 | 2.862 | 0.005 | 0.009 | ** | 0.752 | 0.716 | 0.036 |
| ran0 | AF | R20:L20 | -3.788 | 0.000201 | 0.000402 | *** | 0.561 | 0.627 | -0.066 |
| ran0 | AF | R20:L80 | -4.905 | 1.94e-06 | 1.16e-05 | **** | 0.561 | 0.65 | -0.089 |
| ran0 | AF | R20:L5 | -4.101 | 6e-05 | 0.00018 | *** | 0.561 | 0.632 | -0.071 |
| ran0 | R20:L20 | R20:L80 | -1.35 | 0.178 | 0.267 | ns | 0.627 | 0.65 | -0.023 |
| ran0 | R20:L20 | R20:L5 | -0.2798 | 0.78 | 0.78 | ns | 0.627 | 0.632 | -0.005 |
| ran0 | R20:L80 | R20:L5 | 1.062 | 0.29 | 0.348 | ns | 0.65 | 0.632 | 0.018 |
| ran1 | AF | R20:L20 | -2.756 | 0.006 | 0.01 | ** | 0.472 | 0.528 | -0.056 |
| ran1 | AF | R20:L80 | -6.224 | 2.82e-09 | 1.69e-08 | **** | 0.472 | 0.596 | -0.124 |
| ran1 | AF | R20:L5 | -2.068 | 0.04 | 0.048 | * | 0.472 | 0.516 | -0.044 |
| ran1 | R20:L20 | R20:L80 | -3.59 | 0.000417 | 0.000834 | *** | 0.528 | 0.596 | -0.068 |
| ran1 | R20:L20 | R20:L5 | 0.6386 | 0.524 | 0.524 | ns | 0.528 | 0.516 | 0.012 |
| ran1 | R20:L80 | R20:L5 | 4.205 | 3.95e-05 | 0.000118 | *** | 0.596 | 0.516 | 0.08 |
| ran2 | AF | R20:L20 | -1.304 | 0.194 | 0.194 | ns | 0.606 | 0.63 | -0.024 |
| ran2 | AF | R20:L80 | -6.062 | 6.65e-09 | 3.99e-08 | **** | 0.606 | 0.705 | -0.099 |
| ran2 | AF | R20:L5 | -3.129 | 0.002 | 0.003 | ** | 0.606 | 0.66 | -0.054 |
| ran2 | R20:L20 | R20:L80 | -4.618 | 6.95e-06 | 2.09e-05 | **** | 0.63 | 0.705 | -0.075 |
| ran2 | R20:L20 | R20:L5 | -1.707 | 0.089 | 0.107 | ns | 0.63 | 0.66 | -0.03 |
| ran2 | R20:L80 | R20:L5 | 3.345 | 0.000984 | 0.002 | ** | 0.705 | 0.66 | 0.045 |
| ran3 | AF | R20:L20 | -1.974 | 0.05 | 0.099 | ns | 0.56 | 0.597 | -0.037 |
| ran3 | AF | R20:L80 | -2.796 | 0.006 | 0.034 | * | 0.56 | 0.609 | -0.049 |
| ran3 | AF | R20:L5 | -2.51 | 0.013 | 0.039 | * | 0.56 | 0.605 | -0.045 |
| ran3 | R20:L20 | R20:L80 | -0.62 | 0.536 | 0.791 | ns | 0.597 | 0.609 | -0.012 |
| ran3 | R20:L20 | R20:L5 | -0.4413 | 0.659 | 0.791 | ns | 0.597 | 0.605 | -0.008 |
| ran3 | R20:L80 | R20:L5 | 0.1975 | 0.844 | 0.844 | ns | 0.609 | 0.605 | 0.004 |

| Structure | Model M1 | Model M2 | statistic | $p$ | Adjusted $p$ | Signif. | Mean M1 | Mean M2 | Difference |
|---|---|---|---|---|---|---|---|---|---|
| ran4 | AF | R20:L20 | -2.451 | 0.015 | 0.023 | * | 0.62 | 0.66 | -0.04 |
| ran4 | AF | R20:L80 | -5.383 | 2.05e-07 | 1.23e-06 | **** | 0.62 | 0.705 | -0.085 |
| ran4 | AF | R20:L5 | -3.745 | 0.000236 | 0.000708 | *** | 0.62 | 0.676 | -0.056 |
| ran4 | R20:L20 | R20:L80 | -3.326 | 0.001 | 0.002 | ** | 0.66 | 0.705 | -0.045 |
| ran4 | R20:L20 | R20:L5 | -1.132 | 0.259 | 0.259 | ns | 0.66 | 0.676 | -0.016 |
| ran4 | R20:L80 | R20:L5 | 2.096 | 0.037 | 0.045 | * | 0.705 | 0.676 | 0.029 |
| ran5 | AF | R20:L20 | -2.235 | 0.027 | 0.035 | * | 0.568 | 0.609 | -0.041 |
| ran5 | AF | R20:L80 | -4.465 | 1.34e-05 | 8.04e-05 | **** | 0.568 | 0.649 | -0.081 |
| ran5 | AF | R20:L5 | -2.2 | 0.029 | 0.035 | * | 0.568 | 0.61 | -0.042 |
| ran5 | R20:L20 | R20:L80 | -2.606 | 0.01 | 0.03 | * | 0.609 | 0.649 | -0.04 |
| ran5 | R20:L20 | R20:L5 | -0.03156 | 0.975 | 0.975 | ns | 0.609 | 0.61 | -0.001 |
| ran5 | R20:L80 | R20:L5 | 2.29 | 0.023 | 0.035 | * | 0.649 | 0.61 | 0.039 |
| ran6 | AF | R20:L20 | -3.813 | 0.000183 | 0.000274 | *** | 0.327 | 0.406 | -0.079 |
| ran6 | AF | R20:L80 | -7.616 | 1.03e-12 | 6.18e-12 | **** | 0.327 | 0.487 | -0.16 |
| ran6 | AF | R20:L5 | -2.365 | 0.019 | 0.023 | * | 0.327 | 0.377 | -0.05 |
| ran6 | R20:L20 | R20:L80 | -4.063 | 6.97e-05 | 0.000139 | *** | 0.406 | 0.487 | -0.081 |
| ran6 | R20:L20 | R20:L5 | 1.302 | 0.194 | 0.194 | ns | 0.406 | 0.377 | 0.029 |
| ran6 | R20:L80 | R20:L5 | 5.267 | 3.58e-07 | 1.07e-06 | **** | 0.487 | 0.377 | 0.11 |
| ran7 | AF | R20:L20 | -1.235 | 0.218 | 0.277 | ns | 0.579 | 0.603 | -0.024 |
| ran7 | AF | R20:L80 | -5.11 | 7.53e-07 | 4.52e-06 | **** | 0.579 | 0.664 | -0.085 |
| ran7 | AF | R20:L5 | -1.2 | 0.231 | 0.277 | ns | 0.579 | 0.601 | -0.022 |
| ran7 | R20:L20 | R20:L80 | -3.777 | 0.00021 | 0.00042 | *** | 0.603 | 0.664 | -0.061 |
| ran7 | R20:L20 | R20:L5 | 0.1077 | 0.914 | 0.914 | ns | 0.603 | 0.601 | 0.002 |
| ran7 | R20:L80 | R20:L5 | 4.016 | 8.37e-05 | 0.000251 | *** | 0.664 | 0.601 | 0.063 |
| ran8 | AF | R20:L20 | -1.616 | 0.108 | 0.13 | ns | 0.525 | 0.557 | -0.032 |
| ran8 | AF | R20:L80 | -4.925 | 1.77e-06 | 1.06e-05 | **** | 0.525 | 0.619 | -0.094 |
| ran8 | AF | R20:L5 | -2.538 | 0.012 | 0.024 | * | 0.525 | 0.577 | -0.052 |
| ran8 | R20:L20 | R20:L80 | -3.481 | 0.000614 | 0.002 | ** | 0.557 | 0.619 | -0.062 |
| ran8 | R20:L20 | R20:L5 | -1.038 | 0.301 | 0.301 | ns | 0.557 | 0.577 | -0.02 |
| ran8 | R20:L80 | R20:L5 | 2.285 | 0.023 | 0.035 | * | 0.619 | 0.577 | 0.042 |
| ran9 | AF | R20:L20 | 0.007074 | 0.994 | 0.994 | ns | 0.505 | 0.505 | 0 |
| ran9 | AF | R20:L80 | -4.796 | 3.17e-06 | 1.58e-05 | **** | 0.505 | 0.599 | -0.094 |
| ran9 | AF | R20:L5 | -2.404 | 0.017 | 0.026 | * | 0.505 | 0.555 | -0.05 |
| ran9 | R20:L20 | R20:L80 | -4.681 | 5.27e-06 | 1.58e-05 | **** | 0.505 | 0.599 | -0.094 |
| ran9 | R20:L20 | R20:L5 | -2.557 | 0.011 | 0.023 | * | 0.505 | 0.555 | -0.05 |
| ran9 | R20:L80 | R20:L5 | 2.262 | 0.025 | 0.03 | * | 0.599 | 0.555 | 0.044 |
| snoRNA | AF | R20:L20 | -3.021 | 0.003 | 0.004 | ** | 0.722 | 0.76 | -0.038 |
| snoRNA | AF | R20:L80 | -7.273 | 7.9e-12 | 4.74e-11 | **** | 0.722 | 0.803 | -0.081 |
| snoRNA | AF | R20:L5 | -5.149 | 6.28e-07 | 1.88e-06 | **** | 0.722 | 0.78 | -0.058 |
| snoRNA | R20:L20 | R20:L80 | -3.789 | 2e-04 | 4e-04 | *** | 0.76 | 0.803 | -0.043 |
| snoRNA | R20:L20 | R20:L5 | -1.741 | 0.083 | 0.083 | ns | 0.76 | 0.78 | -0.02 |
| snoRNA | R20:L80 | R20:L5 | 2.209 | 0.028 | 0.034 | * | 0.803 | 0.78 | 0.023 |
| tRNAphe | AF | R20:L20 | -2.813 | 0.005 | 0.014 | * | 0.637 | 0.681 | -0.044 |
| tRNAphe | AF | R20:L80 | -5.282 | 3.33e-07 | 2e-06 | **** | 0.637 | 0.712 | -0.075 |
| tRNAphe | AF | R20:L5 | -2.72 | 0.007 | 0.014 | * | 0.637 | 0.678 | -0.041 |
| tRNAphe | R20:L20 | R20:L80 | -2.221 | 0.028 | 0.033 | * | 0.681 | 0.712 | -0.031 |
| tRNAphe | R20:L20 | R20:L5 | 0.2004 | 0.841 | 0.841 | ns | 0.681 | 0.678 | 0.003 |
| tRNAphe | R20:L80 | R20:L5 | 2.508 | 0.013 | 0.02 | * | 0.712 | 0.678 | 0.034 |

**Table ST10. Two-way ANOVA for the effect of structure and model on maximum sequence divergence.** Repeated measures under different models.

| Effect | DFn | DFd | $F$ | $p$ | $p < 0.05$ | G. $\eta^2$ |
|---|---|---|---|---|---|---|
| Structure | 14 | 2985 | 294 | 0 | * | 0.279 |
| Model | 2.98 | 8895.55 | 189.4 | 2.44e-118 | * | 0.044 |
| Structure:Model | 41.72 | 8895.55 | 1.109 | 0.291 | | 0.004 |

**Table ST11. One-way ANOVA for the effect of model on maximum sequence divergence.** Repeated measures under different models.

| Structure | DFn | DFd | $F$ | $p$ | $p < 0.05$ | G. $\eta^2$ | Adjusted $p$ |
|---|---|---|---|---|---|---|---|
| CPEB3ribozyme | 3 | 597 | 17.46 | 7.03e-11 | * | 0.06 | 2.109e-10 |
| DsrA | 3 | 597 | 16.56 | 2.4e-10 | * | 0.058 | 5.143e-10 |
| HepatitisRibozyme | 3 | 597 | 8.144 | 2.54e-05 | * | 0.027 | 2.721e-05 |
| ran0 | 3 | 597 | 17.14 | 1.09e-10 | * | 0.06 | 2.725e-10 |
| ran1 | 3 | 597 | 29.3 | 1.11e-17 | * | 0.093 | 1.665e-16 |
| ran2 | 3 | 597 | 9.149 | 6.31e-06 | * | 0.031 | 7.888e-06 |
| ran3 | 3 | 597 | 4.009 | 0.008 | * | 0.015 | 0.008 |
| ran4 | 3 | 597 | 8.886 | 9.09e-06 | * | 0.032 | 1.049e-05 |
| ran5 | 3 | 597 | 10 | 1.94e-06 | * | 0.035 | 2.645e-06 |
| ran6 | 2.89 | 575.29 | 29.61 | 2.7e-17 | * | 0.092 | 2.025e-16 |
| ran7 | 3 | 597 | 15.58 | 9.14e-10 | * | 0.056 | 1.714e-09 |
| ran8 | 3 | 597 | 22.11 | 1.38e-13 | * | 0.072 | 6.9e-13 |
| ran9 | 3 | 597 | 21.07 | 5.5e-13 | * | 0.069 | 2.062e-12 |
| snoRNA | 3 | 597 | 10.46 | 1.04e-06 | * | 0.036 | 1.56e-06 |
| tRNAphe | 3 | 597 | 13.63 | 1.3e-08 | * | 0.047 | 2.167e-08 |

**Table ST12. Pair-wise post hoc tests for the effect of structure and model on maximum sequence divergence.** Repeated measures under different models.

| Structure | Model M1 | Model M2 | statistic | $p$ | Adjusted $p$ | Signif. | Mean M1 | Mean M2 | Difference |
|---|---|---|---|---|---|---|---|---|---|
| CPEB3 ribozyme | AF | R20:L20 | -2.806 | 0.006 | 0.007 | ** | 5.8 | 6.45 | -0.65 |
| CPEB3 ribozyme | AF | R20:L80 | -6.923 | 5.94e-11 | 3.56e-10 | **** | 5.8 | 7.42 | -1.62 |
| CPEB3 ribozyme | AF | R20:L5 | -4.096 | 6.11e-05 | 0.000122 | *** | 5.8 | 6.68 | -0.88 |
| CPEB3 ribozyme | R20:L20 | R20:L80 | -4.174 | 4.48e-05 | 0.000122 | *** | 6.45 | 7.42 | -0.97 |
| CPEB3 ribozyme | R20:L20 | R20:L5 | -1.05 | 0.295 | 0.295 | ns | 6.45 | 6.68 | -0.23 |
| CPEB3 ribozyme | R20:L80 | R20:L5 | 3.294 | 0.001 | 0.002 | ** | 7.42 | 6.68 | 0.74 |
| DsrA | AF | R20:L20 | -2.593 | 0.01 | 0.012 | * | 5.48 | 5.96 | -0.48 |
| DsrA | AF | R20:L80 | -7.232 | 1e-11 | 6e-11 | **** | 5.48 | 6.86 | -1.38 |
| DsrA | AF | R20:L5 | -3.881 | 0.000142 | 0.000284 | *** | 5.48 | 6.24 | -0.76 |
| DsrA | R20:L20 | R20:L80 | -4.355 | 2.13e-05 | 6.39e-05 | **** | 5.96 | 6.86 | -0.9 |
| DsrA | R20:L20 | R20:L5 | -1.351 | 0.178 | 0.178 | ns | 5.96 | 6.24 | -0.28 |
| DsrA | R20:L80 | R20:L5 | 2.892 | 0.004 | 0.006 | ** | 6.86 | 6.24 | 0.62 |
| Hepatitis ribozyme | AF | R20:L20 | -0.7441 | 0.458 | 0.458 | ns | 7.25 | 7.445 | -0.195 |
| Hepatitis ribozyme | AF | R20:L80 | -4.88 | 2.17e-06 | 1.3e-05 | **** | 7.25 | 8.44 | -1.19 |
| Hepatitis ribozyme | AF | R20:L5 | -2.002 | 0.047 | 0.07 | ns | 7.25 | 7.755 | -0.505 |
| Hepatitis ribozyme | R20:L20 | R20:L80 | -3.654 | 0.00033 | 0.00099 | *** | 7.445 | 8.44 | -0.995 |
| Hepatitis ribozyme | R20:L20 | R20:L5 | -1.183 | 0.238 | 0.286 | ns | 7.445 | 7.755 | -0.31 |
| Hepatitis ribozyme | R20:L80 | R20:L5 | 2.662 | 0.008 | 0.017 | * | 8.44 | 7.755 | 0.685 |
| ran0 | AF | R20:L20 | -3.079 | 0.002 | 0.004 | ** | 4.995 | 5.575 | -0.58 |
| ran0 | AF | R20:L80 | -7.075 | 2.49e-11 | 1.49e-10 | **** | 4.995 | 6.285 | -1.29 |
| ran0 | AF | R20:L5 | -2.612 | 0.01 | 0.012 | * | 4.995 | 5.435 | -0.44 |
| ran0 | R20:L20 | R20:L80 | -3.7 | 0.000279 | 0.000558 | *** | 5.575 | 6.285 | -0.71 |
| ran0 | R20:L20 | R20:L5 | 0.811 | 0.418 | 0.418 | ns | 5.575 | 5.435 | 0.14 |
| ran0 | R20:L80 | R20:L5 | 4.418 | 1.64e-05 | 4.92e-05 | **** | 6.285 | 5.435 | 0.85 |
| ran1 | AF | R20:L20 | -3.243 | 0.001 | 0.002 | ** | 4.055 | 4.535 | -0.48 |
| ran1 | AF | R20:L80 | -8.472 | 5.33e-15 | 3.2e-14 | **** | 4.055 | 5.4 | -1.345 |
| ran1 | AF | R20:L5 | -2.172 | 0.031 | 0.037 | * | 4.055 | 4.375 | -0.32 |
| ran1 | R20:L20 | R20:L80 | -5.971 | 1.07e-08 | 2.14e-08 | **** | 4.535 | 5.4 | -0.865 |
| ran1 | R20:L20 | R20:L5 | 1.041 | 0.299 | 0.299 | ns | 4.535 | 4.375 | 0.16 |
| ran1 | R20:L80 | R20:L5 | 6.921 | 6.01e-11 | 1.8e-10 | **** | 5.4 | 4.375 | 1.025 |
| ran2 | AF | R20:L20 | -2.925 | 0.004 | 0.008 | ** | 5.52 | 6.185 | -0.665 |
| ran2 | AF | R20:L80 | -5.308 | 2.94e-07 | 1.76e-06 | **** | 5.52 | 6.675 | -1.155 |
| ran2 | AF | R20:L5 | -3.339 | 0.001 | 0.003 | ** | 5.52 | 6.19 | -0.67 |
| ran2 | R20:L20 | R20:L80 | -2.123 | 0.035 | 0.048 | * | 6.185 | 6.675 | -0.49 |
| ran2 | R20:L20 | R20:L5 | -0.02291 | 0.982 | 0.982 | ns | 6.185 | 6.19 | -0.005 |
| ran2 | R20:L80 | R20:L5 | 2.068 | 0.04 | 0.048 | * | 6.675 | 6.19 | 0.485 |
| ran3 | AF | R20:L20 | -0.2278 | 0.82 | 0.82 | ns | 5.11 | 5.15 | -0.04 |
| ran3 | AF | R20:L80 | -2.886 | 0.004 | 0.013 | * | 5.11 | 5.66 | -0.55 |
| ran3 | AF | R20:L5 | -1.472 | 0.143 | 0.214 | ns | 5.11 | 5.38 | -0.27 |
| ran3 | R20:L20 | R20:L80 | -2.977 | 0.003 | 0.013 | * | 5.15 | 5.66 | -0.51 |
| ran3 | R20:L20 | R20:L5 | -1.351 | 0.178 | 0.214 | ns | 5.15 | 5.38 | -0.23 |
| ran3 | R20:L80 | R20:L5 | 1.553 | 0.122 | 0.214 | ns | 5.66 | 5.38 | 0.28 |

| Structure | Model M1 | Model M2 | statistic | $p$ | Adjusted $p$ | Signif. | Mean M1 | Mean M2 | Difference |
|---|---|---|---|---|---|---|---|---|---|
| ran4 | AF | R20:L20 | -2.36 | 0.019 | 0.029 | * | 5.925 | 6.435 | -0.51 |
| ran4 | AF | R20:L80 | -5.264 | 3.63e-07 | 2.18e-06 | **** | 5.925 | 7.035 | -1.11 |
| ran4 | AF | R20:L5 | -3.183 | 0.002 | 0.005 | ** | 5.925 | 6.55 | -0.625 |
| ran4 | R20:L20 | R20:L80 | -2.618 | 0.01 | 0.019 | * | 6.435 | 7.035 | -0.6 |
| ran4 | R20:L20 | R20:L5 | -0.5474 | 0.585 | 0.585 | ns | 6.435 | 6.55 | -0.115 |
| ran4 | R20:L80 | R20:L5 | 2.087 | 0.038 | 0.046 | * | 7.035 | 6.55 | 0.485 |
| ran5 | AF | R20:L20 | -2.67 | 0.008 | 0.012 | * | 5.1 | 5.58 | -0.48 |
| ran5 | AF | R20:L80 | -5.503 | 1.14e-07 | 6.84e-07 | **** | 5.1 | 6.085 | -0.985 |
| ran5 | AF | R20:L5 | -1.819 | 0.07 | 0.084 | ns | 5.1 | 5.455 | -0.355 |
| ran5 | R20:L20 | R20:L80 | -2.847 | 0.005 | 0.01 | ** | 5.58 | 6.085 | -0.505 |
| ran5 | R20:L20 | R20:L5 | 0.7117 | 0.478 | 0.478 | ns | 5.58 | 5.455 | 0.125 |
| ran5 | R20:L80 | R20:L5 | 3.385 | 0.000857 | 0.003 | ** | 6.085 | 5.455 | 0.63 |
| ran6 | AF | R20:L20 | -5.24 | 4.07e-07 | 1.22e-06 | **** | 3.07 | 3.69 | -0.62 |
| ran6 | AF | R20:L80 | -9.293 | 2.71e-17 | 1.63e-16 | **** | 3.07 | 4.235 | -1.165 |
| ran6 | AF | R20:L5 | -4.99 | 1.31e-06 | 2.62e-06 | **** | 3.07 | 3.62 | -0.55 |
| ran6 | R20:L20 | R20:L80 | -4.253 | 3.25e-05 | 3.9e-05 | **** | 3.69 | 4.235 | -0.545 |
| ran6 | R20:L20 | R20:L5 | 0.5659 | 0.572 | 0.572 | ns | 3.69 | 3.62 | 0.07 |
| ran6 | R20:L80 | R20:L5 | 4.531 | 1.01e-05 | 1.52e-05 | **** | 4.235 | 3.62 | 0.615 |
| ran7 | AF | R20:L20 | -2.67 | 0.008 | 0.01 | * | 4.92 | 5.415 | -0.495 |
| ran7 | AF | R20:L80 | -7.503 | 2.04e-12 | 1.22e-11 | **** | 4.92 | 6.18 | -1.26 |
| ran7 | AF | R20:L5 | -2.663 | 0.008 | 0.01 | * | 4.92 | 5.41 | -0.49 |
| ran7 | R20:L20 | R20:L80 | -3.808 | 0.000186 | 0.000372 | *** | 5.415 | 6.18 | -0.765 |
| ran7 | R20:L20 | R20:L5 | 0.0272 | 0.978 | 0.978 | ns | 5.415 | 5.41 | 0.005 |
| ran7 | R20:L80 | R20:L5 | 3.943 | 0.000111 | 0.000333 | *** | 6.18 | 5.41 | 0.77 |
| ran8 | AF | R20:L20 | -3.06 | 0.003 | 0.003 | ** | 4.44 | 4.925 | -0.485 |
| ran8 | AF | R20:L80 | -7.923 | 1.61e-13 | 9.66e-13 | **** | 4.44 | 5.79 | -1.35 |
| ran8 | AF | R20:L5 | -3.986 | 9.43e-05 | 0.000141 | *** | 4.44 | 5.08 | -0.64 |
| ran8 | R20:L20 | R20:L80 | -4.966 | 1.46e-06 | 4.38e-06 | **** | 4.925 | 5.79 | -0.865 |
| ran8 | R20:L20 | R20:L5 | -0.9306 | 0.353 | 0.353 | ns | 4.925 | 5.08 | -0.155 |
| ran8 | R20:L80 | R20:L5 | 4.011 | 8.56e-05 | 0.000141 | *** | 5.79 | 5.08 | 0.71 |
| ran9 | AF | R20:L20 | -1.511 | 0.132 | 0.132 | ns | 4.325 | 4.57 | -0.245 |
| ran9 | AF | R20:L80 | -7.695 | 6.46e-13 | 3.88e-12 | **** | 4.325 | 5.525 | -1.2 |
| ran9 | AF | R20:L5 | -4.085 | 6.38e-05 | 0.000128 | *** | 4.325 | 4.985 | -0.66 |
| ran9 | R20:L20 | R20:L80 | -5.691 | 4.46e-08 | 1.34e-07 | **** | 4.57 | 5.525 | -0.955 |
| ran9 | R20:L20 | R20:L5 | -2.66 | 0.008 | 0.01 | * | 4.57 | 4.985 | -0.415 |
| ran9 | R20:L80 | R20:L5 | 3.231 | 0.001 | 0.002 | ** | 5.525 | 4.985 | 0.54 |
| snoRNA | AF | R20:L20 | -2.547 | 0.012 | 0.014 | * | 8.325 | 9.04 | -0.715 |
| snoRNA | AF | R20:L80 | -5.714 | 3.97e-08 | 2.38e-07 | **** | 8.325 | 9.965 | -1.64 |
| snoRNA | AF | R20:L5 | -2.966 | 0.003 | 0.007 | ** | 8.325 | 9.16 | -0.835 |
| snoRNA | R20:L20 | R20:L80 | -2.895 | 0.004 | 0.007 | ** | 9.04 | 9.965 | -0.925 |
| snoRNA | R20:L20 | R20:L5 | -0.3845 | 0.701 | 0.701 | ns | 9.04 | 9.16 | -0.12 |
| snoRNA | R20:L80 | R20:L5 | 2.879 | 0.004 | 0.007 | ** | 9.965 | 9.16 | 0.805 |
| tRNAphe | AF | R20:L20 | -4.692 | 5.03e-06 | 1.51e-05 | **** | 6 | 7 | -1 |
| tRNAphe | AF | R20:L80 | -6.43 | 9.26e-10 | 5.56e-09 | **** | 6 | 7.37 | -1.37 |
| tRNAphe | AF | R20:L5 | -3.109 | 0.002 | 0.004 | ** | 6 | 6.65 | -0.65 |
| tRNAphe | R20:L20 | R20:L80 | -1.517 | 0.131 | 0.131 | ns | 7 | 7.37 | -0.37 |
| tRNAphe | R20:L20 | R20:L5 | 1.589 | 0.114 | 0.131 | ns | 7 | 6.65 | 0.35 |
| tRNAphe | R20:L80 | R20:L5 | 3.028 | 0.003 | 0.004 | ** | 7.37 | 6.65 | 0.72 |

**Table ST13. Two-way ANOVA for the effect of structure and model on nucleotide diversity.** Repeated measures under different models.

| Effect | DFn | DFd | $F$ | $p$ | $p < 0.05$ | G. $\eta^2$ |
|---|---|---|---|---|---|---|
| Structure | 14 | 2985 | 241.4 | 0 | * | 0.235 |
| Model | 2.99 | 8914.17 | 102.1 | 1.08e-64 | * | 0.024 |
| Structure:Model | 41.81 | 8914.17 | 1.134 | 0.255 | | 0.004 |

**Table ST14. One-way ANOVA for the effect of model on nucleotide diversity.** Repeated measures under different models.

| Structure | DFn | DFd | $F$ | $p$ | $p < 0.05$ | G. $\eta^2$ | Adjusted $p$ |
|---|---|---|---|---|---|---|---|
| CPEB3ribozyme | 3 | 597 | 10.1 | 1.69e-06 | * | 0.036 | 5.07e-06 |
| DsrA | 3 | 597 | 7.522 | 6.01e-05 | * | 0.028 | 9.9e-05 |
| HepatitisRibozyme | 3 | 597 | 5.839 | 0.00062 | * | 0.02 | 0.0007154 |
| ran0 | 3 | 597 | 9.02 | 7.55e-06 | * | 0.032 | 1.871e-05 |
| ran1 | 3 | 597 | 16.28 | 3.52e-10 | * | 0.053 | 2.64e-09 |
| ran2 | 3 | 597 | 5.901 | 0.000568 | * | 0.021 | 0.00071 |
| ran3 | 3 | 597 | 1.174 | 0.319 |  | 0.004 | 0.319 |
| ran4 | 3 | 597 | 7.456 | 6.6e-05 | * | 0.028 | 9.9e-05 |
| ran5 | 3 | 597 | 4.5 | 0.004 | * | 0.016 | 0.004286 |
| ran6 | 3 | 597 | 16.3 | 3.39e-10 | * | 0.056 | 2.64e-09 |
| ran7 | 3 | 597 | 8.914 | 8.73e-06 | * | 0.033 | 1.871e-05 |
| ran8 | 2.87 | 572.07 | 10.94 | 8.45e-07 | * | 0.038 | 3.169e-06 |
| ran9 | 3 | 597 | 13.3 | 2.06e-08 | * | 0.045 | 1.03e-07 |
| snoRNA | 3 | 597 | 7.547 | 5.81e-05 | * | 0.027 | 9.9e-05 |
| tRNAphe | 3 | 597 | 7.041 | 0.000117 | * | 0.025 | 0.0001595 |

**Table ST15. Pair-wise post hoc tests for the effect of structure and model on nucleotide diversity.** Repeated measures under different models.

| Structure | Model M1 | Model M2 | statistic | $p$ | Adjusted $p$ | Signif. | Mean M1 | Mean M2 | Difference |
|---|---|---|---|---|---|---|---|---|---|
| CPEB3 ribozyme | AF | R20:L20 | -0.7441 | 0.458 | 0.458 | ns | 1.571 | 1.649 | -0.078 |
| CPEB3 ribozyme | AF | R20:L80 | -4.909 | 1.9e-06 | 1.14e-05 | **** | 1.571 | 2.083 | -0.512 |
| CPEB3 ribozyme | AF | R20:L5 | -2.508 | 0.013 | 0.019 | * | 1.571 | 1.81 | -0.239 |
| CPEB3 ribozyme | R20:L20 | R20:L80 | -4.137 | 5.18e-05 | 0.000155 | *** | 1.649 | 2.083 | -0.434 |
| CPEB3 ribozyme | R20:L20 | R20:L5 | -1.708 | 0.089 | 0.107 | ns | 1.649 | 1.81 | -0.161 |
| CPEB3 ribozyme | R20:L80 | R20:L5 | 2.704 | 0.007 | 0.015 | * | 2.083 | 1.81 | 0.273 |
| DsrA | AF | R20:L20 | -1.137 | 0.257 | 0.257 | ns | 1.362 | 1.458 | -0.096 |
| DsrA | AF | R20:L80 | -4.853 | 2.45e-06 | 1.47e-05 | **** | 1.362 | 1.755 | -0.393 |
| DsrA | AF | R20:L5 | -2.244 | 0.026 | 0.052 | ns | 1.362 | 1.563 | -0.201 |
| DsrA | R20:L20 | R20:L80 | -3.357 | 0.000943 | 0.003 | ** | 1.458 | 1.755 | -0.297 |
| DsrA | R20:L20 | R20:L5 | -1.269 | 0.206 | 0.247 | ns | 1.458 | 1.563 | -0.105 |
| DsrA | R20:L80 | R20:L5 | 2.04 | 0.043 | 0.064 | ns | 1.755 | 1.563 | 0.192 |
| Hepatitis ribozyme | AF | R20:L20 | 0.548 | 0.584 | 0.584 | ns | 2.122 | 2.056 | 0.066 |
| Hepatitis ribozyme | AF | R20:L80 | -3.297 | 0.001 | 0.003 | ** | 2.122 | 2.514 | -0.392 |
| Hepatitis ribozyme | AF | R20:L5 | -0.8144 | 0.416 | 0.499 | ns | 2.122 | 2.214 | -0.092 |
| Hepatitis ribozyme | R20:L20 | R20:L80 | -3.703 | 0.000276 | 0.002 | ** | 2.056 | 2.514 | -0.458 |
| Hepatitis ribozyme | R20:L20 | R20:L5 | -1.323 | 0.187 | 0.28 | ns | 2.056 | 2.214 | -0.158 |
| Hepatitis ribozyme | R20:L80 | R20:L5 | 2.637 | 0.009 | 0.018 | * | 2.514 | 2.214 | 0.3 |
| ran0 | AF | R20:L20 | -2.362 | 0.019 | 0.029 | * | 1.184 | 1.368 | -0.184 |
| ran0 | AF | R20:L80 | -4.773 | 3.51e-06 | 2.11e-05 | **** | 1.184 | 1.587 | -0.403 |
| ran0 | AF | R20:L5 | -1.959 | 0.052 | 0.062 | ns | 1.184 | 1.324 | -0.14 |
| ran0 | R20:L20 | R20:L80 | -2.759 | 0.006 | 0.013 | * | 1.368 | 1.587 | -0.219 |
| ran0 | R20:L20 | R20:L5 | 0.5922 | 0.554 | 0.554 | ns | 1.368 | 1.324 | 0.044 |
| ran0 | R20:L80 | R20:L5 | 3.132 | 0.002 | 0.006 | ** | 1.587 | 1.324 | 0.263 |
| ran1 | AF | R20:L20 | -1.965 | 0.051 | 0.076 | ns | 0.853 | 0.976 | -0.123 |
| ran1 | AF | R20:L80 | -6.195 | 3.29e-09 | 1.97e-08 | **** | 0.853 | 1.253 | -0.4 |
| ran1 | AF | R20:L5 | -0.8637 | 0.389 | 0.389 | ns | 0.853 | 0.905 | -0.052 |
| ran1 | R20:L20 | R20:L80 | -4.241 | 3.4e-05 | 6.8e-05 | **** | 0.976 | 1.253 | -0.277 |
| ran1 | R20:L20 | R20:L5 | 1.216 | 0.226 | 0.271 | ns | 0.976 | 0.905 | 0.071 |
| ran1 | R20:L80 | R20:L5 | 5.563 | 8.47e-08 | 2.54e-07 | **** | 1.253 | 0.905 | 0.348 |
| ran2 | AF | R20:L20 | -1.537 | 0.126 | 0.151 | ns | 1.42 | 1.568 | -0.148 |
| ran2 | AF | R20:L80 | -4.151 | 4.91e-05 | 0.000295 | *** | 1.42 | 1.823 | -0.403 |
| ran2 | AF | R20:L5 | -2.257 | 0.025 | 0.05 | ns | 1.42 | 1.62 | -0.2 |
| ran2 | R20:L20 | R20:L80 | -2.49 | 0.014 | 0.041 | * | 1.568 | 1.823 | -0.255 |
| ran2 | R20:L20 | R20:L5 | -0.5488 | 0.584 | 0.584 | ns | 1.568 | 1.62 | -0.052 |
| ran2 | R20:L80 | R20:L5 | 1.975 | 0.05 | 0.074 | ns | 1.823 | 1.62 | 0.203 |
| ran3 | AF | R20:L20 | 0.4808 | 0.631 | 0.66 | ns | 1.258 | 1.222 | 0.036 |
| ran3 | AF | R20:L80 | -1.249 | 0.213 | 0.582 | ns | 1.258 | 1.361 | -0.103 |
| ran3 | AF | R20:L5 | -0.4407 | 0.66 | 0.66 | ns | 1.258 | 1.294 | -0.036 |
| ran3 | R20:L20 | R20:L80 | -1.85 | 0.066 | 0.395 | ns | 1.222 | 1.361 | -0.139 |
| ran3 | R20:L20 | R20:L5 | -1.008 | 0.315 | 0.582 | ns | 1.222 | 1.294 | -0.072 |
| ran3 | R20:L80 | R20:L5 | 0.8656 | 0.388 | 0.582 | ns | 1.361 | 1.294 | 0.067 |

| Structure | Model M1 | Model M2 | statistic | $p$ | Adjusted $p$ | Signif. | Mean M1 | Mean M2 | Difference |
|---|---|---|---|---|---|---|---|---|---|
| ran4 | AF | R20:L20 | -1.562 | 0.12 | 0.144 | ns | 1.509 | 1.662 | -0.153 |
| ran4 | AF | R20:L80 | -4.82 | 2.84e-06 | 1.7e-05 | **** | 1.509 | 1.952 | -0.443 |
| ran4 | AF | R20:L5 | -2.65 | 0.009 | 0.017 | * | 1.509 | 1.743 | -0.234 |
| ran4 | R20:L20 | R20:L80 | -2.893 | 0.004 | 0.013 | * | 1.662 | 1.952 | -0.29 |
| ran4 | R20:L20 | R20:L5 | -0.8626 | 0.389 | 0.389 | ns | 1.662 | 1.743 | -0.081 |
| ran4 | R20:L80 | R20:L5 | 2.079 | 0.039 | 0.058 | ns | 1.952 | 1.743 | 0.209 |
| ran5 | AF | R20:L20 | -1.399 | 0.163 | 0.244 | ns | 1.241 | 1.354 | -0.113 |
| ran5 | AF | R20:L80 | -3.452 | 0.00068 | 0.004 | ** | 1.241 | 1.522 | -0.281 |
| ran5 | AF | R20:L5 | -0.5638 | 0.574 | 0.574 | ns | 1.241 | 1.291 | -0.05 |
| ran5 | R20:L20 | R20:L80 | -2.171 | 0.031 | 0.062 | ns | 1.354 | 1.522 | -0.168 |
| ran5 | R20:L20 | R20:L5 | 0.7855 | 0.433 | 0.52 | ns | 1.354 | 1.291 | 0.063 |
| ran5 | R20:L80 | R20:L5 | 2.846 | 0.005 | 0.015 | * | 1.522 | 1.291 | 0.231 |
| ran6 | AF | R20:L20 | -3.597 | 0.000406 | 0.000812 | *** | 0.47 | 0.622 | -0.152 |
| ran6 | AF | R20:L80 | -6.656 | 2.67e-10 | 1.6e-09 | **** | 0.47 | 0.778 | -0.308 |
| ran6 | AF | R20:L5 | -2.033 | 0.043 | 0.052 | ns | 0.47 | 0.554 | -0.084 |
| ran6 | R20:L20 | R20:L80 | -3.25 | 0.001 | 0.002 | ** | 0.622 | 0.778 | -0.156 |
| ran6 | R20:L20 | R20:L5 | 1.412 | 0.159 | 0.159 | ns | 0.622 | 0.554 | 0.068 |
| ran6 | R20:L80 | R20:L5 | 4.68 | 5.31e-06 | 1.59e-05 | **** | 0.778 | 0.554 | 0.224 |
| ran7 | AF | R20:L20 | -0.6897 | 0.491 | 0.589 | ns | 1.231 | 1.285 | -0.054 |
| ran7 | AF | R20:L80 | -4.875 | 2.22e-06 | 1.33e-05 | **** | 1.231 | 1.615 | -0.384 |
| ran7 | AF | R20:L5 | -0.8601 | 0.391 | 0.587 | ns | 1.231 | 1.306 | -0.075 |
| ran7 | R20:L20 | R20:L80 | -4.076 | 6.63e-05 | 0.000199 | *** | 1.285 | 1.615 | -0.33 |
| ran7 | R20:L20 | R20:L5 | -0.2599 | 0.795 | 0.795 | ns | 1.285 | 1.306 | -0.021 |
| ran7 | R20:L80 | R20:L5 | 3.537 | 0.000504 | 0.001 | ** | 1.615 | 1.306 | 0.309 |
| ran8 | AF | R20:L20 | -1.974 | 0.05 | 0.06 | ns | 0.991 | 1.123 | -0.132 |
| ran8 | AF | R20:L80 | -5.556 | 8.77e-08 | 5.26e-07 | **** | 0.991 | 1.405 | -0.414 |
| ran8 | AF | R20:L5 | -2.536 | 0.012 | 0.018 | * | 0.991 | 1.162 | -0.171 |
| ran8 | R20:L20 | R20:L80 | -3.543 | 0.000492 | 0.001 | ** | 1.123 | 1.405 | -0.282 |
| ran8 | R20:L20 | R20:L5 | -0.533 | 0.595 | 0.595 | ns | 1.123 | 1.162 | -0.039 |
| ran8 | R20:L80 | R20:L5 | 2.988 | 0.003 | 0.006 | ** | 1.405 | 1.162 | 0.243 |
| ran9 | AF | R20:L20 | -0.03265 | 0.974 | 0.974 | ns | 0.926 | 0.928 | -0.002 |
| ran9 | AF | R20:L80 | -5.307 | 2.96e-07 | 1.12e-06 | **** | 0.926 | 1.291 | -0.365 |
| ran9 | AF | R20:L5 | -2.756 | 0.006 | 0.01 | ** | 0.926 | 1.124 | -0.198 |
| ran9 | R20:L20 | R20:L80 | -5.259 | 3.72e-07 | 1.12e-06 | **** | 0.928 | 1.291 | -0.363 |
| ran9 | R20:L20 | R20:L5 | -3.092 | 0.002 | 0.005 | ** | 0.928 | 1.124 | -0.196 |
| ran9 | R20:L80 | R20:L5 | 2.322 | 0.021 | 0.026 | * | 1.291 | 1.124 | 0.167 |
| snoRNA | AF | R20:L20 | -2.226 | 0.027 | 0.041 | * | 2.448 | 2.739 | -0.291 |
| snoRNA | AF | R20:L80 | -4.738 | 4.1e-06 | 2.46e-05 | **** | 2.448 | 3.096 | -0.648 |
| snoRNA | AF | R20:L5 | -2.082 | 0.039 | 0.046 | * | 2.448 | 2.721 | -0.273 |
| snoRNA | R20:L20 | R20:L80 | -2.433 | 0.016 | 0.032 | * | 2.739 | 3.096 | -0.357 |
| snoRNA | R20:L20 | R20:L5 | 0.1267 | 0.899 | 0.899 | ns | 2.739 | 2.721 | 0.018 |
| snoRNA | R20:L80 | R20:L5 | 2.84 | 0.005 | 0.015 | * | 3.096 | 2.721 | 0.375 |
| tRNAphe | AF | R20:L20 | -3.011 | 0.003 | 0.009 | ** | 1.591 | 1.868 | -0.277 |
| tRNAphe | AF | R20:L80 | -4.442 | 1.48e-05 | 8.88e-05 | **** | 1.591 | 2.019 | -0.428 |
| tRNAphe | AF | R20:L5 | -1.72 | 0.087 | 0.131 | ns | 1.591 | 1.752 | -0.161 |
| tRNAphe | R20:L20 | R20:L80 | -1.508 | 0.133 | 0.16 | ns | 1.868 | 2.019 | -0.151 |
| tRNAphe | R20:L20 | R20:L5 | 1.239 | 0.217 | 0.217 | ns | 1.868 | 1.752 | 0.116 |
| tRNAphe | R20:L80 | R20:L5 | 2.599 | 0.01 | 0.02 | * | 2.019 | 1.752 | 0.267 |

**Table ST16. Two-way ANOVA for the effect of structure and model on Boltzmann probability of optimal structure.** Repeated measures under different models.

| Effect | DFn | DFd | $F$ | $p$ | $p < 0.05$ | G. $\eta^2$ |
|---|---|---|---|---|---|---|
| Structure | 14 | 2985 | 111.4 | 7.33e-260 | * | 0.29 |
| Model | 2.83 | 8437.57 | 2130 | 0 | * | 0.134 |
| Structure:Model | 39.57 | 8437.57 | 5.031 | 8.17e-23 | * | 0.005 |

**Table ST17. One-way ANOVA for the effect of model on Boltzmann probability of optimal structure.** Repeated measures under different models.

| Structure | DFn | DFd | $F$ | $p$ | $p < 0.05$ | G. $\eta^2$ | Adjusted $p$ |
|---|---|---|---|---|---|---|---|
| CPEB3ribozyme | 2.31 | 460.46 | 150.5 | 1.1e-56 | * | 0.149 | 1.5e-56 |
| DsrA | 2.37 | 472.19 | 160.7 | 5.04e-61 | * | 0.172 | 9.45e-61 |
| HepatitisRibozyme | 2.51 | 499.81 | 341 | 1.8e-108 | * | 0.212 | 9e-108 |
| ran0 | 2.17 | 432.33 | 175.4 | 7.18e-60 | * | 0.244 | 1.077e-59 |
| ran1 | 3 | 597 | 43.24 | 2.69e-25 | * | 0.045 | 2.69e-25 |
| ran2 | 2.45 | 486.56 | 153.1 | 1.55e-60 | * | 0.17 | 2.583e-60 |
| ran3 | 2.55 | 507.76 | 180.3 | 3.16e-71 | * | 0.12 | 9.48e-71 |
| ran4 | 2.65 | 526.56 | 157.1 | 1.55e-66 | * | 0.157 | 3.321e-66 |
| ran5 | 2.77 | 552.12 | 114.8 | 1.71e-54 | * | 0.117 | 2.138e-54 |
| ran6 | 2.83 | 563.55 | 101 | 4.54e-50 | * | 0.102 | 5.238e-50 |
| ran7 | 2.57 | 511.99 | 166.6 | 1.03e-67 | * | 0.13 | 2.575e-67 |
| ran8 | 2.85 | 567.94 | 64.85 | 1.14e-34 | * | 0.053 | 1.221e-34 |
| ran9 | 2.48 | 493.18 | 212.8 | 4.59e-78 | * | 0.192 | 1.721e-77 |
| snoRNA | 2.48 | 492.61 | 511.7 | 2.57e-136 | * | 0.453 | 3.855e-135 |
| tRNAphe | 2.69 | 534.61 | 320.4 | 2.74e-111 | * | 0.245 | 2.055e-110 |

**Table ST18. Pair-wise post hoc tests for the effect of structure and model on Boltzmann probability of optimal structure.** Repeated measures under different models.

| Structure | Model M1 | Model M2 | statistic | $p$ | Adjusted $p$ | Signif. | Mean M1 | Mean M2 | Difference |
|---|---|---|---|---|---|---|---|---|---|
| CPEB3 ribozyme | AF | R20:L20 | 4.895 | 2.03e-06 | 2.44e-06 | **** | 0.837 | 0.804 | 0.033 |
| CPEB3 ribozyme | AF | R20:L80 | 19.37 | 1.14e-47 | 6.84e-47 | **** | 0.837 | 0.691 | 0.146 |
| CPEB3 ribozyme | AF | R20:L5 | 9.613 | 3.26e-18 | 4.89e-18 | **** | 0.837 | 0.792 | 0.045 |
| CPEB3 ribozyme | R20:L20 | R20:L80 | 12.03 | 2.03e-25 | 4.06e-25 | **** | 0.804 | 0.691 | 0.113 |
| CPEB3 ribozyme | R20:L20 | R20:L5 | 1.796 | 0.074 | 0.074 | ns | 0.804 | 0.792 | 0.012 |
| CPEB3 ribozyme | R20:L80 | R20:L5 | -12.93 | 3.68e-28 | 1.1e-27 | **** | 0.691 | 0.792 | -0.101 |
| DsrA | AF | R20:L20 | 7.399 | 3.78e-12 | 4.54e-12 | **** | 0.82 | 0.772 | 0.048 |
| DsrA | AF | R20:L80 | 18.26 | 2.13e-44 | 1.28e-43 | **** | 0.82 | 0.654 | 0.166 |
| DsrA | AF | R20:L5 | 10.56 | 5.49e-21 | 8.24e-21 | **** | 0.82 | 0.757 | 0.063 |
| DsrA | R20:L20 | R20:L80 | 12.63 | 3.04e-27 | 9.12e-27 | **** | 0.772 | 0.654 | 0.118 |
| DsrA | R20:L20 | R20:L5 | 2.409 | 0.017 | 0.017 | * | 0.772 | 0.757 | 0.015 |
| DsrA | R20:L80 | R20:L5 | -11.67 | 2.55e-24 | 5.1e-24 | **** | 0.654 | 0.757 | -0.103 |
| Hepatitis ribozyme | AF | R20:L20 | 8.652 | 1.7e-15 | 2.04e-15 | **** | 0.711 | 0.681 | 0.03 |
| Hepatitis ribozyme | AF | R20:L80 | 26.95 | 2.54e-68 | 1.52e-67 | **** | 0.711 | 0.574 | 0.137 |
| Hepatitis ribozyme | AF | R20:L5 | 10.41 | 1.59e-20 | 2.39e-20 | **** | 0.711 | 0.67 | 0.041 |
| Hepatitis ribozyme | R20:L20 | R20:L80 | 20.34 | 1.71e-50 | 5.13e-50 | **** | 0.681 | 0.574 | 0.107 |
| Hepatitis ribozyme | R20:L20 | R20:L5 | 2.855 | 0.005 | 0.005 | ** | 0.681 | 0.67 | 0.011 |
| Hepatitis ribozyme | R20:L80 | R20:L5 | -18.44 | 6.23e-45 | 1.25e-44 | **** | 0.574 | 0.67 | -0.096 |
| ran0 | AF | R20:L20 | 6.594 | 3.78e-10 | 4.54e-10 | **** | 0.92 | 0.887 | 0.033 |
| ran0 | AF | R20:L80 | 18.3 | 1.59e-44 | 9.54e-44 | **** | 0.92 | 0.766 | 0.154 |
| ran0 | AF | R20:L5 | 10.79 | 1.18e-21 | 1.77e-21 | **** | 0.92 | 0.861 | 0.059 |
| ran0 | R20:L20 | R20:L80 | 14.36 | 1.52e-32 | 4.56e-32 | **** | 0.887 | 0.766 | 0.121 |
| ran0 | R20:L20 | R20:L5 | 4.843 | 2.57e-06 | 2.57e-06 | **** | 0.887 | 0.861 | 0.026 |
| ran0 | R20:L80 | R20:L5 | -11.06 | 1.76e-22 | 3.52e-22 | **** | 0.766 | 0.861 | -0.095 |
| ran1 | AF | R20:L20 | 5.204 | 4.83e-07 | 7.24e-07 | **** | 0.643 | 0.572 | 0.071 |
| ran1 | AF | R20:L80 | 12.24 | 4.83e-26 | 2.9e-25 | **** | 0.643 | 0.498 | 0.145 |
| ran1 | AF | R20:L5 | 5.975 | 1.04e-08 | 2.08e-08 | **** | 0.643 | 0.566 | 0.077 |
| ran1 | R20:L20 | R20:L80 | 6.111 | 5.12e-09 | 1.54e-08 | **** | 0.572 | 0.498 | 0.074 |
| ran1 | R20:L20 | R20:L5 | 0.5007 | 0.617 | 0.617 | ns | 0.572 | 0.566 | 0.006 |
| ran1 | R20:L80 | R20:L5 | -5.147 | 6.34e-07 | 7.61e-07 | **** | 0.498 | 0.566 | -0.068 |
| ran2 | AF | R20:L20 | 4.477 | 1.27e-05 | 1.52e-05 | **** | 0.878 | 0.85 | 0.028 |
| ran2 | AF | R20:L80 | 16.54 | 3.1e-39 | 1.86e-38 | **** | 0.878 | 0.734 | 0.144 |
| ran2 | AF | R20:L5 | 8.306 | 1.51e-14 | 2.26e-14 | **** | 0.878 | 0.83 | 0.048 |
| ran2 | R20:L20 | R20:L80 | 14.94 | 2.37e-34 | 7.11e-34 | **** | 0.85 | 0.734 | 0.116 |
| ran2 | R20:L20 | R20:L5 | 3.496 | 0.000582 | 0.000582 | *** | 0.85 | 0.83 | 0.02 |
| ran2 | R20:L80 | R20:L5 | -11.86 | 6.89e-25 | 1.38e-24 | **** | 0.734 | 0.83 | -0.096 |
| ran3 | AF | R20:L20 | 4.157 | 4.79e-05 | 5.75e-05 | **** | 0.829 | 0.805 | 0.024 |
| ran3 | AF | R20:L80 | 19.85 | 4.53e-49 | 2.72e-48 | **** | 0.829 | 0.681 | 0.148 |
| ran3 | AF | R20:L5 | 7.237 | 9.71e-12 | 1.46e-11 | **** | 0.829 | 0.785 | 0.044 |
| ran3 | R20:L20 | R20:L80 | 17.14 | 4.7e-41 | 1.41e-40 | **** | 0.805 | 0.681 | 0.124 |
| ran3 | R20:L20 | R20:L5 | 3.388 | 0.000849 | 0.000849 | *** | 0.805 | 0.785 | 0.02 |
| ran3 | R20:L80 | R20:L5 | -12.4 | 1.51e-26 | 3.02e-26 | **** | 0.681 | 0.785 | -0.104 |

| Structure | Model M1 | Model M2 | statistic | $p$ | Adjusted $p$ | Signif. | Mean M1 | Mean M2 | Difference |
|---|---|---|---|---|---|---|---|---|---|
| ran4 | AF | R20:L20 | 5.61 | 6.72e-08 | 8.06e-08 | **** | 0.892 | 0.864 | 0.028 |
| ran4 | AF | R20:L80 | 20.59 | 3.23e-51 | 1.94e-50 | **** | 0.892 | 0.764 | 0.128 |
| ran4 | AF | R20:L5 | 8.37 | 1.01e-14 | 1.51e-14 | **** | 0.892 | 0.844 | 0.048 |
| ran4 | R20:L20 | R20:L80 | 14.68 | 1.5e-33 | 4.5e-33 | **** | 0.864 | 0.764 | 0.1 |
| ran4 | R20:L20 | R20:L5 | 3.378 | 0.000877 | 0.000877 | *** | 0.864 | 0.844 | 0.02 |
| ran4 | R20:L80 | R20:L5 | -10.89 | 5.98e-22 | 1.2e-21 | **** | 0.764 | 0.844 | -0.08 |
| ran5 | AF | R20:L20 | 4.443 | 1.47e-05 | 1.76e-05 | **** | 0.777 | 0.74 | 0.037 |
| ran5 | AF | R20:L80 | 16.9 | 2.55e-40 | 1.53e-39 | **** | 0.777 | 0.627 | 0.15 |
| ran5 | AF | R20:L5 | 6.656 | 2.68e-10 | 4.02e-10 | **** | 0.777 | 0.717 | 0.06 |
| ran5 | R20:L20 | R20:L80 | 12.68 | 2.15e-27 | 4.3e-27 | **** | 0.74 | 0.627 | 0.113 |
| ran5 | R20:L20 | R20:L5 | 2.815 | 0.005 | 0.005 | ** | 0.74 | 0.717 | 0.023 |
| ran5 | R20:L80 | R20:L5 | -12.91 | 4.4e-28 | 1.32e-27 | **** | 0.627 | 0.717 | -0.09 |
| ran6 | AF | R20:L20 | 4.611 | 7.17e-06 | 8.6e-06 | **** | 0.684 | 0.652 | 0.032 |
| ran6 | AF | R20:L80 | 15.45 | 6.39e-36 | 3.83e-35 | **** | 0.684 | 0.544 | 0.14 |
| ran6 | AF | R20:L5 | 7.177 | 1.38e-11 | 2.07e-11 | **** | 0.684 | 0.624 | 0.06 |
| ran6 | R20:L20 | R20:L80 | 12.65 | 2.75e-27 | 8.25e-27 | **** | 0.652 | 0.544 | 0.108 |
| ran6 | R20:L20 | R20:L5 | 3.285 | 0.001 | 0.001 | ** | 0.652 | 0.624 | 0.028 |
| ran6 | R20:L80 | R20:L5 | -8.85 | 4.77e-16 | 9.54e-16 | **** | 0.544 | 0.624 | -0.08 |
| ran7 | AF | R20:L20 | 4.438 | 1.5e-05 | 1.8e-05 | **** | 0.878 | 0.855 | 0.023 |
| ran7 | AF | R20:L80 | 19.31 | 1.64e-47 | 9.84e-47 | **** | 0.878 | 0.752 | 0.126 |
| ran7 | AF | R20:L5 | 7.154 | 1.58e-11 | 2.37e-11 | **** | 0.878 | 0.842 | 0.036 |
| ran7 | R20:L20 | R20:L80 | 14.46 | 7.11e-33 | 2.13e-32 | **** | 0.855 | 0.752 | 0.103 |
| ran7 | R20:L20 | R20:L5 | 2.584 | 0.011 | 0.011 | * | 0.855 | 0.842 | 0.013 |
| ran7 | R20:L80 | R20:L5 | -13.12 | 9.7e-29 | 1.94e-28 | **** | 0.752 | 0.842 | -0.09 |
| ran8 | AF | R20:L20 | 3.404 | 0.000803 | 0.000964 | *** | 0.689 | 0.658 | 0.031 |
| ran8 | AF | R20:L80 | 12.71 | 1.7e-27 | 1.02e-26 | **** | 0.689 | 0.554 | 0.135 |
| ran8 | AF | R20:L5 | 4.155 | 4.83e-05 | 7.24e-05 | **** | 0.689 | 0.648 | 0.041 |
| ran8 | R20:L20 | R20:L80 | 9.744 | 1.37e-18 | 4.11e-18 | **** | 0.658 | 0.554 | 0.104 |
| ran8 | R20:L20 | R20:L5 | 1.05 | 0.295 | 0.295 | ns | 0.658 | 0.648 | 0.01 |
| ran8 | R20:L80 | R20:L5 | -8.245 | 2.21e-14 | 4.42e-14 | **** | 0.554 | 0.648 | -0.094 |
| ran9 | AF | R20:L20 | 4.827 | 2.76e-06 | 2.76e-06 | **** | 0.905 | 0.883 | 0.022 |
| ran9 | AF | R20:L80 | 21.05 | 1.61e-52 | 9.66e-52 | **** | 0.905 | 0.751 | 0.154 |
| ran9 | AF | R20:L5 | 9.226 | 4.19e-17 | 6.28e-17 | **** | 0.905 | 0.849 | 0.056 |
| ran9 | R20:L20 | R20:L80 | 18.06 | 8.53e-44 | 2.56e-43 | **** | 0.883 | 0.751 | 0.132 |
| ran9 | R20:L20 | R20:L5 | 5.709 | 4.08e-08 | 4.9e-08 | **** | 0.883 | 0.849 | 0.034 |
| ran9 | R20:L80 | R20:L5 | -12.52 | 6.58e-27 | 1.32e-26 | **** | 0.751 | 0.849 | -0.098 |
| snoRNA | AF | R20:L20 | 7.136 | 1.75e-11 | 2.1e-11 | **** | 0.927 | 0.886 | 0.041 |
| snoRNA | AF | R20:L80 | 37.08 | 2.61e-91 | 1.57e-90 | **** | 0.927 | 0.704 | 0.223 |
| snoRNA | AF | R20:L5 | 17.22 | 2.76e-41 | 4.14e-41 | **** | 0.927 | 0.855 | 0.072 |
| snoRNA | R20:L20 | R20:L80 | 23.88 | 2.48e-60 | 4.96e-60 | **** | 0.886 | 0.704 | 0.182 |
| snoRNA | R20:L20 | R20:L5 | 4.96 | 1.51e-06 | 1.51e-06 | **** | 0.886 | 0.855 | 0.031 |
| snoRNA | R20:L80 | R20:L5 | -24.13 | 5.22e-61 | 1.57e-60 | **** | 0.704 | 0.855 | -0.151 |
| tRNAphe | AF | R20:L20 | 9.918 | 4.29e-19 | 5.15e-19 | **** | 0.865 | 0.821 | 0.044 |
| tRNAphe | AF | R20:L80 | 27.5 | 1.07e-69 | 6.42e-69 | **** | 0.865 | 0.704 | 0.161 |
| tRNAphe | AF | R20:L5 | 14.72 | 1.16e-33 | 1.74e-33 | **** | 0.865 | 0.793 | 0.072 |
| tRNAphe | R20:L20 | R20:L80 | 19.3 | 1.75e-47 | 5.25e-47 | **** | 0.821 | 0.704 | 0.117 |
| tRNAphe | R20:L20 | R20:L5 | 5.867 | 1.83e-08 | 1.83e-08 | **** | 0.821 | 0.793 | 0.028 |
| tRNAphe | R20:L80 | R20:L5 | -15.03 | 1.32e-34 | 2.64e-34 | **** | 0.704 | 0.793 | -0.089 |

**Table ST19. Two-way ANOVA for the effect of structure and model on mutational robustness.** Repeated measures under different models.

| Effect | DFn | DFd | $F$ | $p$ | $p < 0.05$ | G. $\eta^2$ |
|---|---|---|---|---|---|---|
| Structure | 14 | 2985 | 656.3 | 0 | * | 0.631 |
| Model | 2.96 | 8826.42 | 220.6 | 4.06e-136 | * | 0.032 |
| Structure:Model | 41.4 | 8826.42 | 4.407 | 2.25e-19 | * | 0.009 |

**Table ST20. One-way ANOVA for the effect of model on mutational robustness.** Repeated measures under different models.

| Structure | DFn | DFd | $F$ | $p$ | $p < 0.05$ | G. $\eta^2$ | Adjusted $p$ |
|---|---|---|---|---|---|---|---|
| CPEB3ribozyme | 2.64 | 524.87 | 12.74 | 2.32e-07 | * | 0.03 | 2.9e-07 |
| DsrA | 1.66 | 330.62 | 17.85 | 3.82e-07 | * | 0.035 | 4.408e-07 |
| HepatitisRibozyme | 3 | 597 | 33.26 | 6.88e-20 | * | 0.099 | 1.032e-18 |
| ran0 | 2.35 | 467.91 | 12.48 | 1.15e-06 | * | 0.043 | 1.232e-06 |
| ran1 | 2.75 | 546.57 | 18.98 | 5.77e-11 | * | 0.029 | 1.082e-10 |
| ran2 | 2.65 | 528.08 | 19.35 | 7.26e-11 | * | 0.057 | 1.21e-10 |
| ran3 | 3 | 597 | 20.73 | 8.72e-13 | * | 0.059 | 2.18e-12 |
| ran4 | 3 | 597 | 21.75 | 2.22e-13 | * | 0.073 | 8.1e-13 |
| ran5 | 2.31 | 459.36 | 11.01 | 6.8e-06 | * | 0.025 | 6.8e-06 |
| ran6 | 3 | 597 | 20.21 | 1.74e-12 | * | 0.041 | 3.729e-12 |
| ran7 | 3 | 597 | 15.56 | 9.29e-10 | * | 0.043 | 1.394e-09 |
| ran8 | 2.64 | 526.3 | 16.23 | 3.25e-09 | * | 0.037 | 4.432e-09 |
| ran9 | 3 | 597 | 23.28 | 2.9e-14 | * | 0.055 | 1.45e-13 |
| snoRNA | 2.31 | 459.14 | 39.59 | 1.53e-18 | * | 0.111 | 1.148e-17 |
| tRNAphe | 2.82 | 561.13 | 22.82 | 2.7e-13 | * | 0.07 | 8.1e-13 |

**Table ST21. Pair-wise post hoc tests for the effect of structure and model on mutational robustness.** Repeated measures under different models.

| Structure | Model M1 | Model M2 | statistic | $p$ | Adjusted $p$ | Signif. | Mean M1 | Mean M2 | Difference |
|---|---|---|---|---|---|---|---|---|---|
| CPEB3 ribozyme | AF | R20:L20 | 0.9965 | 0.32 | 0.48 | ns | 0.512 | 0.508 | 0.004 |
| CPEB3 ribozyme | AF | R20:L80 | 4.85 | 2.48e-06 | 7.44e-06 | **** | 0.512 | 0.488 | 0.024 |
| CPEB3 ribozyme | AF | R20:L5 | 0.299 | 0.765 | 0.765 | ns | 0.512 | 0.511 | 0.001 |
| CPEB3 ribozyme | R20:L20 | R20:L80 | 3.706 | 0.000273 | 0.000546 | *** | 0.508 | 0.488 | 0.02 |
| CPEB3 ribozyme | R20:L20 | R20:L5 | -0.8421 | 0.401 | 0.481 | ns | 0.508 | 0.511 | -0.003 |
| CPEB3 ribozyme | R20:L80 | R20:L5 | -5.189 | 5.19e-07 | 3.11e-06 | **** | 0.488 | 0.511 | -0.023 |
| DsrA | AF | R20:L20 | 3.267 | 0.001 | 0.002 | ** | 0.329 | 0.325 | 0.004 |
| DsrA | AF | R20:L80 | 5.412 | 1.78e-07 | 1.07e-06 | **** | 0.329 | 0.314 | 0.015 |
| DsrA | AF | R20:L5 | 2.708 | 0.007 | 0.009 | ** | 0.329 | 0.325 | 0.004 |
| DsrA | R20:L20 | R20:L80 | 3.899 | 0.000132 | 0.000264 | *** | 0.325 | 0.314 | 0.011 |
| DsrA | R20:L20 | R20:L5 | -0.3791 | 0.705 | 0.705 | ns | 0.325 | 0.325 | 0 |
| DsrA | R20:L80 | R20:L5 | -4.166 | 4.61e-05 | 0.000138 | *** | 0.314 | 0.325 | -0.011 |
| Hepatitis ribozyme | AF | R20:L20 | 2.693 | 0.008 | 0.012 | * | 0.467 | 0.46 | 0.007 |
| Hepatitis ribozyme | AF | R20:L80 | 9.17 | 6.01e-17 | 3.61e-16 | **** | 0.467 | 0.442 | 0.025 |
| Hepatitis ribozyme | AF | R20:L5 | 2.394 | 0.018 | 0.021 | * | 0.467 | 0.461 | 0.006 |
| Hepatitis ribozyme | R20:L20 | R20:L80 | 6.776 | 1.37e-10 | 2.74e-10 | **** | 0.46 | 0.442 | 0.018 |
| Hepatitis ribozyme | R20:L20 | R20:L5 | -0.3538 | 0.724 | 0.724 | ns | 0.46 | 0.461 | -0.001 |
| Hepatitis ribozyme | R20:L80 | R20:L5 | -7.175 | 1.39e-11 | 4.17e-11 | **** | 0.442 | 0.461 | -0.019 |
| ran0 | AF | R20:L20 | -0.3863 | 0.7 | 0.7 | ns | 0.416 | 0.416 | 0 |
| ran0 | AF | R20:L80 | 4.358 | 2.1e-05 | 6.3e-05 | **** | 0.416 | 0.402 | 0.014 |
| ran0 | AF | R20:L5 | 1.046 | 0.297 | 0.356 | ns | 0.416 | 0.413 | 0.003 |
| ran0 | R20:L20 | R20:L80 | 4.733 | 4.19e-06 | 2.51e-05 | **** | 0.416 | 0.402 | 0.014 |
| ran0 | R20:L20 | R20:L5 | 1.5 | 0.135 | 0.202 | ns | 0.416 | 0.413 | 0.003 |
| ran0 | R20:L80 | R20:L5 | -3.505 | 0.000564 | 0.001 | ** | 0.402 | 0.413 | -0.011 |
| ran1 | AF | R20:L20 | 4.438 | 1.5e-05 | 3e-05 | **** | 0.369 | 0.332 | 0.037 |
| ran1 | AF | R20:L80 | 7.113 | 2e-11 | 1.2e-10 | **** | 0.369 | 0.315 | 0.054 |
| ran1 | AF | R20:L5 | 4.834 | 2.67e-06 | 8.01e-06 | **** | 0.369 | 0.328 | 0.041 |
| ran1 | R20:L20 | R20:L80 | 2.7 | 0.008 | 0.011 | * | 0.332 | 0.315 | 0.017 |
| ran1 | R20:L20 | R20:L5 | 0.4977 | 0.619 | 0.619 | ns | 0.332 | 0.328 | 0.004 |
| ran1 | R20:L80 | R20:L5 | -1.83 | 0.069 | 0.083 | ns | 0.315 | 0.328 | -0.013 |
| ran2 | AF | R20:L20 | 0.2634 | 0.793 | 0.793 | ns | 0.492 | 0.491 | 0.001 |
| ran2 | AF | R20:L80 | 5.819 | 2.34e-08 | 7.02e-08 | **** | 0.492 | 0.469 | 0.023 |
| ran2 | AF | R20:L5 | 1.651 | 0.1 | 0.15 | ns | 0.492 | 0.487 | 0.005 |
| ran2 | R20:L20 | R20:L80 | 5.863 | 1.86e-08 | 7.02e-08 | **** | 0.491 | 0.469 | 0.022 |
| ran2 | R20:L20 | R20:L5 | 1.516 | 0.131 | 0.157 | ns | 0.491 | 0.487 | 0.004 |
| ran2 | R20:L80 | R20:L5 | -4.653 | 5.96e-06 | 1.19e-05 | **** | 0.469 | 0.487 | -0.018 |
| ran3 | AF | R20:L20 | -0.9008 | 0.369 | 0.443 | ns | 0.388 | 0.39 | -0.002 |
| ran3 | AF | R20:L80 | 5.617 | 6.47e-08 | 1.29e-07 | **** | 0.388 | 0.374 | 0.014 |
| ran3 | AF | R20:L5 | 0.2051 | 0.838 | 0.838 | ns | 0.388 | 0.387 | 0.001 |
| ran3 | R20:L20 | R20:L80 | 6.779 | 1.34e-10 | 8.04e-10 | **** | 0.39 | 0.374 | 0.016 |
| ran3 | R20:L20 | R20:L5 | 1.191 | 0.235 | 0.352 | ns | 0.39 | 0.387 | 0.003 |
| ran3 | R20:L80 | R20:L5 | -5.965 | 1.1e-08 | 3.3e-08 | **** | 0.374 | 0.387 | -0.013 |

| Structure | Model M1 | Model M2 | statistic | $p$ | Adjusted $p$ | Signif. | Mean M1 | Mean M2 | Difference |
|---|---|---|---|---|---|---|---|---|---|
| ran4 | AF | R20:L20 | -1.252 | 0.212 | 0.212 | ns | 0.486 | 0.49 | -0.004 |
| ran4 | AF | R20:L80 | 6.106 | 5.26e-09 | 1.58e-08 | **** | 0.486 | 0.47 | 0.016 |
| ran4 | AF | R20:L5 | 2.182 | 0.03 | 0.036 | * | 0.486 | 0.481 | 0.005 |
| ran4 | R20:L20 | R20:L80 | 7.941 | 1.44e-13 | 8.64e-13 | **** | 0.49 | 0.47 | 0.02 |
| ran4 | R20:L20 | R20:L5 | 3.415 | 0.000772 | 0.001 | ** | 0.49 | 0.481 | 0.009 |
| ran4 | R20:L80 | R20:L5 | -3.915 | 0.000124 | 0.000248 | *** | 0.47 | 0.481 | -0.011 |
| ran5 | AF | R20:L20 | 0.4478 | 0.655 | 0.786 | ns | 0.442 | 0.44 | 0.002 |
| ran5 | AF | R20:L80 | 4.191 | 4.18e-05 | 8.36e-05 | **** | 0.442 | 0.421 | 0.021 |
| ran5 | AF | R20:L5 | 0.5854 | 0.559 | 0.786 | ns | 0.442 | 0.439 | 0.003 |
| ran5 | R20:L20 | R20:L80 | 4.351 | 2.16e-05 | 6.48e-05 | **** | 0.44 | 0.421 | 0.019 |
| ran5 | R20:L20 | R20:L5 | 0.2723 | 0.786 | 0.786 | ns | 0.44 | 0.439 | 0.001 |
| ran5 | R20:L80 | R20:L5 | -5.718 | 3.91e-08 | 2.35e-07 | **** | 0.421 | 0.439 | -0.018 |
| ran6 | AF | R20:L20 | 1.209 | 0.228 | 0.228 | ns | 0.323 | 0.318 | 0.005 |
| ran6 | AF | R20:L80 | 7.36 | 4.74e-12 | 2.84e-11 | **** | 0.323 | 0.295 | 0.028 |
| ran6 | AF | R20:L5 | 3.167 | 0.002 | 0.003 | ** | 0.323 | 0.311 | 0.012 |
| ran6 | R20:L20 | R20:L80 | 5.869 | 1.8e-08 | 5.4e-08 | **** | 0.318 | 0.295 | 0.023 |
| ran6 | R20:L20 | R20:L5 | 1.814 | 0.071 | 0.085 | ns | 0.318 | 0.311 | 0.007 |
| ran6 | R20:L80 | R20:L5 | -4.007 | 8.68e-05 | 0.000174 | *** | 0.295 | 0.311 | -0.016 |
| ran7 | AF | R20:L20 | 1.231 | 0.22 | 0.22 | ns | 0.423 | 0.42 | 0.003 |
| ran7 | AF | R20:L80 | 6.626 | 3.15e-10 | 1.89e-09 | **** | 0.423 | 0.406 | 0.017 |
| ran7 | AF | R20:L5 | 3.058 | 0.003 | 0.004 | ** | 0.423 | 0.415 | 0.008 |
| ran7 | R20:L20 | R20:L80 | 5.179 | 5.44e-07 | 1.63e-06 | **** | 0.42 | 0.406 | 0.014 |
| ran7 | R20:L20 | R20:L5 | 1.794 | 0.074 | 0.089 | ns | 0.42 | 0.415 | 0.005 |
| ran7 | R20:L80 | R20:L5 | -3.36 | 0.000934 | 0.002 | ** | 0.406 | 0.415 | -0.009 |
| ran8 | AF | R20:L20 | 2.105 | 0.037 | 0.044 | * | 0.402 | 0.392 | 0.01 |
| ran8 | AF | R20:L80 | 6.203 | 3.14e-09 | 1.88e-08 | **** | 0.402 | 0.36 | 0.042 |
| ran8 | AF | R20:L5 | 2.419 | 0.016 | 0.025 | * | 0.402 | 0.389 | 0.013 |
| ran8 | R20:L20 | R20:L80 | 4.47 | 1.31e-05 | 3.93e-05 | **** | 0.392 | 0.36 | 0.032 |
| ran8 | R20:L20 | R20:L5 | 0.5404 | 0.589 | 0.589 | ns | 0.392 | 0.389 | 0.003 |
| ran8 | R20:L80 | R20:L5 | -3.889 | 0.000137 | 0.000274 | *** | 0.36 | 0.389 | -0.029 |
| ran9 | AF | R20:L20 | 0.9383 | 0.349 | 0.377 | ns | 0.409 | 0.407 | 0.002 |
| ran9 | AF | R20:L80 | 7.603 | 1.12e-12 | 6.72e-12 | **** | 0.409 | 0.388 | 0.021 |
| ran9 | AF | R20:L5 | 1.861 | 0.064 | 0.096 | ns | 0.409 | 0.404 | 0.005 |
| ran9 | R20:L20 | R20:L80 | 6.703 | 2.05e-10 | 6.15e-10 | **** | 0.407 | 0.388 | 0.019 |
| ran9 | R20:L20 | R20:L5 | 0.885 | 0.377 | 0.377 | ns | 0.407 | 0.404 | 0.003 |
| ran9 | R20:L80 | R20:L5 | -5.73 | 3.68e-08 | 7.36e-08 | **** | 0.388 | 0.404 | -0.016 |
| snoRNA | AF | R20:L20 | 0.8975 | 0.371 | 0.445 | ns | 0.551 | 0.547 | 0.004 |
| snoRNA | AF | R20:L80 | 11.74 | 1.55e-24 | 9.3e-24 | **** | 0.551 | 0.52 | 0.031 |
| snoRNA | AF | R20:L5 | 1.791 | 0.075 | 0.112 | ns | 0.551 | 0.546 | 0.005 |
| snoRNA | R20:L20 | R20:L80 | 7.148 | 1.63e-11 | 3.26e-11 | **** | 0.547 | 0.52 | 0.027 |
| snoRNA | R20:L20 | R20:L5 | 0.278 | 0.781 | 0.781 | ns | 0.547 | 0.546 | 0.001 |
| snoRNA | R20:L80 | R20:L5 | -10.22 | 5.7e-20 | 1.71e-19 | **** | 0.52 | 0.546 | -0.026 |
| tRNAphe | AF | R20:L20 | -0.8318 | 0.407 | 0.469 | ns | 0.418 | 0.419 | -0.001 |
| tRNAphe | AF | R20:L80 | 6.21 | 3.03e-09 | 9.09e-09 | **** | 0.418 | 0.403 | 0.015 |
| tRNAphe | AF | R20:L5 | 0.725 | 0.469 | 0.469 | ns | 0.418 | 0.416 | 0.002 |
| tRNAphe | R20:L20 | R20:L80 | 6.757 | 1.52e-10 | 9.12e-10 | **** | 0.419 | 0.403 | 0.016 |
| tRNAphe | R20:L20 | R20:L5 | 1.712 | 0.088 | 0.133 | ns | 0.419 | 0.416 | 0.003 |
| tRNAphe | R20:L80 | R20:L5 | -5.643 | 5.7e-08 | 1.14e-07 | **** | 0.403 | 0.416 | -0.013 |

**Table ST22. Two-way ANOVA for the effect of structure and model on plastic repertoire.** Repeated measures under different models.

| Effect | DFn | DFd | $F$ | $p$ | $p < 0.05$ | G. $\eta^2$ |
|---|---|---|---|---|---|---|
| Structure | 14 | 2985 | 90.51 | 5.09e-217 | * | 0.233 |
| Model | 2.4 | 7160.41 | 2161 | 0 | * | 0.172 |
| Structure:Model | 33.58 | 7160.41 | 9.148 | 8.18e-45 | * | 0.012 |

**Table ST23. One-way ANOVA for the effect of model on plastic repertoire.** Repeated measures under different models.

| Structure | DFn | DFd | $F$ | $p$ | $p < 0.05$ | G. $\eta^2$ | Adjusted $p$ |
|---|---|---|---|---|---|---|---|
| CPEB3ribozyme | 2.02 | 402.84 | 210.2 | 9.13e-64 | * | 0.216 | 1.956e-63 |
| DsrA | 2.38 | 474.32 | 157.5 | 2.3e-60 | * | 0.188 | 4.312e-60 |
| HepatitisRibozyme | 1.86 | 369.18 | 297.8 | 3.13e-74 | * | 0.328 | 2.348e-73 |
| ran0 | 2.15 | 428.73 | 175 | 2.68e-59 | * | 0.26 | 4.02e-59 |
| ran1 | 2.63 | 523.19 | 31.71 | 5.54e-17 | * | 0.049 | 5.54e-17 |
| ran2 | 2.62 | 520.72 | 177.6 | 3.6e-72 | * | 0.192 | 1.35e-71 |
| ran3 | 2.14 | 425.79 | 226.8 | 6.79e-71 | * | 0.254 | 2.037e-70 |
| ran4 | 2.1 | 418.3 | 173.3 | 1.66e-57 | * | 0.207 | 2.264e-57 |
| ran5 | 2.48 | 492.87 | 170.9 | 1.58e-66 | * | 0.198 | 3.95e-66 |
| ran6 | 2.46 | 490.02 | 84.97 | 4.34e-38 | * | 0.09 | 5.008e-38 |
| ran7 | 2.06 | 410.22 | 157.4 | 1.43e-52 | * | 0.184 | 1.787e-52 |
| ran8 | 2.33 | 462.79 | 85.62 | 2.36e-36 | * | 0.096 | 2.529e-36 |
| ran9 | 1.99 | 395.11 | 194.9 | 2.53e-59 | * | 0.253 | 4.02e-59 |
| snoRNA | 1.69 | 335.48 | 587.8 | 3.04e-101 | * | 0.568 | 4.56e-100 |
| tRNAphe | 2.03 | 404.09 | 252.9 | 1.16e-72 | * | 0.288 | 5.8e-72 |

**Table ST24. Pair-wise post hoc tests for the effect of structure and model on plastic repertoire.** Repeated measures under different models.

| Structure | Model M1 | Model M2 | statistic | $p$ | Adjusted $p$ | Signif. | Mean M1 | Mean M2 | Difference |
|---|---|---|---|---|---|---|---|---|---|
| CPEB3 ribozyme | AF | R20:L20 | -5.067 | 9.2e-07 | 1.1e-06 | **** | 3.228 | 3.65 | -0.422 |
| CPEB3 ribozyme | AF | R20:L80 | -18.45 | 5.84e-45 | 3.5e-44 | **** | 3.228 | 5.8 | -2.572 |
| CPEB3 ribozyme | AF | R20:L5 | -8.495 | 4.61e-15 | 6.92e-15 | **** | 3.228 | 3.935 | -0.707 |
| CPEB3 ribozyme | R20:L20 | R20:L80 | -16.34 | 1.25e-38 | 3.75e-38 | **** | 3.65 | 5.8 | -2.15 |
| CPEB3 ribozyme | R20:L20 | R20:L5 | -4.005 | 8.76e-05 | 8.76e-05 | **** | 3.65 | 3.935 | -0.285 |
| CPEB3 ribozyme | R20:L80 | R20:L5 | 13.93 | 3.16e-31 | 6.32e-31 | **** | 5.8 | 3.935 | 1.865 |
| DsrA | AF | R20:L20 | -6.405 | 1.06e-09 | 1.27e-09 | **** | 3.563 | 4.307 | -0.744 |
| DsrA | AF | R20:L80 | -18.77 | 6.66e-46 | 4e-45 | **** | 3.563 | 6.442 | -2.879 |
| DsrA | AF | R20:L5 | -9.348 | 1.87e-17 | 2.8e-17 | **** | 3.563 | 4.587 | -1.024 |
| DsrA | R20:L20 | R20:L80 | -12.93 | 3.64e-28 | 1.09e-27 | **** | 4.307 | 6.442 | -2.135 |
| DsrA | R20:L20 | R20:L5 | -2.601 | 0.01 | 0.01 | ** | 4.307 | 4.587 | -0.28 |
| DsrA | R20:L80 | R20:L5 | 11.47 | 1.07e-23 | 2.14e-23 | **** | 6.442 | 4.587 | 1.855 |
| Hepatitis ribozyme | AF | R20:L20 | -9.48 | 7.88e-18 | 9.46e-18 | **** | 3.691 | 4.428 | -0.737 |
| Hepatitis ribozyme | AF | R20:L80 | -21.98 | 3.94e-55 | 2.36e-54 | **** | 3.691 | 7.387 | -3.696 |
| Hepatitis ribozyme | AF | R20:L5 | -10.53 | 6.92e-21 | 1.04e-20 | **** | 3.691 | 4.647 | -0.956 |
| Hepatitis ribozyme | R20:L20 | R20:L80 | -17.98 | 1.44e-43 | 4.32e-43 | **** | 4.428 | 7.387 | -2.959 |
| Hepatitis ribozyme | R20:L20 | R20:L5 | -2.244 | 0.026 | 0.026 | * | 4.428 | 4.647 | -0.219 |
| Hepatitis ribozyme | R20:L80 | R20:L5 | 16.89 | 2.74e-40 | 5.48e-40 | **** | 7.387 | 4.647 | 2.74 |
| ran0 | AF | R20:L20 | -5.619 | 6.41e-08 | 7.69e-08 | **** | 2.079 | 2.512 | -0.433 |
| ran0 | AF | R20:L80 | -18 | 1.25e-43 | 7.5e-43 | **** | 2.079 | 4.437 | -2.358 |
| ran0 | AF | R20:L5 | -10.15 | 8.98e-20 | 1.35e-19 | **** | 2.079 | 2.911 | -0.832 |
| ran0 | R20:L20 | R20:L80 | -14.78 | 7.68e-34 | 2.3e-33 | **** | 2.512 | 4.437 | -1.925 |
| ran0 | R20:L20 | R20:L5 | -4.591 | 7.82e-06 | 7.82e-06 | **** | 2.512 | 2.911 | -0.399 |
| ran0 | R20:L80 | R20:L5 | 11.42 | 1.47e-23 | 2.94e-23 | **** | 4.437 | 2.911 | 1.526 |
| ran1 | AF | R20:L20 | -0.4534 | 0.651 | 0.651 | ns | 6.072 | 6.164 | -0.092 |
| ran1 | AF | R20:L80 | -7.324 | 5.85e-12 | 1.76e-11 | **** | 6.072 | 8.06 | -1.988 |
| ran1 | AF | R20:L5 | -1.835 | 0.068 | 0.096 | ns | 6.072 | 6.487 | -0.415 |
| ran1 | R20:L20 | R20:L80 | -7.35 | 5.03e-12 | 1.76e-11 | **** | 6.164 | 8.06 | -1.896 |
| ran1 | R20:L20 | R20:L5 | -1.762 | 0.08 | 0.096 | ns | 6.164 | 6.487 | -0.323 |
| ran1 | R20:L80 | R20:L5 | 6.471 | 7.41e-10 | 1.48e-09 | **** | 8.06 | 6.487 | 1.573 |
| ran2 | AF | R20:L20 | -4.213 | 3.81e-05 | 4.57e-05 | **** | 2.545 | 2.914 | -0.369 |
| ran2 | AF | R20:L80 | -18.74 | 7.81e-46 | 4.69e-45 | **** | 2.545 | 4.881 | -2.336 |
| ran2 | AF | R20:L5 | -7.47 | 2.47e-12 | 3.7e-12 | **** | 2.545 | 3.274 | -0.729 |
| ran2 | R20:L20 | R20:L80 | -16.31 | 1.61e-38 | 4.83e-38 | **** | 2.914 | 4.881 | -1.967 |
| ran2 | R20:L20 | R20:L5 | -3.764 | 0.00022 | 0.00022 | *** | 2.914 | 3.274 | -0.36 |
| ran2 | R20:L80 | R20:L5 | 13.08 | 1.3e-28 | 2.6e-28 | **** | 4.881 | 3.274 | 1.607 |
| ran3 | AF | R20:L20 | -4.713 | 4.58e-06 | 5.5e-06 | **** | 2.467 | 2.837 | -0.37 |
| ran3 | AF | R20:L80 | -19.78 | 7.1e-49 | 4.26e-48 | **** | 2.467 | 5.171 | -2.704 |
| ran3 | AF | R20:L5 | -8.369 | 1.02e-14 | 1.53e-14 | **** | 2.467 | 3.199 | -0.732 |
| ran3 | R20:L20 | R20:L80 | -17.37 | 9.82e-42 | 2.95e-41 | **** | 2.837 | 5.171 | -2.334 |
| ran3 | R20:L20 | R20:L5 | -4.168 | 4.57e-05 | 4.57e-05 | **** | 2.837 | 3.199 | -0.362 |
| ran3 | R20:L80 | R20:L5 | 14.35 | 1.6e-32 | 3.2e-32 | **** | 5.171 | 3.199 | 1.972 |

| Structure | Model M1 | Model M2 | statistic | $p$ | Adjusted $p$ | Signif. | Mean M1 | Mean M2 | Difference |
|---|---|---|---|---|---|---|---|---|---|
| ran4 | AF | R20:L20 | -6.271 | 2.19e-09 | 2.63e-09 | **** | 2.272 | 2.618 | -0.346 |
| ran4 | AF | R20:L80 | -18.11 | 5.94e-44 | 3.56e-43 | **** | 2.272 | 4.21 | -1.938 |
| ran4 | AF | R20:L5 | -8.418 | 7.49e-15 | 1.12e-14 | **** | 2.272 | 2.897 | -0.625 |
| ran4 | R20:L20 | R20:L80 | -14.55 | 3.75e-33 | 1.13e-32 | **** | 2.618 | 4.21 | -1.592 |
| ran4 | R20:L20 | R20:L5 | -3.885 | 0.000139 | 0.000139 | *** | 2.618 | 2.897 | -0.279 |
| ran4 | R20:L80 | R20:L5 | 11.73 | 1.72e-24 | 3.44e-24 | **** | 4.21 | 2.897 | 1.313 |
| ran5 | AF | R20:L20 | -6.816 | 1.09e-10 | 1.31e-10 | **** | 3.488 | 4.303 | -0.815 |
| ran5 | AF | R20:L80 | -18.82 | 4.68e-46 | 2.81e-45 | **** | 3.488 | 6.552 | -3.064 |
| ran5 | AF | R20:L5 | -9.465 | 8.7e-18 | 1.3e-17 | **** | 3.488 | 4.572 | -1.084 |
| ran5 | R20:L20 | R20:L80 | -13.61 | 2.99e-30 | 8.97e-30 | **** | 4.303 | 6.552 | -2.249 |
| ran5 | R20:L20 | R20:L5 | -2.376 | 0.018 | 0.018 | * | 4.303 | 4.572 | -0.269 |
| ran5 | R20:L80 | R20:L5 | 12.63 | 3.14e-27 | 6.28e-27 | **** | 6.552 | 4.572 | 1.98 |
| ran6 | AF | R20:L20 | -3.122 | 0.002 | 0.002 | ** | 5.728 | 6.302 | -0.574 |
| ran6 | AF | R20:L80 | -12.39 | 1.63e-26 | 9.78e-26 | **** | 5.728 | 9.258 | -3.53 |
| ran6 | AF | R20:L5 | -4.589 | 7.89e-06 | 1.18e-05 | **** | 5.728 | 6.886 | -1.158 |
| ran6 | R20:L20 | R20:L80 | -11.44 | 1.3e-23 | 3.9e-23 | **** | 6.302 | 9.258 | -2.956 |
| ran6 | R20:L20 | R20:L5 | -2.843 | 0.005 | 0.005 | ** | 6.302 | 6.886 | -0.584 |
| ran6 | R20:L80 | R20:L5 | 10.44 | 1.28e-20 | 2.56e-20 | **** | 9.258 | 6.886 | 2.372 |
| ran7 | AF | R20:L20 | -4.796 | 3.17e-06 | 3.8e-06 | **** | 2.6 | 2.984 | -0.384 |
| ran7 | AF | R20:L80 | -17.11 | 6.03e-41 | 3.62e-40 | **** | 2.6 | 4.839 | -2.239 |
| ran7 | AF | R20:L5 | -6.78 | 1.34e-10 | 2.01e-10 | **** | 2.6 | 3.161 | -0.561 |
| ran7 | R20:L20 | R20:L80 | -13.64 | 2.47e-30 | 7.41e-30 | **** | 2.984 | 4.839 | -1.855 |
| ran7 | R20:L20 | R20:L5 | -2.201 | 0.029 | 0.029 | * | 2.984 | 3.161 | -0.177 |
| ran7 | R20:L80 | R20:L5 | 12.05 | 1.8e-25 | 3.6e-25 | **** | 4.839 | 3.161 | 1.678 |
| ran8 | AF | R20:L20 | -2.701 | 0.007 | 0.009 | ** | 5.004 | 5.416 | -0.412 |
| ran8 | AF | R20:L80 | -12.86 | 5.97e-28 | 3.58e-27 | **** | 5.004 | 7.945 | -2.941 |
| ran8 | AF | R20:L5 | -4.384 | 1.89e-05 | 2.84e-05 | **** | 5.004 | 5.69 | -0.686 |
| ran8 | R20:L20 | R20:L80 | -11.49 | 9.01e-24 | 2.7e-23 | **** | 5.416 | 7.945 | -2.529 |
| ran8 | R20:L20 | R20:L5 | -1.548 | 0.123 | 0.123 | ns | 5.416 | 5.69 | -0.274 |
| ran8 | R20:L80 | R20:L5 | 8.934 | 2.78e-16 | 5.56e-16 | **** | 7.945 | 5.69 | 2.255 |
| ran9 | AF | R20:L20 | -5.829 | 2.22e-08 | 2.66e-08 | **** | 2.21 | 2.593 | -0.383 |
| ran9 | AF | R20:L80 | -18.02 | 1.08e-43 | 6.48e-43 | **** | 2.21 | 5.012 | -2.802 |
| ran9 | AF | R20:L5 | -9.527 | 5.78e-18 | 8.67e-18 | **** | 2.21 | 3.151 | -0.941 |
| ran9 | R20:L20 | R20:L80 | -16.1 | 7e-38 | 2.1e-37 | **** | 2.593 | 5.012 | -2.419 |
| ran9 | R20:L20 | R20:L5 | -5.411 | 1.79e-07 | 1.79e-07 | **** | 2.593 | 3.151 | -0.558 |
| ran9 | R20:L80 | R20:L5 | 12.13 | 1.01e-25 | 2.02e-25 | **** | 5.012 | 3.151 | 1.861 |
| snoRNA | AF | R20:L20 | -9.173 | 5.92e-17 | 7.1e-17 | **** | 1.976 | 2.53 | -0.554 |
| snoRNA | AF | R20:L80 | -30.31 | 1.65e-76 | 9.9e-76 | **** | 1.976 | 6.441 | -4.465 |
| snoRNA | AF | R20:L5 | -15.49 | 4.9e-36 | 7.35e-36 | **** | 1.976 | 3.163 | -1.187 |
| snoRNA | R20:L20 | R20:L80 | -26.71 | 9.94e-68 | 2.98e-67 | **** | 2.53 | 6.441 | -3.911 |
| snoRNA | R20:L20 | R20:L5 | -8.163 | 3.68e-14 | 3.68e-14 | **** | 2.53 | 3.163 | -0.633 |
| snoRNA | R20:L80 | R20:L5 | 21.54 | 6.73e-54 | 1.35e-53 | **** | 6.441 | 3.163 | 3.278 |
| tRNAphe | AF | R20:L20 | -7.085 | 2.36e-11 | 2.83e-11 | **** | 2.67 | 3.144 | -0.474 |
| tRNAphe | AF | R20:L80 | -21.36 | 2.14e-53 | 1.28e-52 | **** | 2.67 | 5.268 | -2.598 |
| tRNAphe | AF | R20:L5 | -11.76 | 1.4e-24 | 2.1e-24 | **** | 2.67 | 3.559 | -0.889 |
| tRNAphe | R20:L20 | R20:L80 | -16.7 | 9.99e-40 | 3e-39 | **** | 3.144 | 5.268 | -2.124 |
| tRNAphe | R20:L20 | R20:L5 | -5.501 | 1.15e-07 | 1.15e-07 | **** | 3.144 | 3.559 | -0.415 |
| tRNAphe | R20:L80 | R20:L5 | 14.48 | 6.25e-33 | 1.25e-32 | **** | 5.268 | 3.559 | 1.709 |

**Table ST25. Two-way ANOVA for the effect of structure and model on mutational repertoire.** Repeated measures under different models.

| Effect | DFn | DFd | $F$ | $p$ | $p < 0.05$ | G. $\eta^2$ |
|---|---|---|---|---|---|---|
| Structure | 14 | 2985 | 585.9 | 0 | * | 0.513 |
| Model | 3 | 8955 | 36.11 | 3.46e-23 | * | 0.007 |
| Structure:Model | 42 | 8955 | 1.991 | 0.000149 | * | 0.006 |

**Table ST26. One-way ANOVA for the effect of model on mutational repertoire.** Repeated measures under different models.

| Structure | DFn | DFd | $F$ | $p$ | $p < 0.05$ | G. $\eta^2$ | Adjusted $p$ |
|---|---|---|---|---|---|---|---|
| CPEB3ribozyme | 3 | 597 | 4.466 | 0.004 | * | 0.013 | 0.008571 |
| DsrA | 3 | 597 | 8.77 | 1.07e-05 | * | 0.028 | 0.0001222 |
| HepatitisRibozyme | 2.85 | 567.43 | 4.977 | 0.002 | * | 0.016 | 0.0075 |
| ran0 | 3 | 597 | 2.216 | 0.085 | | 0.008 | 0.1275 |
| ran1 | 3 | 597 | 1.105 | 0.347 | | 0.003 | 0.3718 |
| ran2 | 3 | 597 | 3.013 | 0.03 | * | 0.009 | 0.05 |
| ran3 | 3 | 597 | 3.169 | 0.024 | * | 0.01 | 0.045 |
| ran4 | 3 | 597 | 1.399 | 0.242 | | 0.004 | 0.3025 |
| ran5 | 3 | 597 | 8.465 | 1.63e-05 | * | 0.026 | 0.0001222 |
| ran6 | 3 | 597 | 4.494 | 0.004 | * | 0.012 | 0.008571 |
| ran7 | 3 | 597 | 0.729 | 0.535 | | 0.002 | 0.535 |
| ran8 | 3 | 597 | 4.608 | 0.003 | * | 0.012 | 0.008571 |
| ran9 | 2.89 | 574.3 | 1.423 | 0.236 | | 0.005 | 0.3025 |
| snoRNA | 2.88 | 572.25 | 6.559 | 0.00029 | * | 0.018 | 0.00145 |
| tRNAphe | 2.73 | 543.82 | 1.253 | 0.29 | | 0.004 | 0.3346 |

**Table ST27. Pair-wise post hoc tests for the effect of structure and model on mutational repertoire.** Repeated measures under different models.

| Structure | Model M1 | Model M2 | statistic | $p$ | Adjusted $p$ | Signif. | Mean M1 | Mean M2 | Difference |
|---|---|---|---|---|---|---|---|---|---|
| CPEB3 ribozyme | AF | R20:L20 | 0.09328 | 0.926 | 0.926 | ns | 57.54 | 57.46 | 0.078 |
| CPEB3 ribozyme | AF | R20:L80 | -1.967 | 0.051 | 0.101 | ns | 57.54 | 59.36 | -1.827 |
| CPEB3 ribozyme | AF | R20:L5 | 1.562 | 0.12 | 0.167 | ns | 57.54 | 56.18 | 1.357 |
| CPEB3 ribozyme | R20:L20 | R20:L80 | -2.109 | 0.036 | 0.101 | ns | 57.46 | 59.36 | -1.905 |
| CPEB3 ribozyme | R20:L20 | R20:L5 | 1.486 | 0.139 | 0.167 | ns | 57.46 | 56.18 | 1.279 |
| CPEB3 ribozyme | R20:L80 | R20:L5 | 3.692 | 0.000287 | 0.002 | ** | 59.36 | 56.18 | 3.184 |
| DsrA | AF | R20:L20 | -1.321 | 0.188 | 0.226 | ns | 82.05 | 83.67 | -1.626 |
| DsrA | AF | R20:L80 | -4.984 | 1.35e-06 | 8.1e-06 | **** | 82.05 | 88.39 | -6.343 |
| DsrA | AF | R20:L5 | -2.213 | 0.028 | 0.042 | * | 82.05 | 84.71 | -2.663 |
| DsrA | R20:L20 | R20:L80 | -3.648 | 0.000338 | 0.001 | ** | 83.67 | 88.39 | -4.717 |
| DsrA | R20:L20 | R20:L5 | -0.8021 | 0.423 | 0.423 | ns | 83.67 | 84.71 | -1.037 |
| DsrA | R20:L80 | R20:L5 | 2.617 | 0.01 | 0.019 | * | 88.39 | 84.71 | 3.68 |
| Hepatitis ribozyme | AF | R20:L20 | -2.03 | 0.044 | 0.087 | ns | 84.04 | 87 | -2.959 |
| Hepatitis ribozyme | AF | R20:L80 | -3.952 | 0.000108 | 0.000648 | *** | 84.04 | 88.94 | -4.906 |
| Hepatitis ribozyme | AF | R20:L5 | -0.7781 | 0.437 | 0.437 | ns | 84.04 | 85.07 | -1.034 |
| Hepatitis ribozyme | R20:L20 | R20:L80 | -1.298 | 0.196 | 0.235 | ns | 87 | 88.94 | -1.947 |
| Hepatitis ribozyme | R20:L20 | R20:L5 | 1.32 | 0.188 | 0.235 | ns | 87 | 85.07 | 1.925 |
| Hepatitis ribozyme | R20:L80 | R20:L5 | 3.141 | 0.002 | 0.006 | ** | 88.94 | 85.07 | 3.872 |
| ran0 | AF | R20:L20 | 0.1358 | 0.892 | 0.892 | ns | 55.15 | 55.04 | 0.107 |
| ran0 | AF | R20:L80 | -1.344 | 0.18 | 0.27 | ns | 55.15 | 56.26 | -1.115 |
| ran0 | AF | R20:L5 | -2.108 | 0.036 | 0.116 | ns | 55.15 | 56.9 | -1.754 |
| ran0 | R20:L20 | R20:L80 | -1.445 | 0.15 | 0.27 | ns | 55.04 | 56.26 | -1.222 |
| ran0 | R20:L20 | R20:L5 | -2.081 | 0.039 | 0.116 | ns | 55.04 | 56.9 | -1.861 |
| ran0 | R20:L80 | R20:L5 | -0.6892 | 0.491 | 0.589 | ns | 56.26 | 56.9 | -0.639 |
| ran1 | AF | R20:L20 | 1.016 | 0.311 | 0.622 | ns | 68.75 | 67.85 | 0.901 |
| ran1 | AF | R20:L80 | -0.7333 | 0.464 | 0.696 | ns | 68.75 | 69.44 | -0.683 |
| ran1 | AF | R20:L5 | -0.3312 | 0.741 | 0.741 | ns | 68.75 | 69.07 | -0.315 |
| ran1 | R20:L20 | R20:L80 | -1.807 | 0.072 | 0.433 | ns | 67.85 | 69.44 | -1.584 |
| ran1 | R20:L20 | R20:L5 | -1.321 | 0.188 | 0.564 | ns | 67.85 | 69.07 | -1.216 |
| ran1 | R20:L80 | R20:L5 | 0.4105 | 0.682 | 0.741 | ns | 69.44 | 69.07 | 0.368 |
| ran2 | AF | R20:L20 | -0.9405 | 0.348 | 0.418 | ns | 53.53 | 54.35 | -0.815 |
| ran2 | AF | R20:L80 | -2.819 | 0.005 | 0.032 | * | 53.53 | 56.08 | -2.544 |
| ran2 | AF | R20:L5 | -1.182 | 0.239 | 0.358 | ns | 53.53 | 54.56 | -1.03 |
| ran2 | R20:L20 | R20:L80 | -1.951 | 0.052 | 0.158 | ns | 54.35 | 56.08 | -1.729 |
| ran2 | R20:L20 | R20:L5 | -0.2739 | 0.784 | 0.784 | ns | 54.35 | 54.56 | -0.215 |
| ran2 | R20:L80 | R20:L5 | 1.741 | 0.083 | 0.167 | ns | 56.08 | 54.56 | 1.514 |
| ran3 | AF | R20:L20 | 2.137 | 0.034 | 0.101 | ns | 61.2 | 59.38 | 1.826 |
| ran3 | AF | R20:L80 | -1.003 | 0.317 | 0.38 | ns | 61.2 | 62.15 | -0.944 |
| ran3 | AF | R20:L5 | 0.4565 | 0.649 | 0.649 | ns | 61.2 | 60.79 | 0.41 |
| ran3 | R20:L20 | R20:L80 | -2.915 | 0.004 | 0.024 | * | 59.38 | 62.15 | -2.77 |
| ran3 | R20:L20 | R20:L5 | -1.579 | 0.116 | 0.228 | ns | 59.38 | 60.79 | -1.416 |
| ran3 | R20:L80 | R20:L5 | 1.438 | 0.152 | 0.228 | ns | 62.15 | 60.79 | 1.354 |

| Structure | Model M1 | Model M2 | statistic | $p$ | Adjusted $p$ | Signif. | Mean M1 | Mean M2 | Difference |
|---|---|---|---|---|---|---|---|---|---|
| ran4 | AF | R20:L20 | 1.464 | 0.145 | 0.435 | ns | 50.55 | 49.35 | 1.203 |
| ran4 | AF | R20:L80 | -0.5042 | 0.615 | 0.615 | ns | 50.55 | 50.97 | -0.414 |
| ran4 | AF | R20:L5 | 0.6153 | 0.539 | 0.615 | ns | 50.55 | 50.04 | 0.518 |
| ran4 | R20:L20 | R20:L80 | -1.98 | 0.049 | 0.295 | ns | 49.35 | 50.97 | -1.617 |
| ran4 | R20:L20 | R20:L5 | -0.781 | 0.436 | 0.615 | ns | 49.35 | 50.04 | -0.685 |
| ran4 | R20:L80 | R20:L5 | 1.13 | 0.26 | 0.52 | ns | 50.97 | 50.04 | 0.932 |
| ran5 | AF | R20:L20 | -0.6847 | 0.494 | 0.494 | ns | 57.28 | 57.89 | -0.61 |
| ran5 | AF | R20:L80 | -3.829 | 0.000172 | 0.000516 | *** | 57.28 | 60.62 | -3.343 |
| ran5 | AF | R20:L5 | 0.813 | 0.417 | 0.494 | ns | 57.28 | 56.61 | 0.675 |
| ran5 | R20:L20 | R20:L80 | -3.044 | 0.003 | 0.005 | ** | 57.89 | 60.62 | -2.733 |
| ran5 | R20:L20 | R20:L5 | 1.577 | 0.116 | 0.174 | ns | 57.89 | 56.61 | 1.285 |
| ran5 | R20:L80 | R20:L5 | 4.853 | 2.45e-06 | 1.47e-05 | **** | 60.62 | 56.61 | 4.018 |
| ran6 | AF | R20:L20 | 0.4571 | 0.648 | 0.648 | ns | 74.18 | 73.74 | 0.441 |
| ran6 | AF | R20:L80 | -2.765 | 0.006 | 0.019 | * | 74.18 | 77.21 | -3.025 |
| ran6 | AF | R20:L5 | -0.8219 | 0.412 | 0.494 | ns | 74.18 | 75.07 | -0.883 |
| ran6 | R20:L20 | R20:L80 | -3.536 | 0.000506 | 0.003 | ** | 73.74 | 77.21 | -3.466 |
| ran6 | R20:L20 | R20:L5 | -1.347 | 0.179 | 0.268 | ns | 73.74 | 75.07 | -1.324 |
| ran6 | R20:L80 | R20:L5 | 2.014 | 0.045 | 0.091 | ns | 77.21 | 75.07 | 2.142 |
| ran7 | AF | R20:L20 | -0.4543 | 0.65 | 0.733 | ns | 63.82 | 64.24 | -0.423 |
| ran7 | AF | R20:L80 | -1.31 | 0.192 | 0.706 | ns | 63.82 | 65.06 | -1.238 |
| ran7 | AF | R20:L5 | -1.049 | 0.295 | 0.706 | ns | 63.82 | 64.74 | -0.923 |
| ran7 | R20:L20 | R20:L80 | -0.9313 | 0.353 | 0.706 | ns | 64.24 | 65.06 | -0.815 |
| ran7 | R20:L20 | R20:L5 | -0.5733 | 0.567 | 0.733 | ns | 64.24 | 64.74 | -0.5 |
| ran7 | R20:L80 | R20:L5 | 0.3421 | 0.733 | 0.733 | ns | 65.06 | 64.74 | 0.315 |
| ran8 | AF | R20:L20 | -1.774 | 0.078 | 0.116 | ns | 62.11 | 63.68 | -1.576 |
| ran8 | AF | R20:L80 | -3.457 | 0.000667 | 0.004 | ** | 62.11 | 65.34 | -3.229 |
| ran8 | AF | R20:L5 | -1.346 | 0.18 | 0.216 | ns | 62.11 | 63.23 | -1.119 |
| ran8 | R20:L20 | R20:L80 | -1.874 | 0.062 | 0.116 | ns | 63.68 | 65.34 | -1.653 |
| ran8 | R20:L20 | R20:L5 | 0.5127 | 0.609 | 0.609 | ns | 63.68 | 63.23 | 0.457 |
| ran8 | R20:L80 | R20:L5 | 2.432 | 0.016 | 0.048 | * | 65.34 | 63.23 | 2.11 |
| ran9 | AF | R20:L20 | 1.305 | 0.193 | 0.502 | ns | 64.43 | 63.34 | 1.09 |
| ran9 | AF | R20:L80 | -0.7173 | 0.474 | 0.569 | ns | 64.43 | 65.02 | -0.592 |
| ran9 | AF | R20:L5 | 0.06253 | 0.95 | 0.95 | ns | 64.43 | 64.38 | 0.054 |
| ran9 | R20:L20 | R20:L80 | -2.303 | 0.022 | 0.134 | ns | 63.34 | 65.02 | -1.682 |
| ran9 | R20:L20 | R20:L5 | -1.15 | 0.251 | 0.502 | ns | 63.34 | 64.38 | -1.036 |
| ran9 | R20:L80 | R20:L5 | 0.8033 | 0.423 | 0.569 | ns | 65.02 | 64.38 | 0.646 |
| snoRNA | AF | R20:L20 | 1.714 | 0.088 | 0.106 | ns | 87.9 | 85.29 | 2.608 |
| snoRNA | AF | R20:L80 | -1.798 | 0.074 | 0.106 | ns | 87.9 | 90.42 | -2.522 |
| snoRNA | AF | R20:L5 | 2.112 | 0.036 | 0.072 | ns | 87.9 | 85.1 | 2.798 |
| snoRNA | R20:L20 | R20:L80 | -3.482 | 0.000612 | 0.002 | ** | 85.29 | 90.42 | -5.13 |
| snoRNA | R20:L20 | R20:L5 | 0.1387 | 0.89 | 0.89 | ns | 85.29 | 85.1 | 0.19 |
| snoRNA | R20:L80 | R20:L5 | 4.421 | 1.61e-05 | 9.66e-05 | **** | 90.42 | 85.1 | 5.32 |
| tRNAphe | AF | R20:L20 | 0.6568 | 0.512 | 0.768 | ns | 72.84 | 72.12 | 0.724 |
| tRNAphe | AF | R20:L80 | -1.283 | 0.201 | 0.402 | ns | 72.84 | 74.19 | -1.346 |
| tRNAphe | AF | R20:L5 | 0.365 | 0.716 | 0.842 | ns | 72.84 | 72.35 | 0.49 |
| tRNAphe | R20:L20 | R20:L80 | -2.048 | 0.042 | 0.251 | ns | 72.12 | 74.19 | -2.07 |
| tRNAphe | R20:L20 | R20:L5 | -0.199 | 0.842 | 0.842 | ns | 72.12 | 72.35 | -0.234 |
| tRNAphe | R20:L80 | R20:L5 | 1.413 | 0.159 | 0.402 | ns | 74.19 | 72.35 | 1.836 |

**Table ST28. Two-way ANOVA for the effect of structure and model on population's plastic repertoire.** Repeated measures under different models.

| Effect | DFn | DFd | $F$ | $p$ | $p < 0.05$ | G. $\eta^2$ |
|---|---|---|---|---|---|---|
| Structure | 14 | 2985 | 73.97 | 4.04e-181 | * | 0.132 |
| Model | 2.76 | 8240.04 | 1001 | 0 | * | 0.159 |
| Structure:Model | 38.65 | 8240.04 | 6.391 | 1.3e-31 | * | 0.017 |

**Table ST29. One-way ANOVA for the effect of model on population's plastic repertoire.** Repeated measures under different models.

| Structure | DFn | DFd | $F$ | $p$ | $p < 0.05$ | G. $\eta^2$ | Adjusted $p$ |
|---|---|---|---|---|---|---|---|
| CPEB3ribozyme | 2.45 | 486.98 | 69.92 | 4.02e-32 | * | 0.17 | 5.482e-32 |
| DsrA | 2.41 | 478.87 | 136.9 | 9.94e-55 | * | 0.303 | 1.491e-53 |
| HepatitisRibozyme | 2.85 | 566.97 | 59 | 7.16e-32 | * | 0.157 | 8.95e-32 |
| ran0 | 2.42 | 482.18 | 129.6 | 8.59e-53 | * | 0.284 | 6.442e-52 |
| ran1 | 2.63 | 522.69 | 76.37 | 5.86e-37 | * | 0.158 | 9.767e-37 |
| ran2 | 2.46 | 489.81 | 61.74 | 5.12e-29 | * | 0.162 | 5.486e-29 |
| ran3 | 2.18 | 433.78 | 102.7 | 1e-39 | * | 0.213 | 2.143e-39 |
| ran4 | 2.46 | 490.27 | 93.11 | 4.15e-41 | * | 0.207 | 1.245e-40 |
| ran5 | 2.73 | 543.64 | 80.62 | 3.92e-40 | * | 0.199 | 9.8e-40 |
| ran6 | 2.83 | 562.3 | 104.3 | 2.66e-51 | * | 0.195 | 1.33e-50 |
| ran7 | 2.7 | 537.94 | 67.83 | 2.73e-34 | * | 0.145 | 4.095e-34 |
| ran8 | 2.71 | 538.44 | 78.62 | 6.12e-39 | * | 0.16 | 1.148e-38 |
| ran9 | 2.66 | 528.82 | 62.74 | 1.49e-31 | * | 0.15 | 1.719e-31 |
| snoRNA | 2.58 | 512.61 | 48.74 | 1.43e-24 | * | 0.121 | 1.43e-24 |
| tRNAphe | 2.83 | 563.72 | 83.49 | 9.31e-43 | * | 0.206 | 3.491e-42 |

**Table ST30. Pair-wise post hoc tests for the effect of structure and model on population's plastic repertoire.** Repeated measures under different models.

| Structure | Model M1 | Model M2 | statistic | $p$ | Adjusted $p$ | Signif. | Mean M1 | Mean M2 | Difference |
|---|---|---|---|---|---|---|---|---|---|
| CPEB3 ribozyme | AF | R20:L20 | -5.356 | 2.34e-07 | 2.81e-07 | **** | 49.8 | 71.96 | -22.16 |
| CPEB3 ribozyme | AF | R20:L80 | -12.28 | 3.66e-26 | 2.2e-25 | **** | 49.8 | 121.2 | -71.4 |
| CPEB3 ribozyme | AF | R20:L5 | -7.125 | 1.86e-11 | 2.79e-11 | **** | 49.8 | 77.01 | -27.21 |
| CPEB3 ribozyme | R20:L20 | R20:L80 | -8.178 | 3.36e-14 | 1.01e-13 | **** | 71.96 | 121.2 | -49.24 |
| CPEB3 ribozyme | R20:L20 | R20:L5 | -1.12 | 0.264 | 0.264 | ns | 71.96 | 77.01 | -5.05 |
| CPEB3 ribozyme | R20:L80 | R20:L5 | 7.886 | 2.02e-13 | 4.04e-13 | **** | 121.2 | 77.01 | 44.18 |
| DsrA | AF | R20:L20 | -8.404 | 8.15e-15 | 9.78e-15 | **** | 49.66 | 83.55 | -33.88 |
| DsrA | AF | R20:L80 | -18 | 1.28e-43 | 7.68e-43 | **** | 49.66 | 155.5 | -105.8 |
| DsrA | AF | R20:L5 | -9.248 | 3.62e-17 | 5.43e-17 | **** | 49.66 | 90.48 | -40.82 |
| DsrA | R20:L20 | R20:L80 | -11.09 | 1.45e-22 | 4.35e-22 | **** | 83.55 | 155.5 | -71.96 |
| DsrA | R20:L20 | R20:L5 | -1.501 | 0.135 | 0.135 | ns | 83.55 | 90.48 | -6.935 |
| DsrA | R20:L80 | R20:L5 | 10.6 | 4.3e-21 | 8.6e-21 | **** | 155.5 | 90.48 | 65.03 |
| Hepatitis ribozyme | AF | R20:L20 | -5.704 | 4.19e-08 | 5.03e-08 | **** | 81.94 | 128.2 | -46.3 |
| Hepatitis ribozyme | AF | R20:L80 | -12.6 | 3.89e-27 | 2.33e-26 | **** | 81.94 | 209.5 | -127.5 |
| Hepatitis ribozyme | AF | R20:L5 | -5.904 | 1.51e-08 | 2.26e-08 | **** | 81.94 | 137.8 | -55.81 |
| Hepatitis ribozyme | R20:L20 | R20:L80 | -8.084 | 6.01e-14 | 1.8e-13 | **** | 128.2 | 209.5 | -81.21 |
| Hepatitis ribozyme | R20:L20 | R20:L5 | -0.9915 | 0.323 | 0.323 | ns | 128.2 | 137.8 | -9.505 |
| Hepatitis ribozyme | R20:L80 | R20:L5 | 6.699 | 2.1e-10 | 4.2e-10 | **** | 209.5 | 137.8 | 71.71 |
| ran0 | AF | R20:L20 | -7.678 | 7.13e-13 | 8.56e-13 | **** | 28.55 | 47.97 | -19.42 |
| ran0 | AF | R20:L80 | -17.13 | 5.1e-41 | 3.06e-40 | **** | 28.55 | 93.28 | -64.73 |
| ran0 | AF | R20:L5 | -11.65 | 2.95e-24 | 5.9e-24 | **** | 28.55 | 58.64 | -30.09 |
| ran0 | R20:L20 | R20:L80 | -11.87 | 6.63e-25 | 1.99e-24 | **** | 47.97 | 93.28 | -45.31 |
| ran0 | R20:L20 | R20:L5 | -3.351 | 0.000962 | 0.000962 | *** | 47.97 | 58.64 | -10.67 |
| ran0 | R20:L80 | R20:L5 | 8.626 | 2e-15 | 3e-15 | **** | 93.28 | 58.64 | 34.64 |
| ran1 | AF | R20:L20 | -4.941 | 1.64e-06 | 1.97e-06 | **** | 67.19 | 91.19 | -24 |
| ran1 | AF | R20:L80 | -13.48 | 7.4e-30 | 4.44e-29 | **** | 67.19 | 153.6 | -86.4 |
| ran1 | AF | R20:L5 | -6.044 | 7.29e-09 | 1.09e-08 | **** | 67.19 | 98.66 | -31.47 |
| ran1 | R20:L20 | R20:L80 | -9.861 | 6.27e-19 | 1.88e-18 | **** | 91.19 | 153.6 | -62.4 |
| ran1 | R20:L20 | R20:L5 | -1.406 | 0.161 | 0.161 | ns | 91.19 | 98.66 | -7.47 |
| ran1 | R20:L80 | R20:L5 | 7.809 | 3.23e-13 | 6.46e-13 | **** | 153.6 | 98.66 | 54.94 |
| ran2 | AF | R20:L20 | -4.794 | 3.19e-06 | 3.83e-06 | **** | 37.91 | 56.24 | -18.33 |
| ran2 | AF | R20:L80 | -11.63 | 3.45e-24 | 2.07e-23 | **** | 37.91 | 100.5 | -62.6 |
| ran2 | AF | R20:L5 | -6.851 | 8.96e-11 | 1.61e-10 | **** | 37.91 | 63.62 | -25.71 |
| ran2 | R20:L20 | R20:L80 | -7.962 | 1.27e-13 | 3.81e-13 | **** | 56.24 | 100.5 | -44.28 |
| ran2 | R20:L20 | R20:L5 | -1.82 | 0.07 | 0.07 | ns | 56.24 | 63.62 | -7.38 |
| ran2 | R20:L80 | R20:L5 | 6.82 | 1.07e-10 | 1.61e-10 | **** | 100.5 | 63.62 | 36.9 |
| ran3 | AF | R20:L20 | -5.882 | 1.69e-08 | 2.03e-08 | **** | 42.18 | 60.46 | -18.29 |
| ran3 | AF | R20:L80 | -13.85 | 5.38e-31 | 3.23e-30 | **** | 42.18 | 112.7 | -70.51 |
| ran3 | AF | R20:L5 | -7.152 | 1.6e-11 | 2.4e-11 | **** | 42.18 | 66.17 | -23.99 |
| ran3 | R20:L20 | R20:L80 | -11.55 | 5.9e-24 | 1.77e-23 | **** | 60.46 | 112.7 | -52.22 |
| ran3 | R20:L20 | R20:L5 | -1.785 | 0.076 | 0.076 | ns | 60.46 | 66.17 | -5.705 |
| ran3 | R20:L80 | R20:L5 | 8.828 | 5.52e-16 | 1.1e-15 | **** | 112.7 | 66.17 | 46.52 |

| Structure | Model M1 | Model M2 | statistic | $p$ | Adjusted $p$ | Signif. | Mean M1 | Mean M2 | Difference |
|---|---|---|---|---|---|---|---|---|---|
| ran4 | AF | R20:L20 | -6.252 | 2.43e-09 | 2.92e-09 | **** | 32.25 | 48.04 | -15.79 |
| ran4 | AF | R20:L80 | -13.77 | 9.84e-31 | 5.9e-30 | **** | 32.25 | 81.18 | -48.94 |
| ran4 | AF | R20:L5 | -7.111 | 2.02e-11 | 3.03e-11 | **** | 32.25 | 48.95 | -16.7 |
| ran4 | R20:L20 | R20:L80 | -9.535 | 5.47e-18 | 1.09e-17 | **** | 48.04 | 81.18 | -33.15 |
| ran4 | R20:L20 | R20:L5 | -0.3563 | 0.722 | 0.722 | ns | 48.04 | 48.95 | -0.91 |
| ran4 | R20:L80 | R20:L5 | 9.621 | 3.11e-18 | 9.33e-18 | **** | 81.18 | 48.95 | 32.24 |
| ran5 | AF | R20:L20 | -6.583 | 4e-10 | 6e-10 | **** | 42.78 | 68.8 | -26.02 |
| ran5 | AF | R20:L80 | -13.9 | 3.94e-31 | 2.36e-30 | **** | 42.78 | 113.1 | -70.36 |
| ran5 | AF | R20:L5 | -6.356 | 1.39e-09 | 1.67e-09 | **** | 42.78 | 67.86 | -25.08 |
| ran5 | R20:L20 | R20:L80 | -8.564 | 2.98e-15 | 5.96e-15 | **** | 68.8 | 113.1 | -44.34 |
| ran5 | R20:L20 | R20:L5 | 0.2113 | 0.833 | 0.833 | ns | 68.8 | 67.86 | 0.935 |
| ran5 | R20:L80 | R20:L5 | 9.203 | 4.85e-17 | 1.46e-16 | **** | 113.1 | 67.86 | 45.28 |
| ran6 | AF | R20:L20 | -6.255 | 2.39e-09 | 2.87e-09 | **** | 73.57 | 107.7 | -34.16 |
| ran6 | AF | R20:L80 | -15.57 | 2.83e-36 | 1.7e-35 | **** | 73.57 | 175.5 | -101.9 |
| ran6 | AF | R20:L5 | -9.132 | 7.71e-17 | 1.54e-16 | **** | 73.57 | 122.8 | -49.24 |
| ran6 | R20:L20 | R20:L80 | -11 | 2.66e-22 | 7.98e-22 | **** | 107.7 | 175.5 | -67.75 |
| ran6 | R20:L20 | R20:L5 | -2.893 | 0.004 | 0.004 | ** | 107.7 | 122.8 | -15.08 |
| ran6 | R20:L80 | R20:L5 | 8.326 | 1.33e-14 | 2e-14 | **** | 175.5 | 122.8 | 52.67 |
| ran7 | AF | R20:L20 | -5.241 | 4.05e-07 | 4.86e-07 | **** | 51.5 | 72.58 | -21.08 |
| ran7 | AF | R20:L80 | -12.58 | 4.33e-27 | 2.6e-26 | **** | 51.5 | 115.9 | -64.38 |
| ran7 | AF | R20:L5 | -6.179 | 3.58e-09 | 5.37e-09 | **** | 51.5 | 75.4 | -23.89 |
| ran7 | R20:L20 | R20:L80 | -8.436 | 6.66e-15 | 2e-14 | **** | 72.58 | 115.9 | -43.3 |
| ran7 | R20:L20 | R20:L5 | -0.6582 | 0.511 | 0.511 | ns | 72.58 | 75.4 | -2.815 |
| ran7 | R20:L80 | R20:L5 | 7.934 | 1.51e-13 | 3.02e-13 | **** | 115.9 | 75.4 | 40.48 |
| ran8 | AF | R20:L20 | -6.201 | 3.19e-09 | 4.78e-09 | **** | 58.9 | 84.28 | -25.38 |
| ran8 | AF | R20:L80 | -13.54 | 4.99e-30 | 2.99e-29 | **** | 58.9 | 131.5 | -72.63 |
| ran8 | AF | R20:L5 | -6.021 | 8.23e-09 | 9.88e-09 | **** | 58.9 | 84.23 | -25.33 |
| ran8 | R20:L20 | R20:L80 | -8.868 | 4.25e-16 | 1.28e-15 | **** | 84.28 | 131.5 | -47.26 |
| ran8 | R20:L20 | R20:L5 | 0.01015 | 0.992 | 0.992 | ns | 84.28 | 84.23 | 0.045 |
| ran8 | R20:L80 | R20:L5 | 8.78 | 7.52e-16 | 1.5e-15 | **** | 131.5 | 84.23 | 47.3 |
| ran9 | AF | R20:L20 | -4.056 | 7.16e-05 | 8.59e-05 | **** | 53.34 | 70.84 | -17.5 |
| ran9 | AF | R20:L80 | -11.5 | 8.2e-24 | 4.92e-23 | **** | 53.34 | 117.2 | -63.9 |
| ran9 | AF | R20:L5 | -6.195 | 3.29e-09 | 4.94e-09 | **** | 53.34 | 77.42 | -24.08 |
| ran9 | R20:L20 | R20:L80 | -8.758 | 8.66e-16 | 2.6e-15 | **** | 70.84 | 117.2 | -46.4 |
| ran9 | R20:L20 | R20:L5 | -1.479 | 0.141 | 0.141 | ns | 70.84 | 77.42 | -6.585 |
| ran9 | R20:L80 | R20:L5 | 7.661 | 7.9e-13 | 1.58e-12 | **** | 117.2 | 77.42 | 39.81 |
| snoRNA | AF | R20:L20 | -2.535 | 0.012 | 0.014 | * | 95.39 | 123.1 | -27.75 |
| snoRNA | AF | R20:L80 | -9.199 | 4.98e-17 | 2.15e-16 | **** | 95.39 | 224.2 | -128.8 |
| snoRNA | AF | R20:L5 | -3.158 | 0.002 | 0.003 | ** | 95.39 | 132.7 | -37.34 |
| snoRNA | R20:L20 | R20:L80 | -9.143 | 7.17e-17 | 2.15e-16 | **** | 123.1 | 224.2 | -101 |
| snoRNA | R20:L20 | R20:L5 | -1.082 | 0.281 | 0.281 | ns | 123.1 | 132.7 | -9.59 |
| snoRNA | R20:L80 | R20:L5 | 8.666 | 1.55e-15 | 3.1e-15 | **** | 224.2 | 132.7 | 91.43 |
| tRNAphe | AF | R20:L20 | -4.858 | 2.4e-06 | 2.88e-06 | **** | 48.2 | 70.9 | -22.7 |
| tRNAphe | AF | R20:L80 | -14.91 | 3.1e-34 | 1.86e-33 | **** | 48.2 | 129.9 | -81.73 |
| tRNAphe | AF | R20:L5 | -7.379 | 4.24e-12 | 6.36e-12 | **** | 48.2 | 83.64 | -35.44 |
| tRNAphe | R20:L20 | R20:L80 | -9.976 | 2.9e-19 | 8.7e-19 | **** | 70.9 | 129.9 | -59.03 |
| tRNAphe | R20:L20 | R20:L5 | -2.425 | 0.016 | 0.016 | * | 70.9 | 83.64 | -12.74 |
| tRNAphe | R20:L80 | R20:L5 | 8.059 | 7e-14 | 1.4e-13 | **** | 129.9 | 83.64 | 46.29 |

**Table ST31. Two-way ANOVA for the effect of structure and model on population's mutational repertoire.** Repeated measures under different models.

| Effect | DFn | DFd | $F$ | $p$ | $p < 0.05$ | G. $\eta^2$ |
|---|---|---|---|---|---|---|
| Structure | 14 | 2985 | 482.3 | 0 | * | 0.514 |
| Model | 2.8 | 8350.89 | 2463 | 0 | * | 0.306 |
| Structure:Model | 39.17 | 8350.89 | 27.77 | 2.19e-189 | * | 0.065 |

**Table ST32. One-way ANOVA for the effect of model on population's mutational repertoire.** Repeated measures under different models.

| Structure | DFn | DFd | $F$ | $p$ | $p < 0.05$ | G. $\eta^2$ | Adjusted $p$ |
|---|---|---|---|---|---|---|---|
| CPEB3ribozyme | 2.69 | 535.37 | 159.4 | 2.29e-68 | * | 0.286 | 2.862e-68 |
| DsrA | 2.73 | 542.98 | 271.1 | 3.04e-101 | * | 0.451 | 4.56e-100 |
| HepatitisRibozyme | 2.86 | 569.99 | 169.9 | 3.76e-76 | * | 0.317 | 6.267e-76 |
| ran0 | 2.74 | 544.33 | 194.9 | 1.38e-80 | * | 0.366 | 5.175e-80 |
| ran1 | 2.68 | 532.35 | 199 | 4.48e-80 | * | 0.321 | 1.344e-79 |
| ran2 | 2.57 | 511.54 | 159.1 | 2.4e-65 | * | 0.323 | 2.769e-65 |
| ran3 | 2.62 | 521 | 136 | 5.5e-59 | * | 0.273 | 5.5e-59 |
| ran4 | 2.83 | 563.5 | 137.6 | 3.95e-64 | * | 0.273 | 4.232e-64 |
| ran5 | 2.76 | 549.9 | 179.6 | 1.15e-76 | * | 0.339 | 2.464e-76 |
| ran6 | 2.65 | 526.37 | 279.2 | 3.52e-100 | * | 0.392 | 2.64e-99 |
| ran7 | 2.85 | 567.22 | 171.4 | 2.75e-76 | * | 0.321 | 5.156e-76 |
| ran8 | 3 | 597 | 179.1 | 8.45e-83 | * | 0.298 | 4.225e-82 |
| ran9 | 2.72 | 541.96 | 158.5 | 7.16e-69 | * | 0.286 | 9.764e-69 |
| snoRNA | 2.71 | 540.01 | 180.7 | 1.12e-75 | * | 0.317 | 1.68e-75 |
| tRNAphe | 2.86 | 569.61 | 181.3 | 7.57e-80 | * | 0.321 | 1.892e-79 |

**Table ST33. Pair-wise post hoc tests for the effect of structure and model on population's mutational repertoire.** Repeated measures under different models.

| Structure | Model M1 | Model M2 | statistic | $p$ | Adjusted $p$ | Signif. | Mean M1 | Mean M2 | Difference |
|---|---|---|---|---|---|---|---|---|---|
| CPEB3 ribozyme | AF | R20:L20 | -11.93 | 4.29e-25 | 8.58e-25 | **** | 307.9 | 429.9 | -122 |
| CPEB3 ribozyme | AF | R20:L80 | -20.38 | 1.33e-50 | 7.98e-50 | **** | 307.9 | 568.1 | -260.2 |
| CPEB3 ribozyme | AF | R20:L5 | -14.15 | 6.45e-32 | 1.94e-31 | **** | 307.9 | 446.8 | -138.9 |
| CPEB3 ribozyme | R20:L20 | R20:L80 | -10.11 | 1.16e-19 | 1.74e-19 | **** | 429.9 | 568.1 | -138.2 |
| CPEB3 ribozyme | R20:L20 | R20:L5 | -1.477 | 0.141 | 0.141 | ns | 429.9 | 446.8 | -16.93 |
| CPEB3 ribozyme | R20:L80 | R20:L5 | 9.279 | 2.95e-17 | 3.54e-17 | **** | 568.1 | 446.8 | 121.3 |
| DsrA | AF | R20:L20 | -14.26 | 2.97e-32 | 5.94e-32 | **** | 480.1 | 735.7 | -255.7 |
| DsrA | AF | R20:L80 | -27.42 | 1.6e-69 | 9.6e-69 | **** | 480.1 | 1061 | -581.4 |
| DsrA | AF | R20:L5 | -16.66 | 1.32e-39 | 3.96e-39 | **** | 480.1 | 769.9 | -289.8 |
| DsrA | R20:L20 | R20:L80 | -13.85 | 5.34e-31 | 8.01e-31 | **** | 735.7 | 1061 | -325.8 |
| DsrA | R20:L20 | R20:L5 | -1.767 | 0.079 | 0.079 | ns | 735.7 | 769.9 | -34.15 |
| DsrA | R20:L80 | R20:L5 | 12.95 | 3.22e-28 | 3.86e-28 | **** | 1061 | 769.9 | 291.6 |
| Hepatitis ribozyme | AF | R20:L20 | -11.41 | 1.59e-23 | 2.38e-23 | **** | 597.3 | 834.6 | -237.3 |
| Hepatitis ribozyme | AF | R20:L80 | -21.32 | 2.88e-53 | 1.73e-52 | **** | 597.3 | 1108 | -510.6 |
| Hepatitis ribozyme | AF | R20:L5 | -11.85 | 7.19e-25 | 1.88e-24 | **** | 597.3 | 847.6 | -250.3 |
| Hepatitis ribozyme | R20:L20 | R20:L80 | -11.82 | 9.38e-25 | 1.88e-24 | **** | 834.6 | 1108 | -273.4 |
| Hepatitis ribozyme | R20:L20 | R20:L5 | -0.6151 | 0.539 | 0.539 | ns | 834.6 | 847.6 | -13.02 |
| Hepatitis ribozyme | R20:L80 | R20:L5 | 10.3 | 3.28e-20 | 3.94e-20 | **** | 1108 | 847.6 | 260.3 |
| ran0 | AF | R20:L20 | -12.72 | 1.66e-27 | 3.32e-27 | **** | 259.7 | 375.2 | -115.5 |
| ran0 | AF | R20:L80 | -23.16 | 2.11e-58 | 1.27e-57 | **** | 259.7 | 516.2 | -256.5 |
| ran0 | AF | R20:L5 | -16.69 | 1.13e-39 | 3.39e-39 | **** | 259.7 | 414.3 | -154.6 |
| ran0 | R20:L20 | R20:L80 | -11.71 | 1.9e-24 | 2.85e-24 | **** | 375.2 | 516.2 | -141 |
| ran0 | R20:L20 | R20:L5 | -3.731 | 0.000249 | 0.000249 | *** | 375.2 | 414.3 | -39.12 |
| ran0 | R20:L80 | R20:L5 | 8.429 | 6.97e-15 | 8.36e-15 | **** | 516.2 | 414.3 | 101.9 |
| ran1 | AF | R20:L20 | -11.89 | 5.48e-25 | 8.22e-25 | **** | 323.8 | 428.6 | -104.7 |
| ran1 | AF | R20:L80 | -22.56 | 9.71e-57 | 5.83e-56 | **** | 323.8 | 589 | -265.2 |
| ran1 | AF | R20:L5 | -13.15 | 7.78e-29 | 1.56e-28 | **** | 323.8 | 449.8 | -126 |
| ran1 | R20:L20 | R20:L80 | -13.43 | 1.1e-29 | 3.3e-29 | **** | 428.6 | 589 | -160.4 |
| ran1 | R20:L20 | R20:L5 | -2.05 | 0.042 | 0.042 | * | 428.6 | 449.8 | -21.25 |
| ran1 | R20:L80 | R20:L5 | 11.02 | 2.43e-22 | 2.92e-22 | **** | 589 | 449.8 | 139.2 |
| ran2 | AF | R20:L20 | -12.35 | 2.21e-26 | 4.42e-26 | **** | 270.9 | 382.8 | -111.8 |
| ran2 | AF | R20:L80 | -20.24 | 3.31e-50 | 1.99e-49 | **** | 270.9 | 514.4 | -243.5 |
| ran2 | AF | R20:L5 | -14.59 | 2.98e-33 | 8.94e-33 | **** | 270.9 | 402.2 | -131.3 |
| ran2 | R20:L20 | R20:L80 | -9.971 | 3e-19 | 4.5e-19 | **** | 382.8 | 514.4 | -131.6 |
| ran2 | R20:L20 | R20:L5 | -1.835 | 0.068 | 0.068 | ns | 382.8 | 402.2 | -19.44 |
| ran2 | R20:L80 | R20:L5 | 8.988 | 1.97e-16 | 2.36e-16 | **** | 514.4 | 402.2 | 112.2 |
| ran3 | AF | R20:L20 | -10.92 | 4.86e-22 | 9.72e-22 | **** | 315.8 | 415.9 | -100.1 |
| ran3 | AF | R20:L80 | -20.06 | 1.09e-49 | 6.54e-49 | **** | 315.8 | 547.1 | -231.2 |
| ran3 | AF | R20:L5 | -11.73 | 1.67e-24 | 5.01e-24 | **** | 315.8 | 438.5 | -122.6 |
| ran3 | R20:L20 | R20:L80 | -9.854 | 6.56e-19 | 9.84e-19 | **** | 415.9 | 547.1 | -131.2 |
| ran3 | R20:L20 | R20:L5 | -2.117 | 0.035 | 0.035 | * | 415.9 | 438.5 | -22.55 |
| ran3 | R20:L80 | R20:L5 | 8.154 | 3.89e-14 | 4.67e-14 | **** | 547.1 | 438.5 | 108.6 |

| Structure | Model M1 | Model M2 | statistic | $p$ | Adjusted $p$ | Signif. | Mean M1 | Mean M2 | Difference |
|---|---|---|---|---|---|---|---|---|---|
| ran4 | AF | R20:L20 | -7.779 | 3.88e-13 | 4.66e-13 | **** | 289 | 363.2 | -74.15 |
| ran4 | AF | R20:L80 | -18.17 | 3.84e-44 | 2.3e-43 | **** | 289 | 483 | -194 |
| ran4 | AF | R20:L5 | -11.18 | 7.81e-23 | 1.56e-22 | **** | 289 | 381.5 | -92.44 |
| ran4 | R20:L20 | R20:L80 | -12.3 | 3.08e-26 | 9.24e-26 | **** | 363.2 | 483 | -119.8 |
| ran4 | R20:L20 | R20:L5 | -1.94 | 0.054 | 0.054 | ns | 363.2 | 381.5 | -18.29 |
| ran4 | R20:L80 | R20:L5 | 10.14 | 9.79e-20 | 1.47e-19 | **** | 483 | 381.5 | 101.5 |
| ran5 | AF | R20:L20 | -10.74 | 1.67e-21 | 2e-21 | **** | 294.8 | 394.8 | -100 |
| ran5 | AF | R20:L80 | -22.29 | 5.48e-56 | 3.29e-55 | **** | 294.8 | 542.7 | -247.9 |
| ran5 | AF | R20:L5 | -11.74 | 1.57e-24 | 3.14e-24 | **** | 294.8 | 407.5 | -112.7 |
| ran5 | R20:L20 | R20:L80 | -12.46 | 1.05e-26 | 3.15e-26 | **** | 394.8 | 542.7 | -147.9 |
| ran5 | R20:L20 | R20:L5 | -1.268 | 0.206 | 0.206 | ns | 394.8 | 407.5 | -12.73 |
| ran5 | R20:L80 | R20:L5 | 11.09 | 1.45e-22 | 2.18e-22 | **** | 542.7 | 407.5 | 135.2 |
| ran6 | AF | R20:L20 | -15.42 | 8.41e-36 | 1.68e-35 | **** | 325.9 | 474.7 | -148.8 |
| ran6 | AF | R20:L80 | -25.61 | 6.54e-65 | 3.92e-64 | **** | 325.9 | 651.1 | -325.2 |
| ran6 | AF | R20:L5 | -17.2 | 3.11e-41 | 9.33e-41 | **** | 325.9 | 497.6 | -171.7 |
| ran6 | R20:L20 | R20:L80 | -14.32 | 2e-32 | 3e-32 | **** | 474.7 | 651.1 | -176.5 |
| ran6 | R20:L20 | R20:L5 | -2.412 | 0.017 | 0.017 | * | 474.7 | 497.6 | -22.95 |
| ran6 | R20:L80 | R20:L5 | 11.95 | 3.69e-25 | 4.43e-25 | **** | 651.1 | 497.6 | 153.5 |
| ran7 | AF | R20:L20 | -10.58 | 4.83e-21 | 7.24e-21 | **** | 331.9 | 430.5 | -98.53 |
| ran7 | AF | R20:L80 | -21.98 | 4.04e-55 | 2.42e-54 | **** | 331.9 | 577.6 | -245.7 |
| ran7 | AF | R20:L5 | -12.95 | 3.22e-28 | 9.66e-28 | **** | 331.9 | 464.6 | -132.6 |
| ran7 | R20:L20 | R20:L80 | -12.61 | 3.51e-27 | 7.02e-27 | **** | 430.5 | 577.6 | -147.2 |
| ran7 | R20:L20 | R20:L5 | -3.072 | 0.002 | 0.002 | ** | 430.5 | 464.6 | -34.09 |
| ran7 | R20:L80 | R20:L5 | 9.444 | 1e-17 | 1.2e-17 | **** | 577.6 | 464.6 | 113.1 |
| ran8 | AF | R20:L20 | -11.67 | 2.6e-24 | 5.2e-24 | **** | 311 | 418.6 | -107.6 |
| ran8 | AF | R20:L80 | -21.9 | 6.51e-55 | 3.91e-54 | **** | 311 | 547.1 | -236 |
| ran8 | AF | R20:L5 | -11.33 | 2.83e-23 | 3.4e-23 | **** | 311 | 420.1 | -109 |
| ran8 | R20:L20 | R20:L80 | -11.6 | 4.23e-24 | 6.34e-24 | **** | 418.6 | 547.1 | -128.4 |
| ran8 | R20:L20 | R20:L5 | -0.1455 | 0.884 | 0.884 | ns | 418.6 | 420.1 | -1.415 |
| ran8 | R20:L80 | R20:L5 | 11.93 | 4.29e-25 | 1.29e-24 | **** | 547.1 | 420.1 | 127 |
| ran9 | AF | R20:L20 | -10.95 | 3.76e-22 | 5.64e-22 | **** | 321.8 | 426 | -104.2 |
| ran9 | AF | R20:L80 | -20.79 | 8.7e-52 | 5.22e-51 | **** | 321.8 | 556.3 | -234.5 |
| ran9 | AF | R20:L5 | -15.43 | 7.79e-36 | 2.34e-35 | **** | 321.8 | 458.7 | -136.9 |
| ran9 | R20:L20 | R20:L80 | -11.28 | 3.95e-23 | 7.9e-23 | **** | 426 | 556.3 | -130.3 |
| ran9 | R20:L20 | R20:L5 | -2.901 | 0.004 | 0.004 | ** | 426 | 458.7 | -32.68 |
| ran9 | R20:L80 | R20:L5 | 7.952 | 1.35e-13 | 1.62e-13 | **** | 556.3 | 458.7 | 97.64 |
| snoRNA | AF | R20:L20 | -10.28 | 3.67e-20 | 4.4e-20 | **** | 668.7 | 901.7 | -233 |
| snoRNA | AF | R20:L80 | -21.98 | 3.87e-55 | 2.32e-54 | **** | 668.7 | 1243 | -573.9 |
| snoRNA | AF | R20:L5 | -13.88 | 4.29e-31 | 1.29e-30 | **** | 668.7 | 946.3 | -277.6 |
| snoRNA | R20:L20 | R20:L80 | -11.72 | 1.84e-24 | 3.68e-24 | **** | 901.7 | 1243 | -340.9 |
| snoRNA | R20:L20 | R20:L5 | -1.819 | 0.07 | 0.07 | ns | 901.7 | 946.3 | -44.67 |
| snoRNA | R20:L80 | R20:L5 | 11.66 | 2.77e-24 | 4.16e-24 | **** | 1243 | 946.3 | 296.2 |
| tRNAphe | AF | R20:L20 | -10.68 | 2.37e-21 | 3.56e-21 | **** | 442.6 | 592.4 | -149.9 |
| tRNAphe | AF | R20:L80 | -22.75 | 2.91e-57 | 1.75e-56 | **** | 442.6 | 796.2 | -353.6 |
| tRNAphe | AF | R20:L5 | -12.82 | 8.15e-28 | 2.44e-27 | **** | 442.6 | 623.4 | -180.8 |
| tRNAphe | R20:L20 | R20:L80 | -12.14 | 9.89e-26 | 1.98e-25 | **** | 592.4 | 796.2 | -203.7 |
| tRNAphe | R20:L20 | R20:L5 | -2.18 | 0.03 | 0.03 | * | 592.4 | 623.4 | -30.97 |
| tRNAphe | R20:L80 | R20:L5 | 10.47 | 9.97e-21 | 1.2e-20 | **** | 796.2 | 623.4 | 172.8 |
